# ANO6 Scaffolds ER-Mitochondria Contacts to Govern Spatial Lipid Compartmentalization and Ferroptosis Evasion in Chemoresistant Breast Cancer

**DOI:** 10.64898/2026.10.22.761200

**Authors:** Qianlu Yang, Jie Yan, Ping Zhao, Qi Tang, Kai Zheng, Jiawen Zhao, Ziting Zhou, Qianxi Yang

**Affiliations:** Department of breast surgery of The Third Affiliated Hospital of Kunming Medical University,Yunnan Cancer Hospital, Peking University Cancer Hospital Yunnan, 519 Kunzhou Road, Xishan District, Kunming 650500, China; Yunnan Eye Institute & Key Laboratory of Yunnan Province, Yunnan Eye Disease Clinical Medical Center, Affiliated Hospital of Yunnan University, Yunnan University, 176 Qingnian, Kunming 650021, China

## Abstract

**Background:** Metabolic glycerophospholipid remodeling promotes tumor persistence under neoadjuvant chemotherapy in breast cancer; however, how resistant cells evade lipotoxicity from accumulating peroxidizable lipids presents an intriguing metabolic conundrum. The underlying organelle-level spatial mechanisms remain poorly defined.

**Methods:** Integrated multi-omic profiling of paired pre- and post-NAC clinical specimens was performed. Engineered ANO6 models (wild-type, knockout, and conformational mutants) were characterized using subcellular lipidomics of purified mitochondria-associated ER membranes (MAMs), electron microscopy, in situ proximity ligation assays, and preclinical xenograft models. Results: Chemoresistant lesions exhibited marked ANO6 upregulation and polyunsaturated phospholipid (PUFA-PL) accumulation without enhanced apoptosis. Beyond its canonical role as a plasma membrane scramblase, ANO6 operates as an organelle contact scaffold at MAMs, leveraging its conformational activation to coordinate inter-organellar tethering. ANO6 directly recruits vimentin (VIM) and VAPB to assemble a VIM-ANO6-VAPB-PTPIP51 tethering platform, structurally bridging intermediate filaments to organelle contact sites. This complex tightens ER-mitochondria coupling, establishing a specialized structural platform that routes and sequesters peroxidizable lipids into protective cytoplasmic lipid droplets, thereby shielding functional membrane bilayers from ferroptotic peroxidation. Functional dissection revealed that the active scaffolding conformation (I612A) sustains MAMs integrity and lipid packaging, whereas scaffold deficiency (D703R) compromises contact architecture. Pharmacologically perturbing this platform with niclosamide destabilized lipid sequestration, selectively resensitizing chemoresistant cells to synthetic lethal ferroptosis in vitro and in vivo.

**Conclusions:** ANO6-mediated contact scaffolding governs spatial lipid compartmentalization, establishing the ANO6-MAMs structural axis as a potential target to exploit lipid metabolic vulnerabilities in refractory breast cancer.

**Statement of significance:** ANO6 scaffolds organelle contacts independent of scramblase activity to govern spatial lipid compartmentalization, providing a structural framework to exploit metabolic vulnerabilities and sensitize chemoresistant breast cancer to ferroptosis.

**Graphical Abstract:** 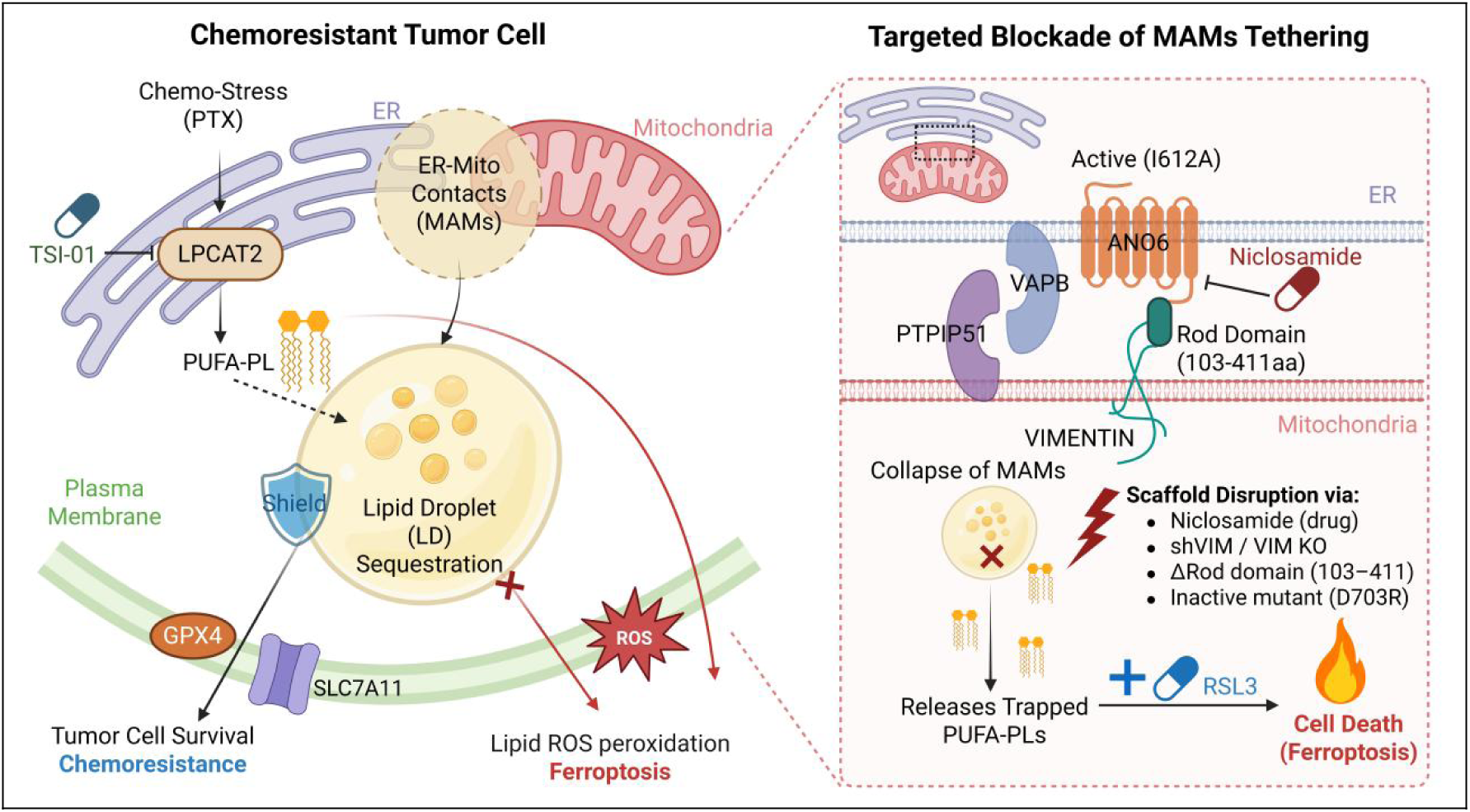

Chemoresistant tumor cells exploit an ANO6-mediated MAMs platform (VIM-ANO6-VAPB-PTPIP51) to coordinate lipid droplet packaging and shield lethal PUFA-PLs from membrane peroxidation. Pharmacological or genetic disruption of this architectural tether collapses the sequestration compartment, liberating trapped lipids to trigger synthetic lethal ferroptosis alongside GPX4 inhibition (RSL3).

## Introduction

Neoadjuvant chemotherapy (NAC) represents a cornerstone of treatment for locally advanced breast cancer; however, approximately 40% of patients fail to achieve a pathological complete response and eventually experience chemoresistant relapse^[1]^. Surviving residual tumor cells actively rewire their metabolic networks to adapt to persistent cytotoxic insults^[2]^. Among diverse metabolic adaptations, aberrant glycerophospholipid remodeling has emerged as a hallmark of therapy persistence^[2–4]^. Exposure to conventional cytotoxic regimens, including taxanes and anthracyclines, provokes profound perturbations in lipid homeostasis and triggers the accumulation of polyunsaturated fatty acid-containing phospholipids (PUFA-PLs)^[5]^. While excessive accumulation of peroxidizable PUFA-PLs theoretically exposes cancer cells to lethal oxidative membrane damage such as ferroptosis^[5]^, chemoresistant clones successfully evade these cytotoxic consequences^[6]^. The precise subcellular strategies by which resistant cells manage and spatially compartmentalize this altered lipid landscape to buffer against lipotoxicity remain poorly understood^[3, 7]^.

Intracellular lipid homeostasis is fundamentally governed by spatial compartmentalization, relying not solely on enzymatic flux but critically on precise organellar distribution^[8]^. Mitochondria-associated endoplasmic reticulum membranes (MAMs) serve as specialized membrane contact sites coordinating phospholipid biosynthesis, inter-organellar trafficking, and calcium signaling^[8, 9]^. MAMs are dynamically enriched with key lipid-modifying enzymes and facilitate the reciprocal exchange of phospholipid precursors between organelles^[10]^. Emerging evidence reveals that MAMs also act as an architectural cradle for lipid droplet (LD) biogenesis, where specialized lipid-transfer machineries (e.g., ORP5 and ORP8) functionalize at the ER-mitochondria interface to coordinate with key LD assembly factors (such as Seipin) to facilitate neutral lipid packaging^[11, 12]^. However, whether MAMs function as an active structural depot to spatially route and partition pro-oxidative lipid species into protective lipid droplets during chemotherapy resistance remains unexplored. Given that MAMs ultrastructure dynamically reorganizes in response to pharmacological stress^[13]^, delineating their structural contribution to spatial lipid compartmentalization and subsequent lipid droplet sequestration is crucial for understanding metabolic persistence in cancer.

Anoctamin 6 (ANO6, also known as TMEM16F) emerges as an intriguing candidate linking inter-organellar contact architecture with adaptive lipid remodeling^[14]^. ANO6 is canonically characterized as a calcium-activated phospholipid scramblase that collapses membrane lipid asymmetry to mediate the externalization of phosphatidylserine (PS) during apoptosis and platelet coagulation^[15]^. Paradoxically, ANO6 is markedly upregulated in aggressive and therapy-refractory malignancies that exhibit negligible apoptotic cell death^[16]^. Consistent with this clinical phenomenon, our initial multi-omic profiling of paired pre- and post-chemotherapy biopsies revealed that residual chemoresistant lesions display pronounced ANO6 induction and altered phospholipid profiles without concurrent apoptotic execution. This clinical and phenotypic disconnect strongly implies that ANO6 exerts an apoptosis-independent, non-canonical function in therapy-resistant states. Given that TMEM16 proteins possess multifaceted structural plasticity that accommodates non-transport cellular functions^[17, 18]^, we hypothesized that the conformational transitions of ANO6 fundamentally dictate not only its canonical lipid scrambling, but also an unrecognized structural scaffolding role at membrane contact sites to coordinate the spatial sheltering of pro-ferroptotic lipids.

In this study, we investigated the non-canonical scaffolding role of ANO6 in governing organelle contact architecture and spatial lipid homeostasis in breast cancer. We demonstrate that ANO6, independent of its lipid scramblase activity, directly recruits vimentin (VIM) and VAPB to assemble a VIM-ANO6-VAPB-PTPIP51 tethering platform at MAMs, effectively bridging the intermediate filament cytoskeleton to inter-organellar contact sites. This structural complex tightens ER-mitochondria contacts, establishing a specialized structural platform that routes and sequesters peroxidizable PUFA-PLs into protective cytoplasmic lipid droplets, thereby shielding functional membrane bilayers from oxidative ferroptotic damage. Utilizing conformational mutants, we demonstrate that the active scaffolding conformation (I612A) sustains MAMs integrity and lipid packaging, whereas scaffold deficiency (D703R) collapses contact architecture and restores metabolic vulnerability. Pharmacologically perturbing this platform with niclosamide dismantles the sequestration compartment, releasing trapped PUFA-PLs and selectively exposing chemoresistant cells to synthetic lethal ferroptotic killing in vitro and in vivo. Collectively, these findings establish organelle-mediated lipid compartmentalization as a key driver of therapy persistence, identifying the ANO6-MAMs structural axis as an actionable target to exploit lipid metabolic vulnerabilities in refractory breast cancer.

## Materials and Methods

### Clinical Specimens and Ethical Approval

Paired primary breast cancer specimens, comprising pre-treatment diagnostic core needle biopsies (Pre-NAC) and matched post-treatment chemoresistant residual surgical lesions (Post-NAC), were obtained from the Third Affiliated Hospital of Kunming Medical University. The study protocol was formally approved by the Institutional Ethics Committee of the Third Affiliated Hospital of Kunming Medical University (Approval No. KYCS2023-056). All enrolled cases were histopathologically diagnosed with primary breast cancer, and written informed consent was acquired from all participants prior to tissue collection. This study adhered strictly to the Declaration of Helsinki, the ethical guidelines of the Council for International Organizations of Medical Sciences (CIOMS), and relevant international ethical frameworks. Harvested tissues were immediately snap-frozen in liquid nitrogen and stored at −80°C for multi-omics profiling or embedded in paraffin and OCT compound for histopathological and multiplex immunofluorescence analyses. Proteomic and metabolomic profiling of clinical tissues was performed by Sinotech Genomics (Shanghai, China). The clinicopathological characteristics of the multi-omics cohort (n = 6 pairs), luminal IHC cohort (n = 35 pairs), and TNBC IHC cohort (n = 25 pairs) are summarized in Supplementary Table S1A, S1B, and S1C, respectively.

### Immunohistochemistry (IHC) Staining and Automated Digital H-Score Quantification

Immunohistochemistry (IHC) Staining and Automated Digital H-Score QuantificationFormalin-fixed paraffin-embedded (FFPE) sections (4 μm) from paired breast cancer tissues were deparaffinized, rehydrated, and subjected to high-pressure antigen retrieval in citrate buffer (pH 6.0). Endogenous peroxidase was quenched with 3% H_2_O_2_, and non-specific binding was blocked with 5% normal goat serum. Sections were incubated overnight at 4°C with primary anti-ANO6 antibody (ThermoFisher, Cat# PA5-58610, 1:200), followed by HRP-conjugated secondary antibodies and DAB visualization. Images were captured under a light microscope (scale bar: 100 μm). ANO6 staining was quantitatively evaluated using an automated ImageJ/Fiji macro with the “H DAB” color deconvolution vector. Staining intensity was stratified by 8-bit grayscale thresholds (0-255): background (>235), weak (121-180), moderate (61-120), and strong (<60). Digital H-scores were calculated as: *H* − *score* = (1 × *p_weak_*) + (2 ×*P_moderate_*) + (3 × *P_strong_*) , Where Pi represents the percentage of stained tumor area at each intensity.

### Cell Culture and Quality Control

Mouse mammary carcinoma EMT6, human breast cancer MDA-MB-231, colorectal adenocarcinoma CACO2, hepatocellular carcinoma HepG2, and non-small cell lung carcinoma A549 cell lines were maintained in RPMI 1640 medium (VivaCell Biosciences, Cat# C3010-0500) supplemented with 10% fetal bovine serum (FBS) and 1% penicillin-streptomycin at 37°C with 5% CO_2_. Human cell lines were authenticated by short tandem repeat (STR) profiling and utilized within 10 passages post-thawing. Cultures were regularly verified to be free of mycoplasma contamination using a Mycoplasma PCR Detection Kit (Beyotime Biotechnology, Cat# C0301S).

### Generation of Engineered Cell Lines

Engineered breast cancer cell models encompassing stable overexpression, CRISPR-Cas9 knockout, conformational mutant reconstitution, and intermediate filament disruption were established in mouse EMT6 and human MDA-MB-231 lines.

Ano6 Overexpression and Knockout: For Ano6 overexpression, mouse EMT6 and human MDA-MB-231 cells were transduced with lentiviruses encoding murine Ano6 (GV721 backbone, puromycin resistance) or human ANO6 (GV492-luciferase backbone, puromycin resistance), with CON547 and CON335 serving as respective negative controls, followed by selection with puromycin. For CRISPR-Cas9-mediated gene disruption, EMT6 cells were transduced with lentiviral vectors (GV708 backbone, puromycin resistance) expressing three sgRNAs targeting murine Ano6 (C25J0047, C25J0048, C25J0049) or a non-targeting control sgRNA (CON544), while MDA-MB-231 cells were transduced with GV708 expressing human ANO6 sgRNAs (C26A1538, C26A1539, C26A1540). Following puromycin selection, single-cell clones were isolated via limiting dilution and validated by immunoblotting.

Conformational Mutant Reconstitution: To genetically uncouple scaffolding architecture from scramblase catalytic activity, full-length murine Ano6 cDNAs encoding the active scaffolding mutant (I612A-HA) or the scaffold-compromised mutant (D703R-HA) were synthesized de novo. To prevent Cas9 endonuclease re-cleavage of the rescue constructs, synonymous mutations were engineered within the sgRNA target sites. Additionally, a C-terminal HA tag followed by a stop codon was introduced to avoid expression of the vector-encoded 3FLAG. Both I612A-HA and D703R-HA constructs were cloned into the GV870 lentiviral vector (neomycin resistance). Ano6-deficient EMT6 cells were stably transduced with I612A-HA or D703R-HA lentiviruses alongside the matched parental empty vector (CON709) to generate control pairs (I612A vs. I612A-EV and D703R vs. D703R-EV). Stable pools were selected with G418/neomycin, and equivalent protein reconstitution was verified by immunoblotting against HA and Ano6.

Intermediate Filament Disruption and Vim Mutants: For Vim knockdown, lentiviruses expressing three independent shRNAs targeting murine Vim (GV116 backbone, hygromycin resistance; P26F1261, P26F1262, P26F1263) or a non-targeting control (CON482) were transduced into EMT6 ANO6-OE and I612A-HA rescue cells, followed by hygromycin selection. To map the Ano6-Vim interaction via reciprocal binding, full-length murine Vim (Vim-HA) and a truncation mutant lacking the central rod/pocket domain [Vim(Δ103-411aa)-HA] were cloned into the GV366 vector (neomycin resistance). Because stable truncation mutants of Ano6 could not be generated, this Vim mutant was used to validate the interaction.

Transient Transfection and CoIP: Transient transfection of the GV366 plasmids and small interfering RNAs targeting Vimentin (siVim: 50 nM; GenePharma) was performed using Lipofectamine 3000 (Thermo Fisher Scientific, Cat# L3000015) in Opti-MEM according to the manufacturer’s instructions. For CoIP, cells were lysed 48-72 h post-transfection, and immunoprecipitation was performed using anti-HA or anti-Ano6 antibodies, followed by western blotting to assess the Ano6-Vim interaction. Detailed sequence information for vectors, sgRNAs, and primers is provided in Supplementary Table S2.

### Live-Cell ER-Mitochondria Co-localization

Cells seeded on glass coverslips were subjected to indicated treatments (such as 5 nM paclitaxel for 24 h). Following treatment, cells were incubated with 1:1,000 diluted ER-Tracker Green (Beyotime, Cat# C1042) and 200 nM Mito-Tracker Red CMXRos (Beyotime, Cat# C1049B) in pre-warmed complete medium at 37°C in the dark for 30 min. Nuclei were counterstained with 1 μg/mL Hoechst 33342 (Servicebio, G1127-1ML) for 15 min. Confocal fluorescent images were acquired using a Leica TCS SP5 laser scanning confocal microscope equipped with a 100× oil-immersion objective in sequential scanning mode. Spatial co-localization and signal alignment between the ER and mitochondria were quantitatively assessed across cross-sectional transects using the Plot Profile tool in ImageJ/Fiji software.

### Transmission Electron Microscopy (TEM) and Morphometric Analysis

Harvested cells were initially fixed in 0.1 M sodium cacodylate buffer (pH 7.4) containing 2.5% glutaraldehyde and 1.5% potassium ferrocyanide at 4°C overnight, followed by post-fixation with 1% osmium tetroxide (OsO_4_) for 1.5 h at room temperature. Samples were dehydrated through a graded acetone series (50%-100%), infiltrated, and embedded in Epon 812 epoxy resin. Ultrathin sections (70 nm) were prepared using an ultramicrotome, collected on copper grids, and double-stained with 2% uranyl acetate and lead citrate. Ultrastructural imaging was performed using a Hitachi HT-7800 transmission electron microscope operated at 80 kV. For intracellular lipid droplet (LD) quantification, cytoplasmic LDs were segmented manually, and total LD burden was calculated as the percentage of cumulative LD cross-sectional area relative to total cytoplasmic area. For mitochondria-associated ER membrane (MAM) morphometrics, close appositions (≤ 60 nm cleft distance, a standard operational distance encompassing both calcium-signaling junctions and wider lipid-transfer microdomains^[8]^) between the outer mitochondrial membrane and smooth/rough ER cisternae were traced. The ER-mitochondria contact coefficient (ERMICC) was defined and computed as the cumulative contact length divided by total mitochondrial perimeter (*ERMICC* = ∑*_contact_* /P*_mito_*) .

### In Situ Proximity Ligation Assay (In Situ PLA)

Protein-protein spatial interactions were detected using the Duolink In Situ Red Starter Kit (Sigma-Aldrich, Cat# DUO92101) according to the manufacturer’s instructions. Briefly, cells cultured on chamber slides were fixed with ice-cold methanol at −20°C for 10 min and blocked with Duolink Blocking Solution at 37°C for 1 h. Samples were incubated overnight at 4°C with validated primary antibody pairs from different host species: rabbit anti-ANO6 (Thermo Fisher Scientific, Cat# PA5-58610, 1:200) paired with mouse anti-Vimentin (Proteintech, Cat# 60330-1-Ig, 1:500); rabbit anti-ANO6 paired with mouse anti-VAPB (Proteintech, Cat# 66191-1-Ig, 1:200); or mouse anti-VAPB paired with rabbit anti-PTPIP51 (Proteintech, Cat# 20641-1-AP, 1:200). After washing with Wash Buffer A, preparations were incubated with species-specific secondary PLA probes (anti-rabbit PLUS and anti-mouse MINUS) at 37°C for 1 h. Oligonucleotides were hybridized, ligated, and amplified by rolling circle amplification (RCA) using fluorescently labeled nucleotides (detection excitation/emission = 594/613 nm). Coverslips were mounted with Duolink In Situ Mounting Medium containing DAPI. Confocal z-stack micrographs were acquired under a laser scanning confocal microscope (100× oil objective), and distinct interaction puncta per cell (representing physical proximity < 40 nm) were enumerated across > 50 randomly selected cells per condition using the Find Maxima function in ImageJ/Fiji.

### Membrane Lipid Packaging and DiI Labeling

Following indicated treatments (vehicle, 5 nM PTX, PTX plus 50 μM arachidonic acid (AA; Sigma-Aldrich, Cat# A3555), or PTX plus 10 μM TSI-01(LPCAT2 inhibitor; MedChemExpress, Cat# HY-112921)), cells grown on coverslips were incubated with the lipophilic fluorescent membrane probe DiI (BBcellProbe, Cat# BB-441914; diluted 1:1,000 in serum-free medium) at 37°C in the dark for 1 h. Cells were washed three times with pre-warmed PBS, mildly fixed with 4% paraformaldehyde (PFA) for 15 min at room temperature, and counterstained with DAPI (1 μg/mL) for 10 min. Fluorescent signals reflecting cellular lipid uptake, internalization, and packaging were acquired using a fluorescence/confocal microscope (excitation/emission = 549/565 nm) under identical exposure settings across cohorts. The extent of lipid internalization was quantified from at least five random fields per group using ImageJ/Fiji as the integrated fluorescence density normalized to the number of DAPI-positive nuclei (% DiI-positive signal per cell).

### Subcellular Fractionation and Purification of MAMs

Mitochondria-associated endoplasmic reticulum membrane (MAM) fractions were isolated from engineered breast cancer cells following chemotherapy challenge (5 nM paclitaxel for 24 h) using a density-gradient ultracentrifugation isolation system (Servicebio, Cat# swsMAM02) with optimization. Briefly, cells harvested at logarithmic growth were washed twice with ice-cold Ca^2+^/Mg^2+^-free PBS and gently disrupted in pre-chilled TB1 homogenization buffer using a Dounce homogenizer. Homogenates were centrifuged at 600g for 5 min at 4°C to pellet intact cells and nuclei. The post-nuclear supernatant was collected and centrifuged at 10,000 g for 10 min at 4°C to pellet crude mitochondria and associated membrane structures. The resulting pellet was resuspended in 2 mL of MSB separation buffer. In an ultracentrifuge tube, 8 mL of PE organelle isolation density medium was pre-loaded, and the 2 mL sample suspension was carefully layered on top, followed by precise balance with MSB buffer (mass error < 0.001 g). Density gradient centrifugation was performed at 95,000g for 30 min at 4°C in a Beckman Optima ultracentrifuge (SW41 Ti swinging-bucket rotor). The diffuse white MAM band at the upper interface was carefully aspirated, diluted with 10 volumes of MSB buffer, and centrifuged at 6,300g for 10 min at 4°C to eliminate residual dense contaminants. The supernatant was transferred to a fresh tube and ultracentrifuged at 100,000g for 1 h at 4°C to pellet purified MAMs. Pellets were resuspended in MSB buffer, assessed for biochemical purity by immunoblotting against PACS2 (MAMs marker) (Immunoway, YN5211, 1:1,000), VDAC1 (outer mitochondrial membrane) (Immunoway, PT0214R, 1:2,000), Calnexin (ER)(Immunoway, YT0613, 1:2,000), and GAPDH (cytosolic negative control (Immunoway, PT0582R, 1:2,000); Supplementary Fig. S11), and promptly subjected to downstream targeted lipidomic profiling.

### Targeted Subcellular Lipidomic Profiling

Lipids from whole-cell inputs and purified MAM fractions were extracted using an optimized methanol/methyl tert-butyl ether (MTBE) liquid-liquid extraction method. Briefly, samples normalized to equal protein amounts were spiked with an internal standard mixture of stable isotope-labeled phospholipids, mixed with methanol and MTBE, sonicated in an ice-water bath for 20 min, and phase-separated by adding mass spectrometry-grade water. Following centrifugation at 12,000g for 10 min at 4°C, the upper organic phase containing lipids was harvested, dried under a gentle nitrogen stream, and reconstituted in 90% isopropanol/acetonitrile. Chromatographic separation was achieved on a Shimadzu LC-30AD UHPLC system equipped with Phenomenex Kinetex C18 and Phenomenex Luna NH2 columns. Targeted mass spectrometric detection was performed on an AB SCIEX QTRAP 7500+ triple quadrupole system utilizing electrospray ionization (ESI) operating in both positive and negative multiple reaction monitoring (MRM) modes. Chromatographic peak identification and integration were conducted using Sciex OS software to calculate absolute abundances of individual lipid species normalized to their respective internal standards. Variable Importance in Projection (VIP) scores were evaluated, and the organellar enrichment coefficient for core PUFA-PLs was calculated as the ratio of normalized abundance in purified MAMs to that in the corresponding whole-cell input (*EnricℎmentScrore* = *Abundance_MAMs_*/*Abundance_input_*) using MetaboAnalyst 5.0.

### Immunoblotting, Co-immunoprecipitation (Co-IP), and IP-MS

For immunoblotting, cultured cells or tissue samples were lysed in RIPA buffer supplemented with protease and phosphatase inhibitor cocktails (Roche). Protein concentrations were quantified using a BCA protein assay kit (Thermo Fisher Scientific). Equal amounts of protein (20-50 μg) were resolved by SDS-PAGE and transferred onto PVDF membranes (Millipore). Membranes were blocked with 5% non-fat milk in TBST for 1 h and incubated overnight at 4°C with primary antibodies against ferroptosis and lipid metabolic surveillance markers: anti-GPX4 (ImmunoWay, Cat# YM8430, 1:2,000), anti-SLC7A11 (ImmunoWay, Cat# PT0398R, 1:2,000), anti-ACSL4 (ImmunoWay, Cat# PT0448R, 1:2,000), and anti-LPCAT2 (Proteintech, Cat# 15082-1-AP, 1:2,000). Membranes were also probed with antibodies directed against structural complex components and subcellular markers: anti-ANO6 (Thermo Fisher Scientific, Cat# PA5-58610, 1:2,000), anti-Vimentin (Proteintech, Cat# 60330-1-Ig, 1:10,000), anti-VAPB (Proteintech, Cat# 66191-1-Ig, 1:2,000), anti-PTPIP51 (Proteintech, Cat# 20641-1-AP, 1:2,000), anti-PACS2 (ImmunoWay, Cat# YN5211, 1:1,000), anti-VDAC1 (ImmunoWay, Cat# PT0214R, 1:2,000), anti-Calnexin (ImmunoWay, Cat# YT0613, 1:2,000), anti-FLAG (Sigma-Aldrich, Cat# F1804, 1:5,000), anti-HA (Cell Signaling Technology, Cat# 3724, 1:2,000), anti-β-actin (Proteintech, Cat# 66009-1-Ig, 1:10,000), anti-β-tubulin (ImmunoWay, Cat# PT0505R, 1:2,000), and anti-GAPDH (ImmunoWay, Cat# PT0582R, 1:2,000). Blots were incubated with HRP-conjugated secondary antibodies (1:5,000) for 1 h and developed using ECL chemiluminescent substrates.

For co-immunoprecipitation (Co-IP), direct immunoprecipitation was conducted using the Pierce Direct IP Kit (Thermo Fisher Scientific, Cat# 26148) according to the manufacturer’s protocol. Specific antibodies targeting structural components (anti-ANO6, anti-VIM, anti-VAPB, anti-FLAG, or anti-HA) and control IgG were covalently cross-linked onto AminoLink Plus coupling resin to prevent antibody co-elution and avoid IgG heavy and light chain interference during subsequent immunoblot detection. Pre-cleared cell lysates were incubated with the immobilized antibody-coupled resin overnight at 4°C with gentle agitation. Interacting immune complexes were eluted by boiling in SDS sample loading buffer and subsequently analyzed by immunoblotting, enabling unobstructed detection of co-precipitated targets near immunoglobulin heavy and light chain molecular weights.

For the IP-MS interactome, immunoprecipitated ANO6 complexes were reduced with DTT, alkylated with iodoacetamide, and digested with sequencing-grade modified trypsin (1:50 w/w) for 20 h at 37°C. Peptides were separated on a nanoElute UHPLC system and analyzed on a timsTOF Pro mass spectrometer operating in PASEF mode. Raw data were searched against UniProt databases using MaxQuant 1.6.14 with an FDR cutoff <0.01.

### Viability, Apoptosis, and Liposome Rescue

For cell viability assays, breast cancer cells were seeded into 96-well plates (5 × 10^3^ cells/well) and allowed to adhere overnight. Cells were exposed to concentration gradients of paclitaxel (PTX; MedChemExpress, Cat# HY-B0015), RSL3 (MedChemExpress, Cat# HY-100218A), or niclosamide (MedChemExpress, Cat# HY-B0497) for 48 h. Following treatment, 10 μL of CCK-8 reagent (BioSharp, Cat# BS350A) was added to each well and incubated at 37°C for 1-2 h. Absorbance at 450 nm was measured using a microplate spectrophotometer, and relative cell viability was calculated normalized to vehicle-treated controls.

To discriminate non-lethal phosphatidylserine (PS) externalization from true apoptotic membrane permeabilization, dual staining with Annexin V-FITC and propidium iodide (PI) (Elabscience, Cat# E-CK-A211) was conducted. Vector control and ANO6-OE cells seeded in 6-well plates were treated with vehicle or PTX (5 nM) for 48 h. Both floating and adherent cells were harvested, with adherent cells gently detached using 0.25% trypsin without EDTA to preserve cell surface membrane integrity. Harvested cells were washed twice with cold PBS and resuspended in 500 μL of 1X Binding Buffer. Cell suspensions were stained with 5 μL Annexin V-FITC and 5 μL PI for 15 min at room temperature in the dark, followed by flow cytometric acquisition on a flow cytometer (Beckman Coulter CytoFLEX) within 1 h. Flow cytometric gating quadrants were defined to resolve distinct membrane states: Q4 (Annexin V^−^/PI^−^) represents viable cells with preserved inner leaflet PS asymmetry; Q3 (Annexin V^+^/PI^−^) delineates viable cells undergoing non-lethal PS scrambling without loss of plasma membrane integrity; and Q2 (Annexin V^+^/PI^+^) designates terminal cell death with ruptured membranes and complete loss of viability.

For artificial liposome preparation and rescue experiments, 1,2-dioleoyl-sn-glycero-3-phosphocholine (DOPC; Sigma-Aldrich, Cat# 850375P) and 1,2-dioleoyl-sn-glycero-3-phospho-L-serine (DOPS; MedChemExpress, Cat# HY-141613) were dissolved in absolute ethanol at a 7:3 molar ratio. The lipid mixture was rapidly injected into vortexing PBS to achieve an initial 0.8 mM liposome suspension, followed by extrusion through 0.22-μm polyethersulfone (PES) syringe filters. Cells subjected to ferroptosis induction were supplemented with 20-100 μM liposomes to assess functional membrane protection and viability rescue.

### In Vivo Orthotopic Xenograft Models

All animal procedures were approved by the Animal Experimental Ethics Committee of Kunming Medical University (Protocol No. kmmu20230811). Female BALB/c mice (6-8 weeks old) were maintained under specific pathogen-free conditions. To model established pre-existing tumors and clinical neoadjuvant chemotherapy (NAC) resistance, EMT6-Vector or EMT6-ANO6-OE cells (1 × 10^6^ cells) were orthotopically inoculated into the fourth mammary fat pad of mice. Given differing tumor growth kinetics, therapeutic intervention was initiated when tumors reached a predefined bulk volume. Mice were then treated with vehicle or paclitaxel (PTX; 5 mg/kg administered intraperitoneally every 3 days for 8 cycles; Day 24-48 for ANO6-OE and Day 30-54 for Vector). Tumor dimensions were measured every 6 days using digital calipers, and volumes were calculated as *Volume* = 0.5 × *Lengt*ℎ × *Widt*ℎ^2^. Baseline diagnostic tumors (Pre-PTX: Day 24 for ANO6-OE, Day 30 for Vector) and post-treatment residual lesions (Post-PTX: Day 48 for ANO6-OE, Day 54 for Vector) were resected (n = 5 mice per group), photographed, weighed, and subjected to comparative multi-omics and histopathological profiling.For the conformational mutant progression model, mice were orthotopically inoculated into mammary fat pads with Vector, ANO6-OE, ANO6-KO, active scaffold I612A, or scaffold-compromised mutant D703R cells (n = 5 mice per group), and tumor growth kinetics were monitored for 15 days prior to terminal resection. For combination efficacy studies, mice bearing EMT6-ANO6-OE orthotopic tumors were randomized into four cohorts (n = 5 per group): Vehicle (5% DMSO, 5% Tween 80, 40% PEG-300, and 50% sterile water), Niclosamide alone (20 mg/kg), RSL3 alone (5 mg/kg), or combined Niclosamide (20 mg/kg) plus RSL3 (5 mg/kg) administered intraperitoneally every 3 days for 8 cycles.

### Multiplex Tissue Immunofluorescence Staining and Cross-Sectional Line-Scan Profiling

Multiplex immunofluorescence staining of fresh-frozen tissue sections (8-10μm thick) from clinical breast cancer specimens and excised xenograft tumors was performed using the Mouse/Rabbit 3-Plex 4-Color Fluorescence Detection Kit (ImmunoWay Biotechnology, Cat# RS0035) via sequential tyramide signal amplification (TSA) according to the manufacturer’s instructions. Frozen sections were air-dried at room temperature, fixed in ice-cold 4% paraformaldehyde for 15 min, and permeabilized with 0.3% Triton X-100 in PBS for 10 min. Endogenous peroxidase activity was quenched with 3% H_2_O_2_ for 15 min, and non-specific binding was blocked with 5% normal goat serum for 1 h at room temperature. Sections were subjected to sequential rounds of target detection: primary antibodies (anti-ANO6, Thermo Fisher Scientific, Cat# PA5-58610, 1:200; anti-VAPB, Proteintech, Cat# 66191-1-Ig, 1:200; and anti-Vimentin, Proteintech, Cat# 60330-1-Ig, 1:500) were sequentially applied overnight at 4°C, followed by incubation with corresponding HRP-conjugated secondary antibodies and catalytic deposition of designated fluorophore-tyramides (TSA-488, TSA-594, and TSA-647). Between successive staining cycles, antibody complexes were thermally stripped in sodium citrate buffer (pH 6.0) using microwave heating to prevent cross-reactivity. Cell nuclei were counterstained with DAPI (Sigma-Aldrich, Cat# D9542) for 10 min in the dark. Multi-channel fluorescent confocal micrographs were captured on a Zeiss LSM 880 laser scanning confocal microscope equipped with a 20× immersion objective in sequential scanning mode to avoid spectral bleed-through. Spatial cross-correlation and co-localization profiles along cross-sectional cell transects were analyzed using the Plot Profile tool in ImageJ/Fiji software to track synchronized fluorescence intensity peaks across organelle contact sites.

### Bioinformatics and Statistical Analysis

Differential gene expression from clinical transcriptomic datasets was analyzed using the DESeq2 package with thresholds set at log_2_FC > 1 and adjusted P(FDR) < 0.05. Gene Set Enrichment Analysis (GSEA) was performed via the clusterProfiler package using canonical collections from the Molecular Signatures Database (MSigDB; normalized enrichment score NES > 1.5, FDR < 0.05). Kaplan-Meier survival trajectories and hazard ratios (HR) with 95% confidence intervals were evaluated using log-rank tests based on clinical breast cancer cohorts from TCGA and Kaplan-Meier Plotter databases. Targeted lipidomic and metabolomic pathway topological impact analyses, orthogonal partial least squares-discriminant analysis (OPLS-DA), and VIP rankings were conducted using MetaboAnalyst 6.0 (https://www.metaboanalyst.ca/). Endogenous RNA and protein expression profiles of ANO6, VIM, and VAPB across diverse human cancer cell lines were retrieved from The Human Protein Atlas (HPA, version 23.0; https://www.proteinatlas.org) to guide cell line selection for pan-cancer validation.

Experimental data are presented as mean ± standard deviation (SD) or mean ± standard error of the mean (SEM) from at least three independent biological replicates. Prior to parametric testing, data normality and homogeneity of variance were assessed using the Shapiro-Wilk test and Brown-Forsythe test, respectively. Pairwise comparisons between two groups were evaluated using two-tailed unpaired or paired Student’s t-tests as appropriate (paired t-tests applied for matched pre- and post-treatment clinical biopsies and paired tissue specimens). Differences among multiple experimental groups were assessed by one-way ANOVA followed by Tukey’s or Dunnett’s post-hoc multiple comparison test. Longitudinal in vivo tumor growth trajectories and combinatorial drug responses were analyzed using two-way repeated-measures ANOVA with interaction terms (P_interaction_ < 0.05 considered indicative of pharmacological synergy). Statistical analyses and graphical illustrations were executed using GraphPad Prism (version 9.5) and R (version 4.2.2). Thresholds for statistical significance were designated as *P < 0.05,**P < 0.01, ***P < 0.001, and ****P < 0.0001; ns indicates not statistically significant (P ≥ 0.05).

## Result

### Glycerophospholipid remodeling and ANO6 upregulation characterize chemoresistant breast cancer lesions and alter ferroptotic vulnerability

To delineate the metabolic rewiring underlying breast cancer persistence under neoadjuvant chemotherapy (NAC), we performed integrated transcriptomic and metabolomic profiling on paired primary breast cancer specimens collected prior to treatment (Pre-NAC) and from residual resistant lesions following chemotherapy (Post-NAC) (Fig. 1A; clinical characteristics in Supplementary Table S1A). Unsupervised clustering revealed substantial metabolic reprogramming in Post-NAC residual tumors, marked by a pronounced suppression of the tricarboxylic acid (TCA) cycle and amino acid metabolism, alongside a marked hyperactivation of lipid and glycerolipid metabolism (Fig. 1A). Gene Set Enrichment Analysis (GSEA) further corroborated that lipid biosynthetic cascades—particularly triglyceride, neutral lipid, and glycerolipid metabolic processes—were significantly enriched in chemotherapy-refractory tumors (NES > 2, FDR < 0.05; Fig. 1B). Gene Ontology molecular function (GO-MF) analysis identified phospholipid scramblase activity and phospholipid transporter activity among the top dysregulated functions (Fig. 1C). Integrative pathway-impact analysis pinpointed glycerolipid metabolism as the most prominently perturbed metabolic axis in post-chemotherapy residual tissues (Fig. 1D). Metabolic flux reconstruction highlighted a shift away from glycolytic pyruvate-driven TCA oxidation toward glycerol-3-phosphate (G3P)-dependent glycerophospholipid biosynthesis, wherein AGPAT1 and ANO6 (TMEM16F) govern critical intermediate steps (Fig. 1E). Untargeted metabolomic profiling further confirmed that G3P, along with multiple saturated and medium-chain fatty acids, exhibited marked accumulation in post-NAC residual lesions (Supplementary Fig. S1A-C). Parallel proteomic analysis substantiated this phenotypic shift, demonstrating robust enrichment of extracellular matrix, adipogenesis, and glycerolipid metabolic pathways alongside widespread suppression of cell cycle and apoptotic cascades (Supplementary Fig. S2A-E).

**Figure 1.**
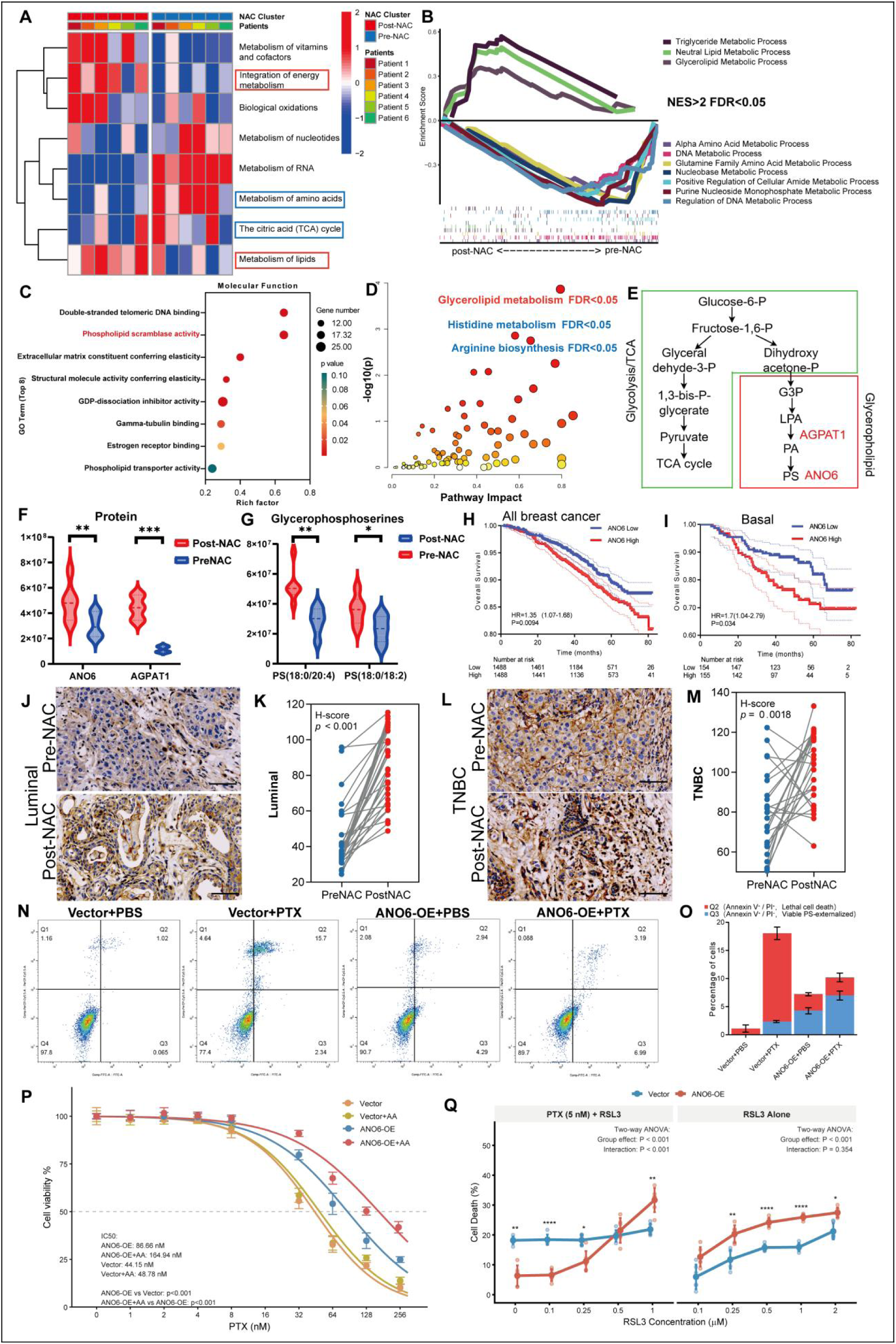
ANO6-mediated glycerophospholipid remodeling promotes chemoresistance and alters ferroptotic vulnerability in breast cancer. (A) Hierarchical clustering heatmap of differentially enriched metabolic pathways between paired pre- and post-neoadjuvant chemotherapy (NAC) breast cancer patient tissues (n = 6 pairs; Supplementary Table S1A). The red bounding box highlights pathways involved in energy integration and lipid metabolism. (B) Gene Set Enrichment Analysis (GSEA) plot displaying marked enrichment of lipid metabolic processes, including triglyceride and glycerolipid pathways, in post-NAC residual specimens (NES > 2.0, FDR < 0.05). (C) Bubble chart of Gene Ontology Molecular Function (GO-MF) terms for significantly altered genes, with phospholipid scramblase activity highlighted. (D) Integrative joint pathway analysis illustrating significant perturbations in glycerolipid metabolism (FDR < 0.05). (E) Schematic diagram illustrating metabolic rewiring: redirection of glycolytic carbon from the TCA cycle toward glycerol-3-phosphate (G3P) for glycerophospholipid biosynthesis, highlighting enzymatic nodes governed by AGPAT1 and ANO6. (F) Relative protein abundance of ANO6 and AGPAT1 derived from proteomic profiling of matched pre-NAC versus post-NAC patient tumor specimens (n = 6 pairs, Supplementary Table S1A; evaluated by paired two-tailed Student’s t-test). (G) Targeted lipidomic quantification of polyunsaturated glycerophosphoserine species [PS(18:0/20:4) and PS(18:0/18:2)] in paired pre- and post-NAC tissues (n = 6 pairs; paired two-tailed Student’s t-test). (H, I) Kaplan-Meier overall survival (OS) curves of breast cancer patients stratified by ANO6 mRNA expression in the complete breast cancer cohort (H) and the basal-like subtype (I) from public datasets. Hazard ratios (HR) with 95% confidence intervals and log-rank P values are indicated. (J-M) Representative immunohistochemistry (IHC) images and quantitative H-scores of ANO6 in paired pre-NAC versus post-NAC specimens from luminal (J, K; n = 35 pairs, Supplementary Table S1B; P < 0.001) and triple-negative breast cancer (L, M; n = 25 pairs, Supplementary Table S1C; P = 0.0018) cohorts by paired Student’s t-test. Scale bar = 100 μm. (N, O) Representative flow cytometry plots (Annexin V-FITC/PI; N) and stacked bar graph (O) depicting the percentage of terminal cell death (Q2: Annexin V⁺/PI⁺, red) and non-lethal, viable PS-externalizing populations (Q3: Annexin V⁺/PI⁻, blue) in control (Vector) and ANO6-overexpressing (ANO6-OE) cells following vehicle (PBS) or paclitaxel (5 nM PTX) challenge. (P) Cell viability dose-response curves and fitted half-maximal inhibitory concentration (IC₅₀) values of Vector and ANO6-OE cells treated with increasing concentrations of PTX in the presence or absence of exogenous arachidonic acid (AA) (statistical differences evaluated by two-way ANOVA). (Q) Percentage of cell death in Vector and ANO6-OE cells treated with indicated concentration gradients of RSL3 alone (right) or in combination with PTX (5 nM, left). Statistical significance was evaluated by two-way ANOVA (group effect and interaction P values are shown). Data are presented as mean ± SD from three independent biological replicates unless otherwise noted. *P < 0.05, **P < 0.01, ***P < 0.001, ****P < 0.0001; ns, not significant.

Validation across paired clinical specimens confirmed that the protein levels of both ANO6 and AGPAT1 were substantially elevated in Post-NAC residual tissues compared with their pre-treatment counterparts (Fig. 1F). Targeted lipidomic analysis demonstrated a selective enrichment of polyunsaturated phosphatidylserines, specifically PS(18:0/20:4) and PS(18:0/18:2), in Post-NAC tumors (Fig. 1G). Clinical survival analysis in large-scale patient cohorts revealed that high ANO6 transcript abundance was significantly associated with shortened overall survival across all breast cancer subtypes (HR = 1.35, P = 0.0094; Fig. 1H), with this adverse prognostic association being particularly pronounced in basal-like breast cancer (HR = 1.71, P = 0.034; Fig. 1I). Automated digital immunohistochemical (IHC) quantification of matched biopsy pairs reaffirmed that ANO6 protein was markedly upregulated in residual tumors following NAC across both luminal (Fig. 1J, K; Supplementary Table S1B; P < 0.001) and triple-negative breast cancer (TNBC) subtypes (Fig. 1L, M; Supplementary Table S1C; P = 0.0018).

Because ANO6 is canonically known as a Ca²⁺-dependent scramblase facilitating phosphatidylserine (PS) externalization during apoptosis, we next investigated whether its induction in resistant cells reflects an active apoptotic program. Flow cytometric profiling using Annexin V and propidium iodide (PI) revealed that paclitaxel (PTX) challenge provoked substantial membrane permeabilization and lethal cell death (15.7% Annexin V⁺/PI⁺, Q2) in Vector controls (Fig. 1N, O). In stark contrast, ANO6-OE cells exhibited minimal lethality (<3.2% Annexin V⁺/PI⁺), with only a minor fraction displaying non-lethal PS exposure (6.99% Annexin V⁺/PI⁻, Q3) (Fig. 1N, O). This distinct functional divergence confirmed that ANO6 upregulation does not execute lethal apoptosis under chemotherapy stress, but instead drives non-canonical membrane lipid remodeling in surviving cells. High-resolution in situ imaging using lactadherin-FITC corroborated that ANO6 overexpression promoted non-lethal, punctate cell-surface PS accumulation without compromising viability, whereas ANO6 knockout maintained baseline fluorescence (Supplementary Fig. S6, top panels).

Given that chemotherapy-driven glycerophospholipid accumulation introduces polyunsaturated acyl chains that can alter lipid metabolic vulnerability, we next evaluated whether ANO6 modulates cellular sensitivity to chemotherapy and ferroptotic stress. Dose-response cell viability assays demonstrated that ANO6-OE cells displayed markedly enhanced resistance to escalating doses of PTX compared with control cells (two-way ANOVA, P < 0.001; Fig. 1P). Exogenous supplementation of arachidonic acid (AA, an oxidizable PUFA driver) further augmented the survival advantage of ANO6-OE cells under PTX challenge, increasing their IC₅₀ from 86.86 nM to 164.94 nM, whereas control cells showed minimal alteration (Fig. 1P). To assess whether this lipid remodeling incurs an exploitable liability, we subjected cells to the GPX4 inhibitor RSL3. Cell death profiling revealed that while ANO6 overexpression buffered basal cytotoxicity induced by low-dose PTX (5 nM), it rendered cells significantly more vulnerable to escalating doses of RSL3 both in the presence and absence of PTX, culminating in pronounced ferroptotic cell death (Fig. 1Q; two-way ANOVA, interaction P < 0.001 for PTX + RSL3). Together, these findings demonstrate that chemoresistant breast cancer cells upregulate ANO6 to rewire glycerophospholipid metabolism, thereby shielding cells against chemotherapy-induced cytotoxicity while simultaneously creating an exploitable ferroptotic vulnerability under GPX4 inhibition.

### ANO6 drives chemoresistance in vivo and restrains ferroptotic execution through dynamic lipid droplet remodeling and PUFA-PL packaging

To establish the physiological relevance of ANO6 in chemotherapy persistence in vivo, we orthotopically inoculated control (Vector) or ANO6-overexpressing (ANO6-OE) breast cancer cells into the mammary fat pads of immunocompetent mice, followed by paclitaxel (PTX) administration upon reaching established bulk tumors (Fig. 2A, B). While Vector tumors exhibited pronounced growth arrest and therapeutic regression during PTX treatment, ANO6-OE tumors displayed sustained, refractory persistence without regression throughout the therapeutic timeframe (Fig. 2A, B; two-way ANOVA, ****P < 0.0001). Untargeted metabolomic and lipidomic profiling of resected xenograft tissues before and after PTX treatment revealed pronounced shifts in the cellular lipidome (Fig. 2C-E). Pathway impact scores and fold-change distributions demonstrated extensive rewiring within glycerophospholipids, eicosanoids, and diacylglycerols across the comparative cohorts (Fig. 2C-E). Detailed chromatographic profiling of individual phospholipid species unveiled that PTX challenge selectively triggered the accumulation of pro-ferroptotic polyunsaturated fatty acid-containing phospholipids (PUFA-PLs), including PUFA-esterified PC, PE, and PS specie, in ANO6-OE lesions, whereas monounsaturated fatty acid-containing species (MUFA-PLs) remained largely unaltered (Fig. 2F). Consistent with this PUFA-selective acyl-chain remodeling, lysophosphatidylcholine acyltransferase 2 (LPCAT2) transcript abundance was significantly elevated in Post-PTX ANO6-OE tumors (Fig. 2G; **P < 0.01), with corresponding marked protein induction observed in PTX-treated ANO6-OE cells (Fig. 2H; full uncropped blots in Supplementary Fig. S10). Gene Set Enrichment Analysis confirmed the selective activation of the phosphatidylcholine acyl-chain remodeling pathway (the Lands cycle) in resistant lesions (Supplementary Fig. S4A-C), which positively correlated with ANO6 abundance in patient cohorts (Supplementary Fig. S4D). To delineate how resistant cells cope with this substantial accumulation of pro-ferroptotic substrates, we examined key regulatory nodes of the ferroptotic surveillance cascade (Fig. 2I). Immunoblotting revealed that chemotherapy challenge provoked a marked induction of the pro-ferroptotic fatty acid-activating enzyme ACSL4 selectively in ANO6-OE cells, providing the enzymatic driver for continuous PUFA-PL incorporation (Fig. 2I). Concurrently, chemotherapy challenge triggered pronounced compensatory upregulation of GPX4 and the cystine/glutamate antiporter SLC7A11 selectively in ANO6-OE cells, indicating an adaptive activation of the antioxidant defense system (Fig. 2I). However, given the persistent burden of unoxidized PUFA-PLs, we hypothesized that biochemical clearance alone is insufficient and that resistant cells cooperatively deploy an organellar spatial strategy to physically compartmentalize and shelter lethal substrates away from membrane peroxidation hotspots. In support of this spatial compartmentalization hypothesis, tracking lipid flux using the lipophilic fluorescent probe DiI demonstrated that ANO6 overexpression accelerated cellular lipid uptake and internal packaging under PTX stress, which was further boosted by exogenous arachidonic acid (AA) supplementation but abolished by the selective LPCAT2 inhibitor TSI-01 (Fig. 2J, K; P < 0.001), confirming that this lipid packaging is enzymatically driven by LPCAT2-dependent acyl-chain remodeling. Transmission electron microscopy (TEM) further revealed marked intracellular ultrastructural remodeling: ANO6-OE cells harbored a prominent accumulation of cytoplasmic neutral lipid droplets (LDs) under PTX stress, whereas CRISPR-mediated ablation of ANO6 (ANO6-KO) significantly reduced LD abundance (Fig. 2L, M; P < 0.0001, P = 0.0367). Functionally, genetic knockout of ANO6 impaired this protective lipid sequestration, rendering cells hyper-susceptible to GPX4 inhibition by RSL3 alone or in combination with PTX (Fig. 2N; two-way ANOVA, group effect P < 0.001). Collectively, these data indicate that chemoresistant cells cope with lethal lipid burden through a dual mechanism: while mounting a compensatory antioxidant defense, they crucially rely on ANO6-dependent structural remodeling to package and sequester pro-ferroptotic lipids into lipid droplets, thereby evading lethal ferroptotic execution in vivo and in vitro.

**Figure 2.**
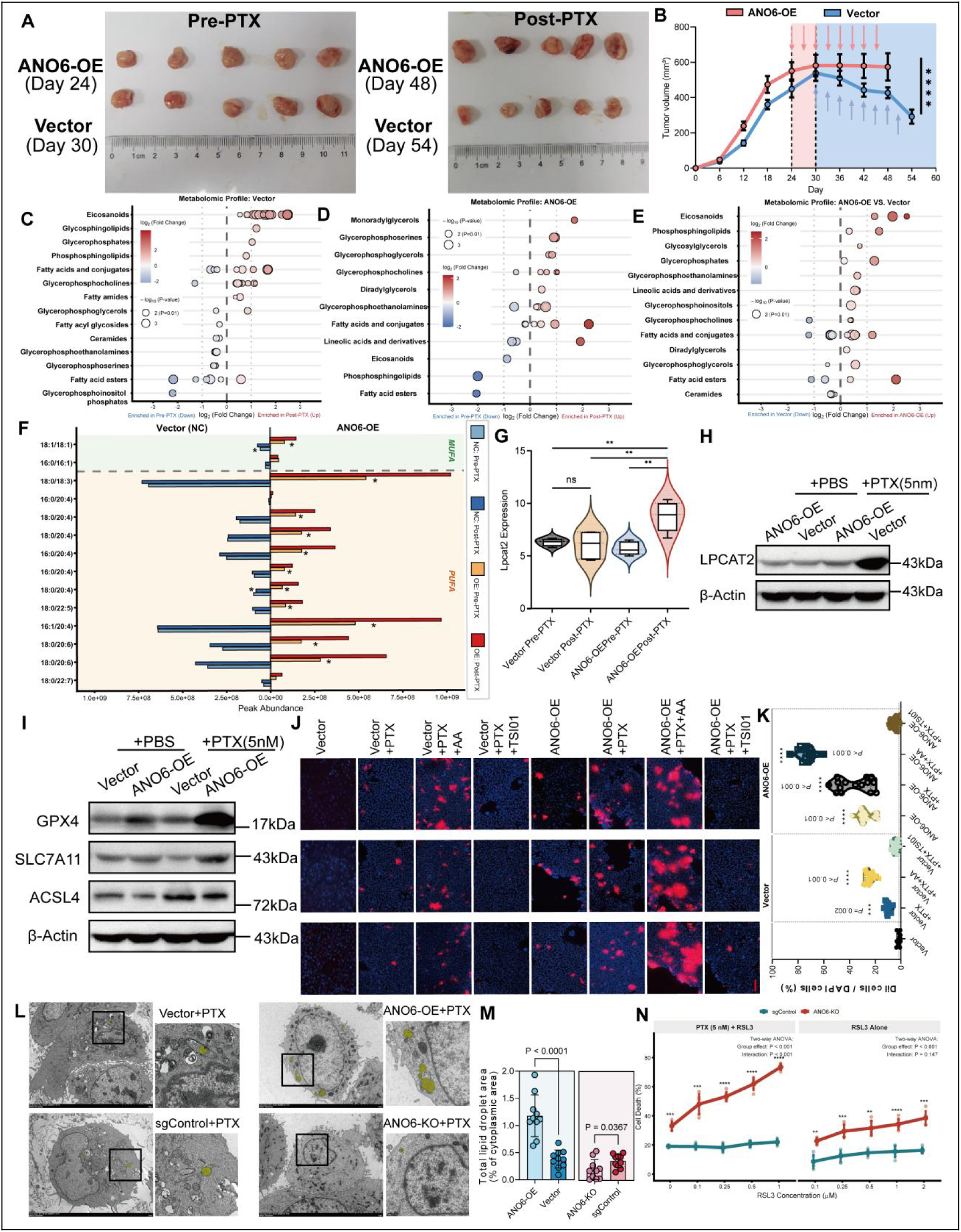
ANO6 promotes in vivo chemoresistance and circumvents ferroptosis via lipid droplet remodeling and PUFA-PL packaging. (A) Representative gross morphology of orthotopic xenograft tumors excised from BALB/c mice inoculated with Vector or ANO6-overexpressing (ANO6-OE) cells before paclitaxel treatment (Pre-PTX; Day 24 for ANO6-OE, Day 30 for Vector) and at the study endpoint (Post-PTX; Day 48 for ANO6-OE, Day 54 for Vector; scale in cm). (B) Longitudinal growth curves of Vector and ANO6-OE xenografts throughout the duration of PTX therapy (n = 5 mice per group; colored arrows indicate intraperitoneal PTX administration time points; **P < 0.0001 by two-way ANOVA). (C-E) Bubble plots illustrating untargeted metabolomic and lipidomic alterations across the comparative groups: Vector (Post- vs. Pre-PTX, C), ANO6-OE (Post- vs. Pre-PTX, D), and ANO6-OE versus Vector (Post-PTX, E). Dot color reflects log_2_(fold change); dot size corresponds to -log_10_(P value). (F) Chromatographic peak abundance of individual monounsaturated fatty acid (MUFA) and polyunsaturated fatty acid (PUFA) containing phospholipids (PC, PE, and PS species) quantified in Vector and ANO6-OE tumors under the indicated conditions (P < 0.05). (G, H) Relative mRNA expression of Lpcat2 in xenograft tumors across treatment cohorts (G; **P < 0.01; ns, not significant by one-way ANOVA) and representative Western blot analysis of LPCAT2 protein levels in Vector and ANO6-OE cells following vehicle (PBS) or PTX (5 nM) treatment (H; β-actin served as a loading control). (I) Immunoblot analysis of canonical ferroptosis surveillance proteins (GPX4, SLC7A11, ACSL4) in Vector and ANO6-OE cells treated with vehicle (PBS) or PTX (5 nM). β-actin served as a loading control. (J, K) Representative confocal fluorescence micrographs (J; scale bar = 200 μm) and violin plot quantification (K) of DiI fluorescent lipid probe uptake (red) counterstained with DAPI (blue) in Vector and ANO6-OE cells treated with vehicle, PTX, PTX + AA, or PTX + TSI-01 (LPCAT2 inhibitor; *P* < 0.001). (L, M) Representative transmission electron microscopy (TEM) micrographs depicting cytoplasmic lipid droplets (yellow pseudocolor; black boxes indicate magnified areas; scale bars indicated in micrographs) (L) and corresponding quantification of total lipid droplet area relative to cytoplasmic area (M) in Vector, ANO6-OE, sgControl, and ANO6-KO cells following PTX treatment (*P* < 0.0001, *P* = 0.0367 by two-tailed unpaired Student’s t-test). (N) Percentage of cell death in sgControl and ANO6-KO cells treated with indicated concentration gradients of the GPX4 inhibitor RSL3 alone (right) or in combination with PTX (5 nM, left). Statistical significance was determined by two-way ANOVA (group effect *P* < 0.001). Data are presented as mean ± SD from at least three independent biological replicates unless otherwise indicated. \**P* < 0.05, \*\**P* < 0.01, \*\*\**P* < 0.001, \*\*\*\**P* < 0.0001; ns, not significant.

### ANO6 scaffolds a VIM-VAPB structural tethering complex to orchestrate mitochondria-ER contact sites under chemotherapy stress

Given that chemoresistant cells remodel intracellular lipid architecture to package pro-ferroptotic lipids, we next explored the specific sub-organellar sites facilitating this compartmentalization. Because mitochondria-associated endoplasmic reticulum (ER) membranes (MAMs) serve as paramount hubs for phospholipid flux and organelle communication, we hypothesized that ANO6 functions at MAMs microdomains. Dual-color confocal imaging of mitochondria (MitoTracker, red) and the ER (ER-Tracker, green) demonstrated that paclitaxel (PTX) treatment markedly heightened ER-mitochondria spatial overlap and synchronized fluorescence line-scan intensity profiles in ANO6-overexpressing (ANO6-OE) cells relative to controls (Fig. 3A). Conversely, CRISPR-mediated knockout of ANO6 (ANO6-KO) severely disrupted ER-mitochondria alignment (Fig. 3A). Prior to functional assessments, targeted genetic knockout and stable reconstitution of conformational mutants (I612A and D703R) were verified by Western blot in EMT6 cells (Supplementary Fig. S3A, B; full uncropped blots in Supplementary Fig. S8). Reconstitution with the active scaffolding mutant I612A induced striking organellar co-alignment, whereas cells bearing the D703R mutant exhibited dispersed, non-overlapping signals comparable to their respective empty vector controls (Fig. 3A).

**Figure 3.**
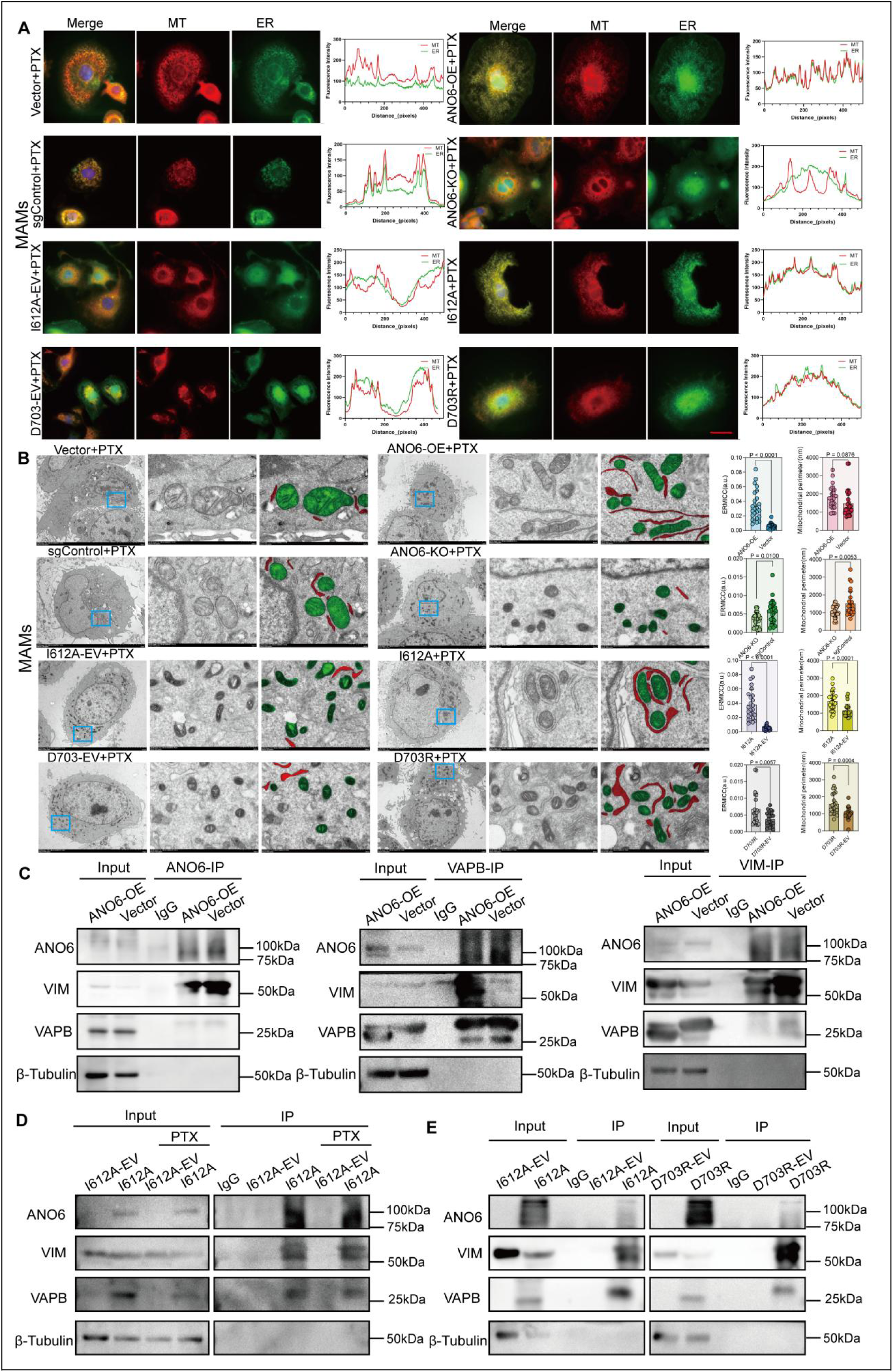
ANO6 recruits Vimentin and VAPB to assemble an active structural tethering complex and maintain MAMs architecture. (A) Representative confocal fluorescence micrographs and line-scan profiles depicting the spatial co-localization of mitochondria (MT, red) and the endoplasmic reticulum (ER, green) in Vector, ANO6-OE, sgControl, ANO6-KO, I612A-EV, I612A, D703R-EV, and D703R cells following paclitaxel (PTX) treatment (scale bar = 25 μm). Corresponding cross-sectional fluorescence intensity profiles (arbitrary units vs. distance in pixels) across indicated cellular transects are displayed on the right. (B) Representative transmission electron microscopy (TEM) micrographs visualizing mitochondria-associated endoplasmic reticulum membrane (MAM) contact sites under PTX challenge across the indicated genetic cohorts. Magnified contact zones (blue dashed boxes) show mitochondria and ER membranes pseudocolored in green and red, respectively (scale bars indicated in micrographs). Quantitative morphometric analyses of the ER-mitochondria contact coefficient (ERMICC, left plots) and mitochondrial perimeter (right plots; reflecting preservation of mitochondrial architecture versus ferroptotic mitochondrial shrinkage) determined from TEM images across experimental cohorts are shown on the right (P values indicated; ns, not significant by two-tailed unpaired Student’s t-test). (C) Reciprocal co-immunoprecipitation (Co-IP) assays detecting endogenous physical interactions among ANO6, Vimentin (VIM), and VAPB in Vector and ANO6-OE cells. Western blots display input and immunoprecipitated fractions obtained using anti-ANO6 (left), anti-VAPB (middle), or anti-VIM (right) antibodies, with IgG serving as the negative control and β-tubulin as the loading control. (D) Co-IP analysis assessing the interaction of the active scaffolding mutant I612A with endogenous VIM and VAPB in the presence or absence of paclitaxel (PTX) treatment. (E) Comparative Co-IP analysis evaluating the physical binding capacity of the hyper-scaffolding mutant I612A and the scaffold-compromised mutant D703R toward VIM and VAPB relative to their matched empty vector controls (I612A-EV and D703R-EV). Data are presented as mean ± SD from at least three independent biological replicates. *P < 0.05, **P < 0.01, ***P < 0.001, ****P < 0.0001; ns, not significant.

To directly visualize organelle contact architecture at nanometer resolution, we performed transmission electron microscopy (TEM) under PTX challenge (Fig. 3B). Ultrastructural analysis demonstrated that ANO6 overexpression profoundly expanded the close juxtaposition between the ER and outer mitochondrial membranes, yielding a notable surge in the ER-mitochondria contact coefficient (ERMICC; P < 0.0001; Fig. 3B), whereas genetic ablation of ANO6 led to substantial contact disconnection (Fig. 3B; P < 0.0001). Concurrently, while control and KO cells displayed fragmented and shrunken mitochondria, a classic morphological hallmark of ferroptotic damage under chemotherapeutic stress, ANO6-OE significantly preserved mitochondrial perimeter, preventing chemotherapy-induced mitochondrial shrinkage (Fig. 3B). In structural mutant lines, cells harboring the active scaffolding mutant I612A displayed robustly elongated and tightened MAMs junctions (P < 0.0001, reaching ERMICC values up to 0.04) along with preserved mitochondrial architecture. Conversely, the D703R mutant exhibited only a marginal baseline tethering capacity that, despite reaching statistical significance over empty vector (Fig. 3B; P = 0.0057, and P = 0.0004 for perimeter), remained severely stunted and functionally compromised, failing to avert ferroptotic mitochondrial shrinkage compared with the pronounced contact expansion driven by I612A or ANO6-OE. Consistent with this phenotype, baseline ultrastructural profiling in human MDA-MB-231 breast cancer cells revealed that ANO6 overexpression alone was sufficient to markedly expand ERMICC, whereas CRISPR knockout significantly dismantled these contacts (Supplementary Fig. S7), demonstrating that ANO6 dictates organellar contact tightness across species.

To identify candidate scaffolding partners mediating this organelle contact, we performed immunoprecipitation coupled with mass spectrometry (IP-MS) using an anti-ANO6 antibody (Supplementary Fig. S5A). Unbiased intersection of ANO6-binding proteins, post-NAC upregulated genes, annotated MAMs components, and TCGA co-expression networks uniquely identified two core candidates: VIM and VAPB (Supplementary Fig. S5B-D). Validating this physical assembly, three-way reciprocal Co-IP assays confirmed robust endogenous mutual interactions among ANO6, VIM, and VAPB in ANO6-OE cells, demonstrating that ANO6 physically bridges the intermediate filament VIM and the ER tether VAPB into a ternary structural tethering complex (Fig. 3C; full uncropped blots in Supplementary Fig. S9).

To determine whether this structural nucleation depends on chemotherapy stress and specific conformational states, we interrogated the mutant constructs. Co-IP assays confirmed that cells harboring the active mutant I612A maintained baseline complex formation that was substantially enhanced upon PTX challenge (Fig. 3D). In striking contrast, parallel comparative Co-IP demonstrated that while the active scaffolding conformation I612A potently co-precipitated both VIM and VAPB, the catalytically inactive D703R mutant displayed a marked reduction in its physical binding capacity toward both partners (Fig. 3E), indicating that the closed/calcium-compromised mutant conformation compromises stable structural complex assembly. Collectively, these structural and biochemical data establish that ANO6 acts as a molecular scaffold at organelle interfaces: upon chemotherapy challenge, it engages an active scaffolding conformation to directly recruit VIM and VAPB, nucleating a tethering complex that mechanically stabilizes MAMs architecture.

### The active structural scaffolding conformation of ANO6 governs lipid packaging, ferroptosis evasion, and in vivo tumor progression

Having established that ANO6 physically interacts with VIM and VAPB, we investigated how its well-characterized conformational states, represented by the open-gate, constitutively active mutant (I612A) and the calcium-binding-compromised mutant (D703R)^[19]^, impact this inter-organellar architecture and cell fate. We evaluated cell survival across engineered structural mutant models: the active scaffolding mutant (I612A) and the scaffolding-compromised mutant/catalytically inactive mutant (D703R). Dose-dependent cell death profiling demonstrated that reconstitution with I612A conferred potent protection against paclitaxel (PTX)-induced cytotoxicity (Fig. 4A) as well as GPX4 inhibition by RSL3 (Fig. 4B). In sharp contrast, cells harboring the scaffold-compromised mutant D703R mutant exhibited marked hypersensitivity to both agents, displaying elevated death rates indistinguishable from matched empty vector controls (Fig. 4A, B). Strikingly, exogenous supplementation with serine-containing liposomes (DOPC-DOPS) selectively rescued D703R-expressing cells from RSL3-induced ferroptotic lethality, whereas neutral DOPC alone failed to confer protection (Fig. 4C; two-way ANOVA, interaction P < 0.001). Ultrastructural transmission electron microscopy (TEM) further revealed that I612A-expressing cells accumulated abundant cytoplasmic lipid droplets (LDs) under chemotherapy pressure, whereas D703R cells displayed negligible LD formation comparable to baseline vector levels (Fig. 4D, E; P = 0.0010 for I612A vs. I612A-EV, P = 0.0476 for D703R vs. D703R-EV). In line with this functional restoration, in situ lactadherin imaging confirmed that the active scaffolding mutant I612A sustained cell-surface PS puncta, whereas the catalytically compromised mutant D703R mutant lacked surface PS exposure unless exogenously reconstituted with DOPC-DOPS liposomes (Supplementary Fig. S6, middle and bottom panels).

**Figure 4.**
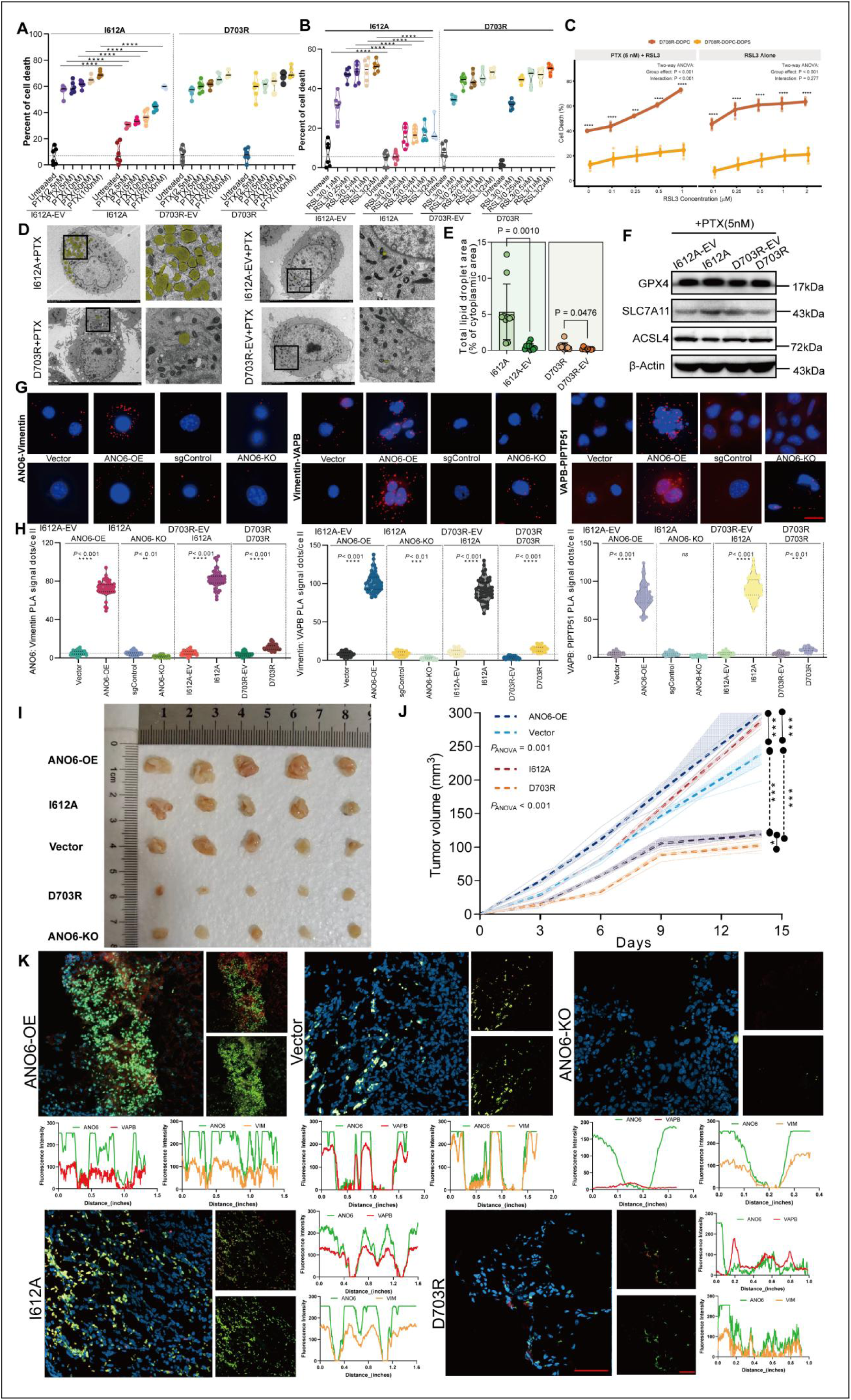
The active structural scaffolding conformation of ANO6 governs lipid packaging, ferroptosis resistance, and tumor progression.(A, B) Percentage of cell death in I612A, I612A-EV, D703R, and D703R-EV cells exposed to concentration gradients of paclitaxel (PTX, 10-100 nM; A) or the GPX4 inhibitor RSL3 (0.25-2 μM; B) (****P < 0.0001 by two-way ANOVA). (C) Ferroptotic cell death curves of D703R cells treated with RSL3 alone (right) or in combination with PTX (5 nM, left) following exogenous supplementation with neutral DOPC or serine-containing DOPC-DOPS liposomes (two-way ANOVA, group effect *P < 0.0001, interaction P < 0.001). (D, E) Representative transmission electron microscopy (TEM) micrographs depicting intracellular lipid droplets (yellow pseudocolor; black boxes indicate magnified regions; scale bars indicated in micrographs) (D) and morphometric quantification of total lipid droplet area relative to cytoplasmic area (E) in I612A, I612A-EV, D703R, and D703R-EV cells under PTX challenge (P = 0.0010 for I612A vs. I612A-EV; P = 0.0476 for D703R vs. D703R-EV by two-tailed unpaired Student’s t-test). (F) Western blot analysis of canonical ferroptosis surveillance proteins (GPX4, SLC7A11, ACSL4) in I612A-EV, I612A, D703R-EV, and D703R cells following paclitaxel (5 nM PTX) challenge. β-actin served as the loading control. (G, H) Representative in situ proximity ligation assay (PLA) confocal micrographs (G; scale bar = 25 μm) and single-cell puncta violin plot quantifications (H) evaluating spatial proximity (<40 nm) of ANO6-Vimentin, canonical VAPB-PTPIP51, and Vimentin-VAPB across Vector, ANO6-OE, sgControl, ANO6-KO, I612A-EV, I612A, D703R-EV, and D703R cells. Red puncta denote spatial interaction sites; nuclei are counterstained with DAPI (blue) (P < 0.001, ****P < 0.0001; ns, not significant by one-way ANOVA). (I) Gross morphology photographs of orthotopic xenograft tumors excised at the study endpoint (Day 15) from mice inoculated with ANO6-OE, I612A, Vector, D703R, or ANO6-KO cells (n = 5 mice per group; scale in centimeters). (J) Longitudinal xenograft growth trajectories monitored over 15 days. Shaded ribbons represent 95% confidence intervals (P_ANOVA = 0.001 for ANO6-OE vs. Vector; P_ANOVA < 0.001 for I612A vs. D703R; two-way repeated-measures ANOVA). (K) Representative multi-color confocal immunofluorescence images (scale bar = 200 μm) and spatial cross-sectional line-scan intensity profiles evaluating spatial co-localization of ANO6, VAPB, and Vimentin within resected tumor sections across ANO6-OE, Vector, ANO6-KO, I612A, and D703R cohorts. Synchronized fluorescence intensity peaks along transects denote coordinated physical co-alignment. Data are presented as mean ± SD from at least three independent biological replicates. *P < 0.05, **P < 0.01, ***P < 0.001, ****P < 0.0001; ns, not significant.

To delineate the molecular response of structural mutants to chemotherapy stress, we assessed the canonical ferroptosis surveillance machinery under paclitaxel challenge (Fig. 4F). Immunoblotting revealed that upon PTX exposure, reconstitution with the active scaffolding mutant I612A promoted an adaptive upregulation of the key antioxidant defenses GPX4 and SLC7A11 relative to its empty vector control, accompanied by a marked downregulation of the pro-ferroptotic lipid-activating enzyme ACSL4 (Fig. 4F). Interestingly, while the scaffold-compromised mutant D703R also exhibited elevated GPX4 and SLC7A11 levels, it sustained high ACSL4 expression and, more critically, lacked the capacity to physically recruit VIM/VAPB and stabilize MAMs junctions, leaving cells fully susceptible to ferroptotic execution (Fig. 4A-C, F). This confirms that robust ferroptosis evasion requires the concerted integration of antioxidant surveillance, suppressive regulation of ACSL4-driven lipid peroxidation, and ANO6-dependent structural compartmentalization at MAMs.

We next tracked nanometer-scale complex assembly using in situ proximity ligation assays (PLA). Cells harboring I612A displayed extensive spatial PLA puncta clusters for ANO6-Vimentin, canonical MAMs tethering VAPB-PTPIP51, and Vimentin-VAPB, mirroring or exceeding ANO6-OE levels (Fig. 4G, H; P < 0.001). Conversely, D703R cells, similar to ANO6-KO, yielded only background-level interaction dots (Fig. 4G, H; P < 0.001). To determine whether this structural scaffolding dictates malignant progression in vivo, immunocompetent mice were inoculated with the panel of cell lines (Fig. 4I, J). Tumors derived from ANO6-OE and I612A cells exhibited rapid outgrowth, with volumes reaching approximately 300 mm³ by day 15 (Fig. 4I, J; P_ANOVA = 0.001 and P_ANOVA < 0.001, respectively). In stark contrast, D703R tumors exhibited stunted kinetics, remaining as small and indolent as ANO6-KO lesions (Fig. 4I, J). Multiplex confocal immunofluorescence and cross-sectional spatial line-scan profiling on resected tumor sections demonstrated synchronized fluorescence intensity peaks for ANO6, VAPB, and VIM in ANO6-OE and I612A tissues, whereas spatial co-alignment collapsed into baseline profiles in Vector, ANO6-KO, and D703R sections (Fig. 4K).

Together, these genetic, biochemical, and in vivo findings establish that the pro-survival capacity of ANO6 is strictly governed by its active structural scaffolding conformation, which orchestrates lipid droplet packaging and VIM-VAPB platform assembly, rather than its canonical scramblase enzymatic activity alone.

### Subcellular lipidomics of purified MAMs and patient cohorts demonstrate that ANO6 facilitates compartmentalized sequestration of pro-ferroptotic PUFA-PLs via its scaffolding conformation

Because the structural tethering complex assembled by ANO6, VIM, and VAPB stabilizes ER-mitochondria contact interfaces, we hypothesized that this microdomain functions as a physical sequestration reservoir to compartmentalize lethal pro-ferroptotic substrates away from membrane peroxidation hotspots. To test this directly, we isolated pure mitochondria-associated ER membrane (MAMs) fractions along with matched whole-cell lysates (Input) via subcellular differential ultracentrifugation from our panel of engineered cell models (Vector, ANO6-OE, sgControl, ANO6-KO, I612A-EV, I612A, D703R-EV, and D703R) under paclitaxel challenge (5 nM PTX, 24 h), followed by high-resolution targeted lipidomic profiling (Fig. 5A; MAMs biochemical purity verified by PACS2, VDAC1, and Calnexin enrichment in Supplementary Fig. S11).

**Figure 5.**
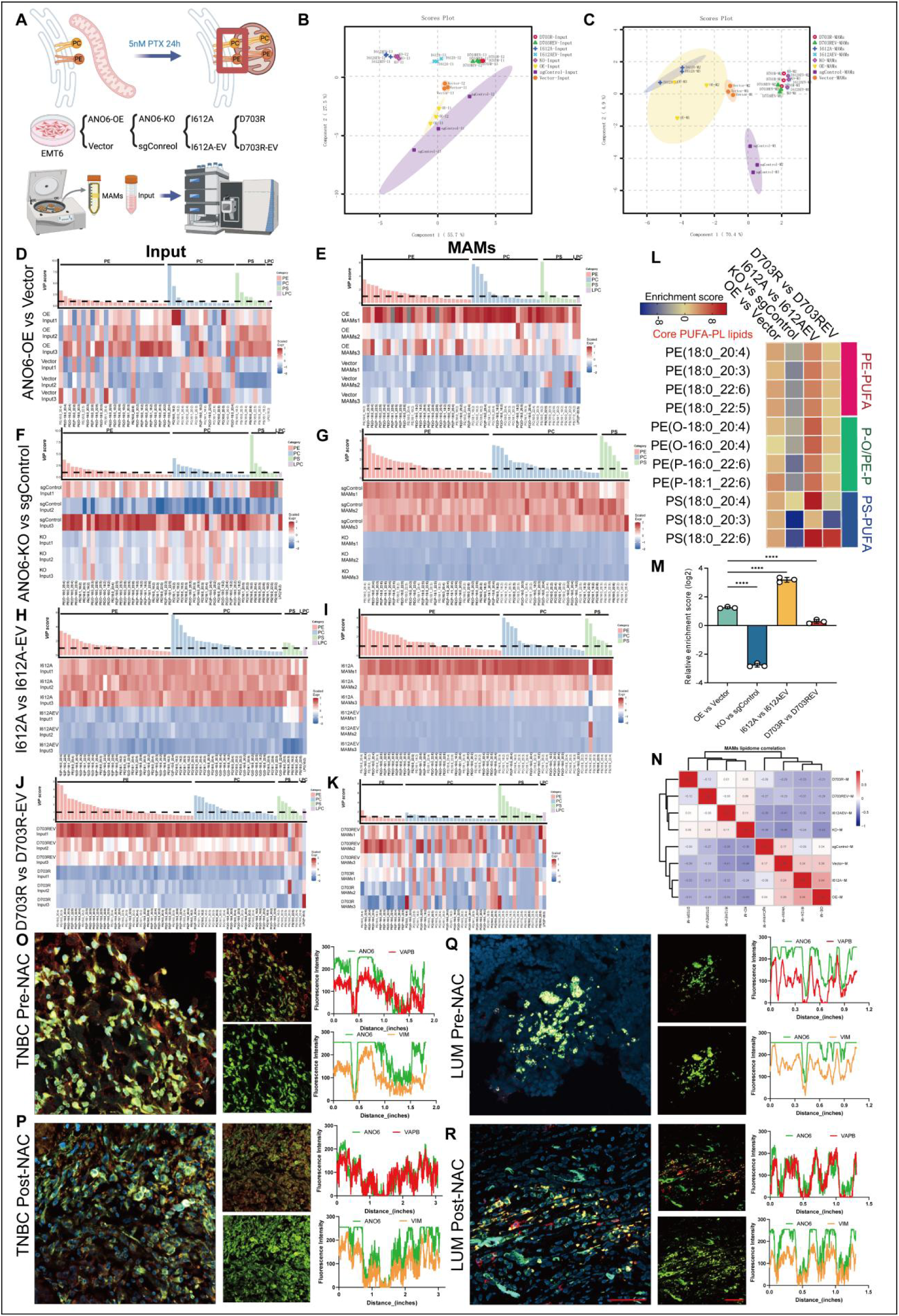
Subcellular lipidomics of purified MAMs and patient cohorts demonstrate that ANO6 facilitates compartmentalized sequestration of pro-ferroptotic PUFA-PLs via its scaffolding conformation. (A) Schematic diagram illustrating the workflow: isolation of whole-cell lysates (Input) and pure mitochondria-associated endoplasmic reticulum membrane (MAMs) fractions by differential ultracentrifugation from engineered EMT6 lines (Vector, ANO6-OE, sgControl, ANO6-KO, I612A-EV, I612A, D703R-EV, and D703R) exposed to paclitaxel (5 nM PTX, 24 h), followed by LC-MS/MS lipidomic analysis. (B, C) Principal component analysis (PCA) score plots derived from metabolic fingerprints of total cell Inputs (B) and purified MAMs fractions (C), illustrating distinct spatial lipid segregation among genetic cohorts. (D-K) Hierarchical clustering heatmaps and VIP score distributions displaying relative abundance changes of major phospholipid classes (PE, PC, PS, and LPC) in whole-cell Inputs (D, F, H, J) and purified MAMs fractions (E, G, I, K) across comparative cohorts (ANO6-OE vs. Vector, ANO6-KO vs. sgControl, I612A vs. I612A-EV, and D703R vs. D703R-EV). Red and blue denote relative upregulation and downregulation normalized by Z-score, respectively. (L) Heatmap depicting organellar enrichment scores of Core PUFA-PL species (including PE-PUFA, ether-linked PE-O/PE-P, and PS-PUFA) within isolated MAMs fractions across experimental cohorts. (M) Quantification of relative enrichment scores (log_2_) for Core PUFA-PLs sequestered within MAMs across cell models (****P < 0.0001 by one-way ANOVA). (N) Heatmap matrix representing Pearson correlation coefficients of MAMs lipidomic profiles across cell lines. (O-R) Representative multi-color confocal immunofluorescence images (scale bar = 200 μm) and spatial line-scan cross-sectional intensity profiles (fluorescence intensity in arbitrary units vs. distance in inches) evaluating the spatial co-localization of ANO6, VAPB, and VIM in matched pre-NAC and post-NAC residual tumor specimens from triple-negative breast cancer (TNBC; O, P) and luminal breast cancer (LUM; Q, R) patients. Data are presented as mean ± SD from at least three independent biological replicates. *P < 0.05, **P < 0.01, ***P < 0.001, ****P < 0.0001; ns, not significant.

Principal component analysis (PCA) revealed distinct spatial segregation and phenotypic clustering in both whole-cell inputs and purified MAMs fractions across the engineered models (Fig. 5B, C). Hierarchical clustering heatmaps of top-ranking phospholipids ranked by Variable Importance in Projection (VIP) scores uncovered that while whole-cell inputs exhibited moderate, generalized lipid shifts (Fig. 5D, F, H, J), the isolated MAMs fractions underwent marked, compartment-specific remodeling dominated by polyunsaturated phospholipid species (Fig. 5E, G, I, K).

To directly quantify the spatial partitioning of ferroptotic execution substrates, we evaluated the organellar enrichment profiles of Core PUFA-PL species, encompassing arachidonic acid (AA)- and docosahexaenoic acid (DHA)-esterified PE, ether-linked phospholipids (PE-O/PE-P), and PS species (Fig. 5L). Both ANO6 overexpression (ANO6-OE) and the active scaffolding mutant I612A drove massive physical trapping and selective enrichment of these lethal Core PUFA-PLs, such as PE(18:0_20:4), PE(18:0_22:6), and PS(18:0_20:4), specifically within the MAMs microenvironment (Fig. 5L, M; ****P < 0.0001). Conversely, CRISPR knockout of ANO6 resulted in significant depletion of Core PUFA-PLs from MAMs, yielding negative relative enrichment scores (Fig. 5L, M; ****P < 0.0001). Crucially, the scaffolding-compromised mutant D703R completely failed to compartmentalize pro-ferroptotic lipids, displaying minimal baseline enrichment indistinguishable from its control (Fig. 5L, M), confirming that the disruption of VIM-VAPB platform assembly abolishes organellar lipid sequestration capacity. Pearson correlation matrix analysis further confirmed that the MAMs lipidomic fingerprint of I612A closely mirrored that of ANO6-OE, while diverging sharply from KO and D703R profiles (Fig. 5N).

Finally, we interrogated the clinical relevance of this spatial assembly in paired patient specimens collected prior to neoadjuvant chemotherapy (Pre-NAC) and from residual chemoresistant lesions following chemotherapy (Post-NAC) across triple-negative breast cancer (TNBC) and luminal breast cancer cohorts (Fig. 5O-R). Multi-color confocal immunofluorescence and cross-sectional spatial line-scan profiling revealed minimal, disordered spatial overlap among ANO6, VAPB, and VIM in chemotherapy-naive biopsies (Fig. 5O, Q). In stark contrast, chemoresistant residual lesions from Post-NAC specimens exhibited tight spatial coordination, characterized by synchronized fluorescence intensity peaks for ANO6, VAPB, and VIM along cell transects in both TNBC and luminal subtypes (Fig. 5P, R). Together, these findings demonstrate that resistant breast cancer cells exploit the structural scaffolding conformation of ANO6 to assemble a MAMs-localized lipid reservoir, physically isolating lethal PUFA-PLs to evade ferroptosis under chemotherapy pressure.

### Vimentin serves as an indispensable structural pillar for the ANO6-nucleated MAMs complex and ferroptosis evasion

Having confirmed that ANO6 physically associates with Vimentin (VIM) and VAPB to form a tethering platform at MAMs, we investigated whether VIM acts as an obligatory structural pillar required for the integrity of this sequestration compartment. To test this, we transfected siRNA targeting Vimentin (siVimentin) into ANO6-OE and hyper-scaffolding I612A cells under paclitaxel (PTX) challenge, confirming efficient depletion of VIM protein without altering ANO6 abundance (Supplementary Fig. S3C). Transmission electron microscopy (TEM) revealed that while control cells (siNC) sustained elongated ER-mitochondria junctions, silencing VIM triggered extensive disassembly of MAMs, causing a pronounced drop in the ER-mitochondria contact coefficient (ERMICC; P < 0.0001) alongside reduced mitochondrial perimeters in both ANO6-OE (P = 0.0026) and I612A lines (P = 0.0168) (Fig. 6A). Concurrently, VIM knockdown resulted in a significant loss of cytoplasmic lipid droplets (LDs), reversing the lipid packaging phenotype driven by ANO6 overexpression or the I612A mutation (Fig. 6B; P = 0.0032 for ANO6-OE; P = 0.0216 for I612A). We next evaluated spatial tethering using in situ proximity ligation assays (PLA). In both ANO6-OE and I612A background cells, VIM depletion virtually abolished the PLA signals for ANO6-VIM and VIM-VAPB complexes (Fig. 6C). More importantly, the interaction dots for the canonical MAMs tethering pair VAPB-PTPIP51 were also depleted down to basal levels following VIM knockdown (Fig. 6C; P < 0.001, ****P < 0.0001). This demonstrates that the intermediate filament pillar is essential to mechanically stabilize the broader organelle tethering network.

**Figure 6.**
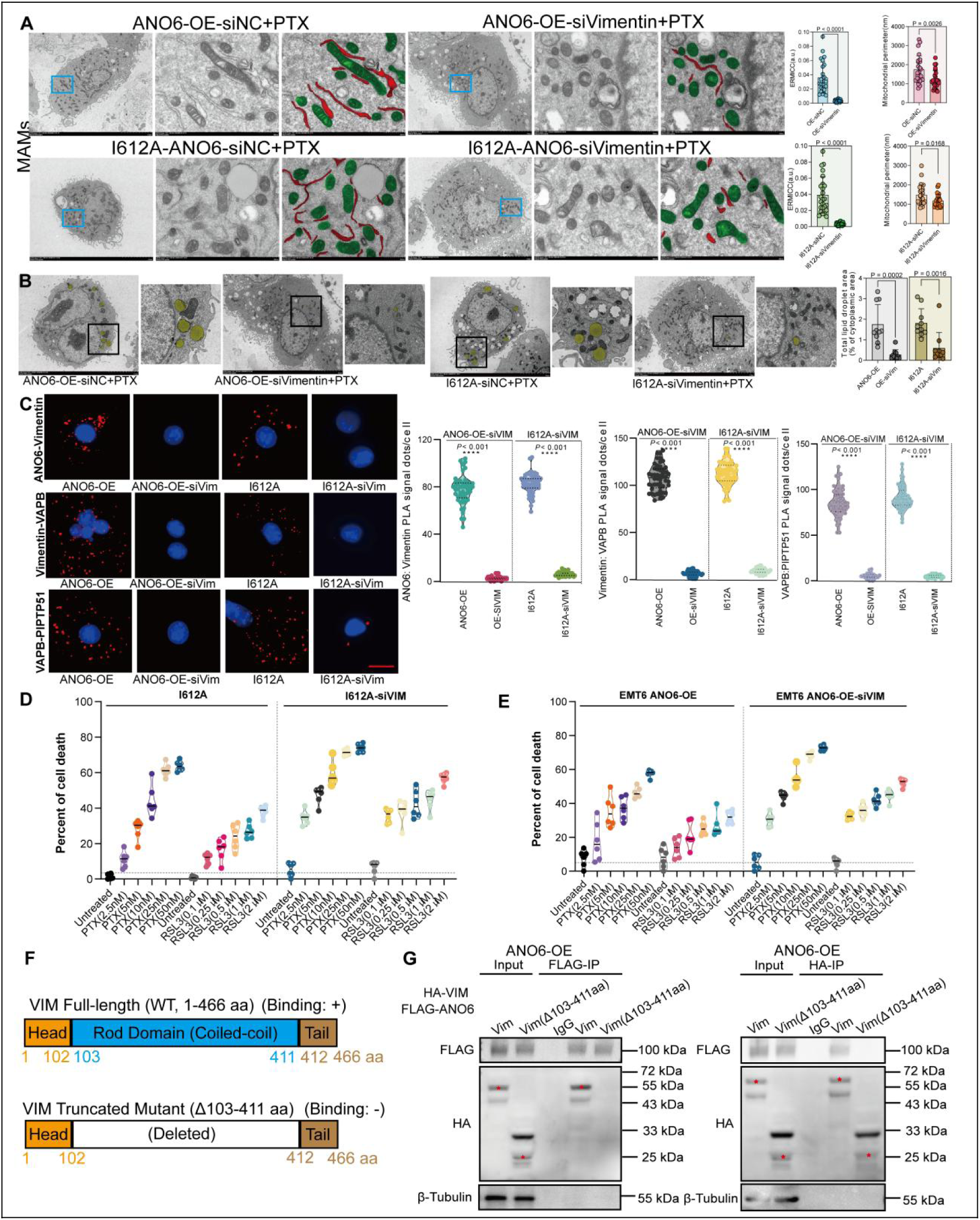
(A) Vimentin acts as an essential structural pillar for the ANO6-mediated MAMs tethering complex to drive lipid droplet packaging and chemoresistance.(A) Representative transmission electron microscopy (TEM) micrographs depicting mitochondria-associated endoplasmic reticulum membrane (MAMs) ultrastructure in ANO6-OE and I612A cells transfected with non-targeting control siRNA (siNC) or siRNA targeting Vimentin (siVimentin) in the presence of paclitaxel (5 nM PTX). ER and outer mitochondrial membranes within magnified insets (blue boxes) are pseudocolored in red and green, respectively (scale bars indicated in micrographs). Right: Quantitative evaluation of the ER-mitochondria contact coefficient (ERMICC; P < 0.0001) and mitochondrial perimeter (P = 0.0026 for ANO6-OE; P = 0.0168 for I612A by two-tailed unpaired Student’s t-test). (B) Representative TEM images showing intracellular lipid droplets (yellow pseudocolor; black boxes indicate magnified areas) and quantitative violin plots of total lipid droplet area relative to cytoplasmic area in ANO6-OE and I612A cells transfected with siNC or siVimentin following PTX treatment (P = 0.0032 for ANO6-OE; P = 0.0216 for I612A by two-tailed unpaired Student’s t-test). (C) Representative in situ proximity ligation assay (PLA) confocal micrographs (left; scale bar = 25 μm) and single-cell puncta quantifications (right) evaluating direct spatial proximity (<40 nm) of ANO6-Vimentin, Vimentin-VAPB, and VAPB-PTPIP51 in ANO6-OE and I612A cells upon VIM knockdown. Red puncta denote PLA interaction complexes; nuclei were counterstained with DAPI (blue) (P < 0.001, ****P < 0.0001 by one-way ANOVA). (D, E) Percentages of cell death in I612A (D) and ANO6-OE cells (E) transfected with siNC or siVimentin following treatment with escalating concentrations of paclitaxel (PTX, 2.5-50 nM) or the GPX4 inhibitor RSL3 (0.1-2 μM). (F) Schematic domain organization of full-length Vimentin (WT, 1-466 aa) and the central coiled-coil rod-domain deletion mutant VIM(Δ103-411aa). (G) Reciprocal co-immunoprecipitation (Co-IP) assays in ANO6-OE cells co-transfected with FLAG-tagged ANO6 and HA-tagged full-length VIM or HA-Vim(Δ103-411aa). Immunoprecipitates pulled down with anti-FLAG (left) or anti-HA (right) antibodies were blotted with the indicated antibodies; normal IgG served as the negative control and β-tubulin as the input loading control. Data are presented as mean ± SD from at least three independent biological replicates. *P < 0.05, **P < 0.01, ***P < 0.001, ****P < 0.0001; ns, not significant.

Functionally, we assessed whether disrupting this structural pillar resensitizes resistant cells to cytotoxic and ferroptotic lethality. Dose-escalation cell death assays demonstrated that VIM silencing markedly resensitized I612A (Fig. 6D) and ANO6-OE cells (Fig. 6E) to paclitaxel (2.5-50 nM) as well as to the ferroptosis inducer RSL3 (0.1-2 μM). Across multiple drug doses, the cell death rates of siVimentin cells approached 70-80%, abolishing the survival advantage conferred by ANO6 overexpression or the active I612A conformation (Fig. 6D, E).

To map the exact biochemical binding interface between ANO6 and VIM, we constructed a truncation mutant of Vimentin lacking its central coiled-coil rod domain, HA-Vim(Δ103-411aa) (Fig. 6F). Reciprocal co-immunoprecipitation (Co-IP) using anti-FLAG (ANO6) and anti-HA (VIM) antibodies in ANO6-OE cells revealed that full-length VIM strongly co-precipitated with ANO6, whereas deletion of the 103-411 aa rod domain completely abolished the physical interaction between VIM and ANO6 in both pull-down directions (Fig. 6G).

Together, these genetic and biochemical experiments establish that the central rod domain of VIM serves as an essential structural backbone that directly binds ANO6 to assemble MAMs contacts; breaking this pillar collapses the sequestration compartment and restores ferroptotic execution in chemoresistant breast cancer cells.

### Pharmacological disruption of the ANO6-MAMs structural axis dismantles lipid sequestration and unleashes ferroptotic death in chemoresistant tumors

Given that the structural complex assembled by ANO6, VIM, and VAPB constitutes the essential backbone maintaining the MAMs lipid sequestration compartment, we hypothesized that pharmacological targeting of this architecture would liberate trapped pro-ferroptotic PUFA-PLs, thereby resensitizing chemoresistant cells to ferroptosis and cytotoxic therapy. Because niclosamide (NIC) binds to an intramembrane pocket and locks ANO6 in an inactive, closed conformation, we hypothesized that this conformational lock allosterically disrupts its cytoplasmic docking surface and dismantles the MAMs tethering platform. In situ proximity ligation assays (PLA) revealed that in both ANO6-OE and hyper-scaffolding I612A cells, niclosamide treatment rapidly disassembled the spatial interactions of ANO6-Vimentin, Vimentin-VAPB, and VAPB-PTPIP51, reducing positive PLA signal dots down to basal levels (Fig. 7A; P < 0.001, ***P < 0.0001). This demonstrates that niclosamide physically dissociates the ANO6-nucleated organellar tethering platform at the single-cell level.

**Figure 7.**
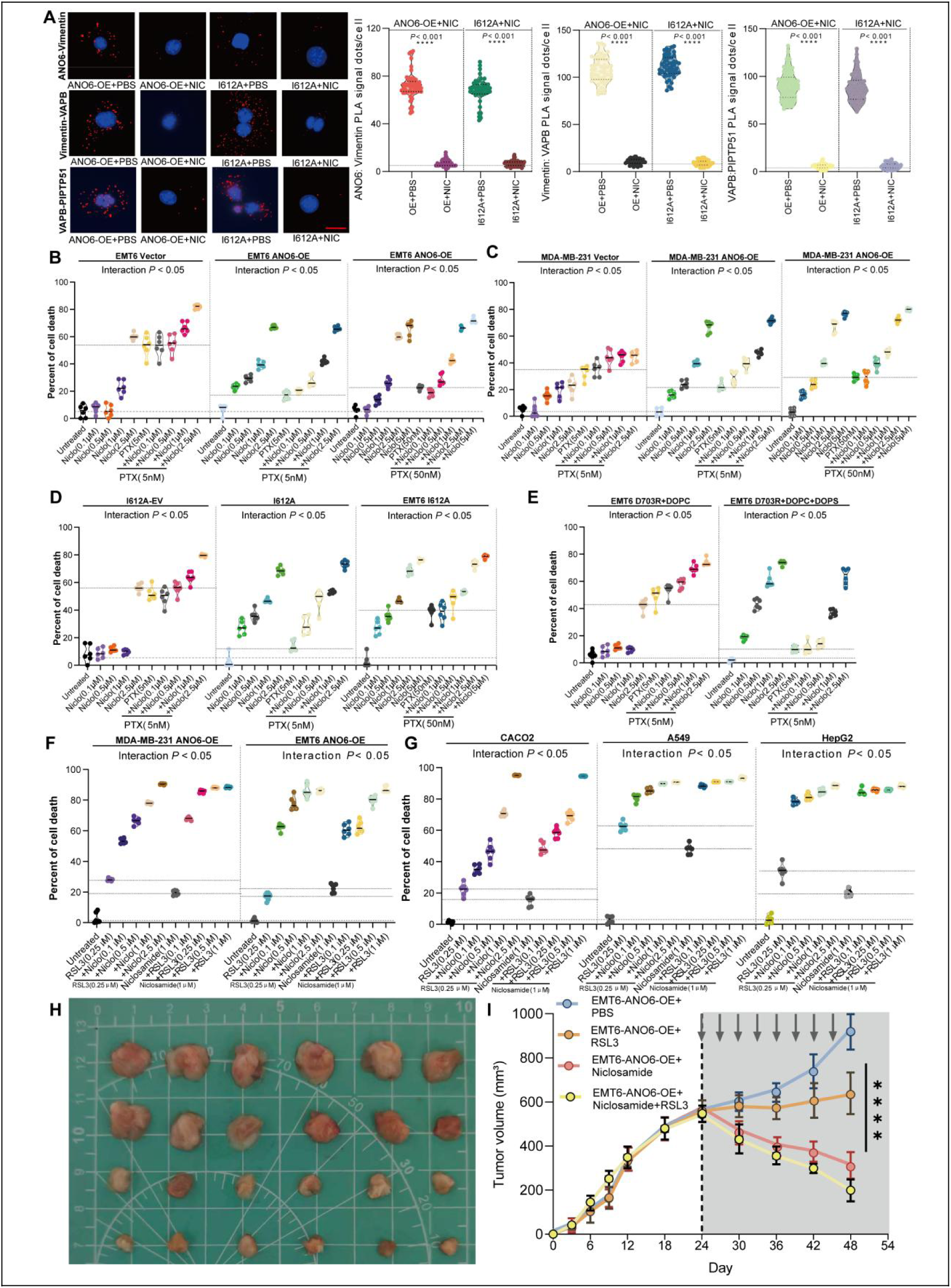
Pharmacological disruption of the ANO6-MAMs axis collapses lipid sequestration and synergistically triggers ferroptotic demise in chemoresistant tumors. (A) Representative in situ proximity ligation assay (PLA) confocal micrographs (left; scale bar = 25 μm) and single-cell puncta quantifications (right) evaluating direct spatial proximity (<40 nm) of ANO6-Vimentin, Vimentin-VAPB, and VAPB-PTPIP51 in ANO6-OE and I612A cells treated with vehicle (PBS) or niclosamide (NIC, 1 μM). Red dots indicate PLA complexes; cell nuclei were counterstained with DAPI (blue) (P < 0.0001 by two-tailed unpaired Student’s t-test). (B) Percentage of cell death in mouse EMT6 Vector vs. EMT6 ANO6-OE cells exposed to escalating doses of niclosamide (0.1-2.5 μM or 0.1-5 μM) alone or in combination with standard (5 nM) or escalated (50 nM) paclitaxel (PTX) challenge. Statistical synergy was determined by two-way ANOVA (interaction P < 0.05). (C) Percentage of cell death in human MDA-MB-231 Vector vs. MDA-MB-231 ANO6-OE cells treated with dose gradients of niclosamide alone or in combination with 5 nM or 50 nM PTX (interaction P < 0.05). (D) Comparative cell death responses evaluating synergistic interaction between PTX (5 nM or 50 nM) and niclosamide in I612A-EV vs. active scaffolding I612A cells (interaction P < 0.05). (E) Cell death percentages of EMT6 D703R cells pre-incubated with neutral liposomes (DOPC alone) or serine-containing liposomes (DOPC + DOPS) followed by challenge with PTX (5 nM) and increasing concentrations of niclosamide (interaction P < 0.05). (F) Synergistic induction of cell death in MDA-MB-231 ANO6-OE and EMT6 ANO6-OE cells co-treated with niclosamide (1 μM) and escalating doses of the GPX4 inhibitor RSL3 (0.25-1 μM; interaction P < 0.05). (G) Cross-cancer validation of synergistic cell death in CACO2 (colorectal), A549 (lung), and HepG2 (hepatocellular) carcinoma cells treated with niclosamide and RSL3 combinations (interaction P < 0.05). (H) Gross morphological photographs of orthotopic xenograft tumors excised at the study endpoint (Day 54) from mice bearing established EMT6 ANO6-OE tumors across four treatment cohorts: PBS control, single-agent RSL3, single-agent niclosamide, and combination regimen (n = 6 mice per group; scale in centimeters). (I) Longitudinal in vivo tumor growth trajectories monitored over 54 days (n = 6 mice per group). Gray vertical arrows indicate therapeutic administration days initiated on day 24 post-inoculation. Statistical differences among cohorts were evaluated by two-way repeated-measures ANOVA (P < 0.0001). Data are presented as mean ± SD from at least three independent experiments. *P < 0.05, **P < 0.01, ***P < 0.001, ****P < 0.0001; ns, not significant.

We next evaluated whether collapsing this sequestration depot restores therapeutic vulnerability. In mouse mammary carcinoma EMT6 cells, monotherapy with dose gradients of niclosamide elicited limited cytotoxicity; however, when combined with paclitaxel (PTX, 5 nM), it robustly abolished the survival advantage conferred by ANO6 overexpression, triggering 70%-80% synergistic cell death (Fig. 7B; two-way ANOVA, interaction P < 0.05). Notably, to account for the heightened chemoresistance threshold in ANO6-overexpressing cells, an escalated challenge (50 nM PTX) was applied alongside standard dosing (5 nM PTX); under both calibrated pressures, niclosamide co-treatment effectively overcame chemotherapy resistance (Fig. 7B; interaction P < 0.05). Consistent results were observed in human triple-negative breast cancer MDA-MB-231 models, where niclosamide synergized with both standard (5 nM) and high-dose (50 nM) PTX to trigger marked cell death in ANO6-OE cells (Fig. 7C; interaction P < 0.05). In structural mutant lines, cells expressing the active scaffolding mutant I612A displayed resistance to standard 5 nM PTX, but were profoundly resensitized by niclosamide even under escalated 50 nM PTX pressure (Fig. 7D; interaction P < 0.05). Conversely, in scaffold-compromised mutant D703R cells, although exogenous replenishment of serine-containing liposomes (DOPC + DOPS) transiently mitigated baseline PTX cytotoxicity compared to neutral DOPC alone, co-treatment with niclosamide completely overrode this protection and triggered pronounced cell death (Fig. 7E; interaction P < 0.05). Notably, in scramblase-compromised mutant D703R cells pre-incubated with DOPS liposomes to replenish surface PS signals, niclosamide co-treatment still completely overcame this baseline protection and triggered robust cell death, demonstrating that its synthetic lethality is primarily driven by dismantling the physical contact architecture rather than by inhibiting phospholipid scrambling per se.

To directly substantiate that this synergistic lethality is driven by the deliberate release of compartmentalized lipids and triggering of ferroptotic cascades, we co-treated cells with niclosamide and the GPX4 inhibitor RSL3. Combined exposure to niclosamide and escalating doses of RSL3 provoked massive, synergistic cell death in both MDA-MB-231 ANO6-OE and EMT6 ANO6-OE cells (Fig. 7F; two-way ANOVA, interaction P < 0.05). Crucially, this synthetic ferroptotic sensitivity extended beyond breast cancer. Based on expression profiling from The Human Protein Atlas demonstrating robust baseline abundance of ANO6 alongside its tethering partners VIM and VAPB across human carcinoma lines (Supplementary Fig. S12), we selected colorectal (CACO-2), non-small cell lung (A549), and hepatocellular carcinoma (HepG2) models for cross-cancer validation. Niclosamide co-treatment displayed consistent, potent synergistic killing with RSL3 across all three lineages (Fig. 7G; interaction P < 0.05), underscoring the broad pan-cancer therapeutic utility of disrupting MAMs-mediated lipid compartmentalization.

Finally, we translated this combination strategy in vivo using immunocompetent mice bearing orthotopic EMT6 ANO6-OE xenografts (Fig. 7H, I). Once tumors reached established burden (day 24), animals were randomized into four cohorts to receive vehicle (PBS), single-agent RSL3, single-agent niclosamide, or the concurrent combination regimen (Fig. 7I). While individual monotherapies exhibited only modest tumor retardation, the concurrent administration of niclosamide and RSL3 completely arrested chemoresistant tumor outgrowth and induced substantial therapeutic regression (reducing average volumes below 200 mm³), as corroborated by significant reductions in excised tumor dimensions and mass at the study endpoint (Fig. 7H, I; ***P < 0.0001).

Together, these pharmacological and in vivo findings establish that disrupting the ANO6-MAMs structural axis with niclosamide releases compartmentalized pro-ferroptotic PUFA-PLs, providing a rational combination strategy to eliminate chemoresistant malignancies via targeted ferroptosis execution.

## Discussion

Initiating from clinical cohorts of neoadjuvant chemotherapy-refractory breast cancers, this study delineates an organelle-level spatial paradigm by which chemoresistant cells manage perturbed lipid metabolism, integrating multi-omic profiling, organellar lipidomics, ultrastructural imaging, and conformational uncoupling assays. Aberrant lipid remodeling has increasingly been recognized as a central metabolic adaptation sustaining therapy persistence^[6, 20]^. However, prior investigations have largely focused on the altered expression of rate-limiting biosynthetic enzymes or total steady-state lipid abundances, paying insufficient attention to the spatial distribution of lipids among subcellular compartments^[12, 21]^. Here, we demonstrate that chemoresistant breast cancer cells utilize remodeled ER-mitochondria contact sites as a coordinated structural platform to route and package peroxidizable glycerophospholipids into neutral lipid droplets^[21–23]^. This organelle-level strategy provides a spatial dimension to cancer lipid biology, suggesting that the physical compartmentalization of altered lipids substantially influences cellular survival under cytotoxic chemotherapy^[22, 24, 25]^.

Our findings provide mechanistic insight into an intriguing metabolic paradox in chemoresistance. Under paclitaxel challenge, ANO6-overexpressing cells exhibit an upregulation of LPCAT2 alongside significant accumulation of polyunsaturated phospholipids (PUFA-PLs)^[26, 27]^. Under standard metabolic assumptions, an elevated burden of unsaturated fatty acyl chains would be expected to heighten sensitivity to cytotoxic oxidative stress, likely rendering cells more susceptible to ferroptotic cell death^[28, 29]^. Nevertheless, ANO6-proficient cells display sustained resistance in vitro and in vivo. MAMs microdomains represent primary hubs where phospholipid synthesis intersects with organellar metabolic fluxes^[23, 30, 31]^. Our organellar lipidomics and electron microscopy data indicate that resistant cells cope with this metabolic burden through spatial compartmentalization: newly generated PUFA-PLs are enriched within the MAMs microenvironment and subsequently packaged into the neutral cores of cytoplasmic lipid droplets^[21]^. By physically isolating these pro-oxidative lipid substrates away from functional membrane bilayers, resistant cells appear to buffer themselves against potential lipotoxic cascades^[21, 32]^.

This spatial lipid buffering likely operates in conjunction with adaptive biochemical adjustments. Under chemotherapy stress, cells expressing the active scaffolding mutant I612A exhibited coordinated adjustments in metabolic regulatory nodes, including induction of GPX4 and SLC7A11 alongside a repression of the fatty acid-activating enzyme ACSL4^[28, 33]^. These enzymatic shifts foster a favorable local antioxidant milieu to mitigate nascent oxidative signals at contact zones. However, the phenotypic collapse of the D703R mutant indicates that structural scaffolding serves as a major determinant of cell survival. Despite undergoing basal stress responses, D703R cells fail to repress ACSL4 and are structurally impaired in stabilizing MAMs or driving lipid droplet packaging^[21, 23]^. As a consequence, unsequestered lipid species may overwhelm cellular defenses, leaving these cells acutely vulnerable to cytotoxic stress and secondary lipid peroxidation^[34]^. This distinction supports the model that biochemical adaptations rely on an intact organelle architecture to provide effective metabolic protection^[23, 31]^.

At the molecular level, this study expands our understanding of TMEM16F/ANO6 biology by demonstrating that its established gating states simultaneously dictate its macromolecular scaffolding capacity at intracellular contact sites^[14, 35]^. Structural studies showed that I612 acts as an inner steric gate whose mutation (I612A) locks the pore in an open conformation, whereas D703 coordinates the primary Ca^2+^-binding pocket, the disruption of which (D703R) prevents calcium-dependent activation^[19]^. Our findings demonstrate that the open/activated conformation (I612A) exposes an accessible structural interface that readily recruits VIM and VAPB to assemble MAMs^[36]^, whereas the closed/calcium-deficient conformation (D703R) severely impairs inter-protein assembly (Fig. 3E, 4G). The faint residual Co-IP band and the modest, residual increase in ERMICC observed for D703R over empty vector (P = 0.0057; Fig. 3B, E) reflect basal low-affinity docking preserved on the folded cytosolic loops. However, this residual binding is physiologically insufficient to support high-order, functionally intact organelle tethering in intact cells, as corroborated by the virtual collapse of in situ PLA puncta (Fig. 4G, H) and the failure to promote protective lipid droplet sequestration (Fig. 4D, E). Furthermore, reciprocal Co-IP and deletion mapping identified the central coiled-coil rod domain of VIM (residues 103-411) as the indispensable docking interface that directly engages ANO6 (Fig. 6F, G). Targeted depletion of VIM or selective deletion of its rod domain severely dismantled MAMs architecture and abrogated lipid droplet accumulation, establishing that intermediate filaments actively provide mechanical and structural support to anchor organellar contact sites and govern intracellular lipid partitioning ^[36, 37]^.

Importantly, although restoring cell surface PS via DOPS liposomes transiently buffered baseline cytotoxicity in D703R cells, it failed to confer protection once MAMs tethering was pharmacologically dismantled by niclosamide (Fig. 7D). Niclosamide has been shown to allosterically lock TMEM16F in an inactive ground state^[38]^ ; here we show that this rigid lock distorts the cytosolic docking surface, disassembling the ANO6-VIM-VAPB platform and releasing compartmentalized PUFA-PLs to trigger synthetic lethal ferroptosis (Fig. 7A, D, G). Thus, the conformational gating of ANO6 operates as a unified molecular switch governing both lipid scrambling and organelle contact architecture, with the latter serving as the primary driver of spatial lipid compartmentalization and chemoresistance.

Elucidating this spatial lipid buffering mechanism reveals an actionable therapeutic vulnerability in chemoresistant malignancies. Given that resistant cells persist in part by structurally sequestering pro-oxidative lipid species, pharmacological disruption of this tethering architecture presents an attractive strategy to release compartmentalized lipids and expose hidden metabolic liabilities^[22]^. We show that treatment with niclosamide rapidly dissociates the ANO6-VIM-VAPB platform, collapsing MAMs contacts and destabilizing lipid packaging^[38, 39]^. This structural collapse prompts the unregulated distribution of sequestered PUFA-PLs toward vulnerable membrane bilayers, markedly potentiating tumor sensitivity to secondary lipid stress and GPX4 inhibition across multiple carcinoma models in vitro and in vivo^[28, 38]^. Combining organelle contact disruptors with ferroptosis-inducing agents or standard chemotherapy represents a rational approach to exploit lipid metabolic vulnerabilities and target persistent tumor populations^[28, 38]^.

In summary, our findings uncover a spatial paradigm wherein chemoresistant breast cancer cells exploit ANO6-mediated contact scaffolding to partition aberrant glycerophospholipids into protective intracellular compartments^[21]^. Structurally disrupting this organellar tethering platform abolishes lipid buffering, providing a conceptual framework to exploit metabolic lipid liabilities and overcome chemotherapy persistence in refractory cancer^[38]^.

## Declarations

### Ethics Approval and Consent to Participate

This study was approved by the Institutional Ethics Committee of the Third Affiliated Hospital of Kunming Medical University (Approval No. KYCS2023-056). Written informed consent was obtained from all participating patients prior to tissue collection, in strict compliance with the Declaration of Helsinki, the ethical guidelines of the Council for International Organizations of Medical Sciences (CIOMS), and relevant international and institutional regulations.

All animal experiments were formally approved by the Animal Experimental Ethics Committee of Kunming Medical University (Protocol No. kmmu20230811 / KMMU20230811) and conducted in rigorous accordance with institutional and national guidelines for laboratory animal welfare, as well as the ARRIVE guidelines (Animal Research: Reporting of In Vivo Experiments; https://arriveguidelines.org). Established human and mouse cell lines used in this work complied with relevant institutional guidelines and did not require separate ethical permits.

### Consent for Publication

Informed consent for the publication of clinical baseline characteristics, histopathological micrographs, and molecular profiling data was obtained from all enrolled patients or their legally authorized representatives. De-identified patient details were used throughout the manuscript to protect patient confidentiality.

### Availability of Data and Materials

All data supporting the findings of this study are available within the article and its supplementary files. The newly generated high-throughput sequencing datasets (including transcriptomic and multi-omic profiling data of clinical specimens and engineered models), raw untargeted/targeted lipidomic and metabolomic mass spectrometry data, uncropped immunoblot images, and plasmid/construct sequence files generated during the current study are available from the corresponding author upon reasonable request. Custom bioinformatic analysis scripts used in this study are also available from the corresponding author upon request.

### Authors’ Contributions

Conception and Supervision: QLY, JY and QXY, Experimental Design: QLY, JY, and QXY; Data Acquisition and In Vivo/In Vitro Assays: QLY, QXY, JY, PZ, QT, KZ, JWZ and ZTZ; Bioinformatics and Multi-Omics Data Analysis: QXY, QLY and JY, Data Interpretation and Mechanistic Dissection: QLY, QXY, JY, PZ, QT, KZ, JWZ and ZTZ; Figure Preparation and Visualization: QLY, JY and QXY, manuscript draft: QLY, JY, and QXY, manuscript editing: QLY, QXY, JY, PZ, QT, KZ, JWZ and ZTZ. All authors have approved the submitted version.

## Acknowledgements

We sincerely thank all the patients who generously contributed their clinical specimens to make this investigation possible. We also gratefully acknowledge the core biomedical imaging, high-throughput sequencing, and mass spectrometry platforms at Kunming Medical University for their valuable technical assistance and infrastructural support.

## Clinical Trial Number

Not applicable.

## Supplementary Figure

**Supplementary Figure S1.**
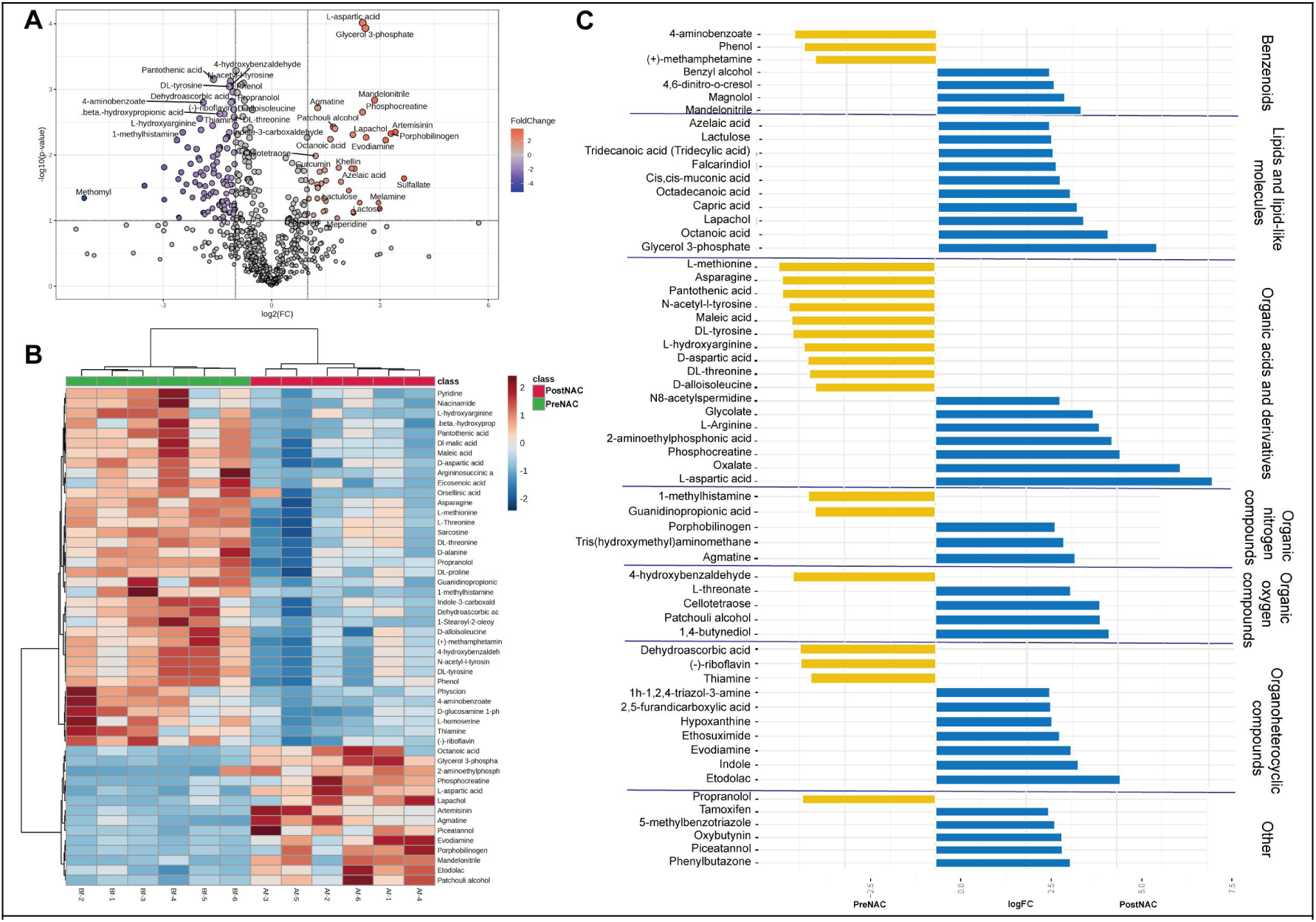
Untargeted metabolomic landscape of paired breast cancer clinical specimens before and after neoadjuvant chemotherapy. (A) Volcano plot illustrating differentially abundant metabolites between pre-NAC and post-NAC residual breast cancer lesions (n = 6 paired patient specimens). The horizontal dashed line denotes the statistical significance threshold (-log_10_(P-value) = 1.3, corresponding to P = 0.05 by two-tailed Student’s t-test); vertical dashed lines denote fold-change thresholds (log_2_FC ≥ 0.58, corresponding to a 1.5-fold difference). Data points are colored according to fold-change gradient (blue: downregulated; red: upregulated in post-NAC residual tissues; gray circles: non-significant). Prominently dysregulated metabolites, including glycerol 3-phosphate, L-aspartic acid, and dehydroascorbic acid, are highlighted. (B) Hierarchical clustering heatmap of significantly altered metabolites across paired patient biopsies. The top column annotations indicate matched clinical cohorts (green: PreNAC; red: PostNAC). Row-normalized relative abundances are expressed as Z-scores, colored along a continuous gradient from blue (low relative abundance) to red (high relative abundance). (C) Class-stratified horizontal bar plot displaying the log_2_FC of representative altered metabolites organized by major biochemical superclasses (Benzenoids, Lipids and lipid-like molecules, Organic acids and derivatives, Organic nitrogen compounds, Organic oxygen compounds, Organoheterocyclic compounds, and Other). Blue bars denote metabolites upregulated in post-chemotherapy residual lesions (PostNAC, log_2_FC > 0, highlighted by pronounced accumulation of glycerol 3-phosphate and octanoic acid), and yellow bars denote metabolites enriched in chemotherapy-naive biopsies (PreNAC, log_2_FC < 0, such as 4-aminobenzoate and L-methionine).

**Supplementary Figure S2.**
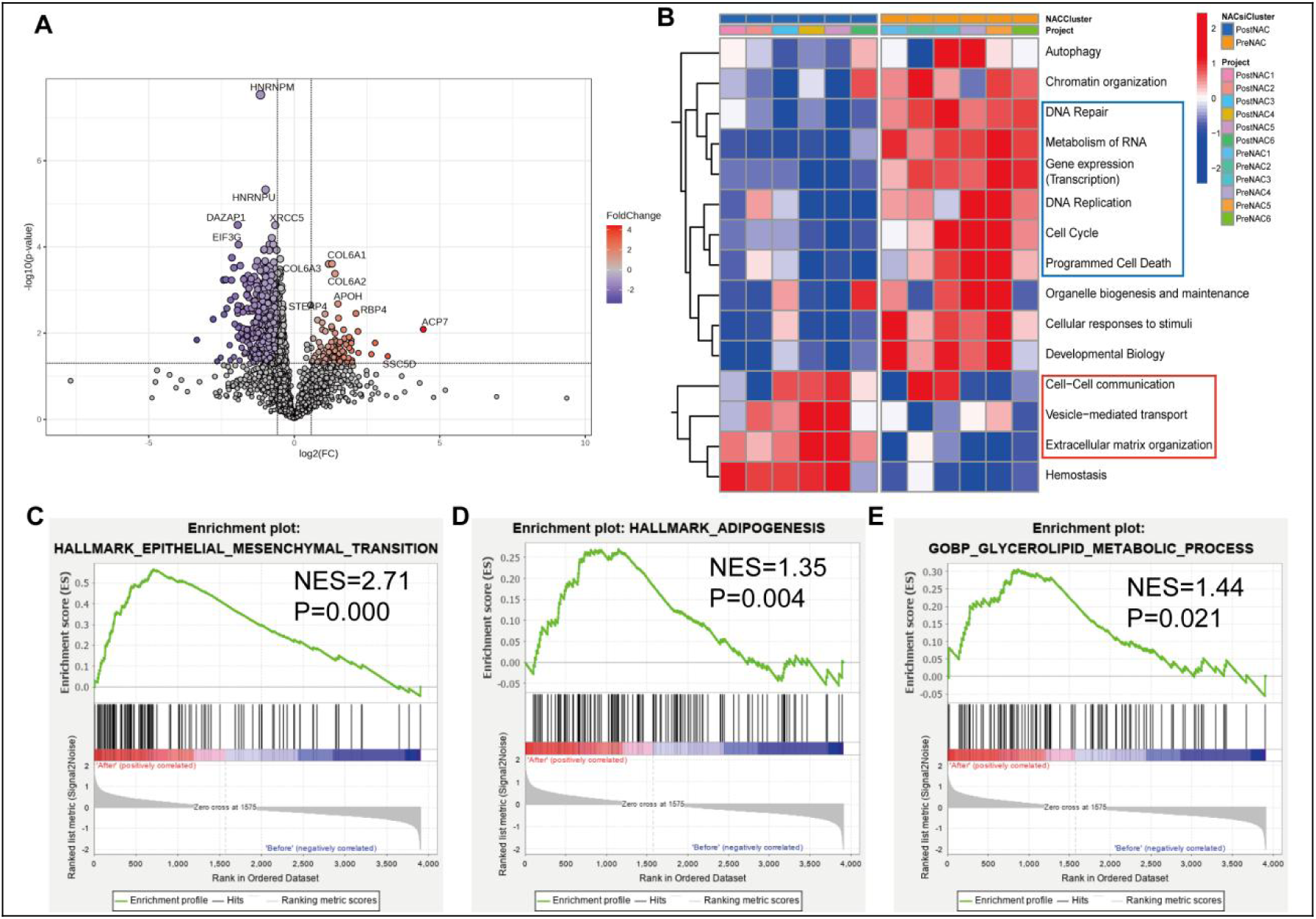
Proteomic landscape and pathway enrichment analysis of paired clinical breast cancer specimens before and after neoadjuvant chemotherapy. (A) Volcano plot displaying differentially expressed proteins between pre-NAC baseline biopsies (PreNAC) and post-NAC chemoresistant residual breast cancer tissues (PostNAC) (n = 6 paired patient specimens). The horizontal dashed line denotes the statistical significance cutoff (-log_10_(P-value) = 1.3, equivalent to P = 0.05 by two-tailed Student’s t-test); vertical dashed lines represent fold-change thresholds (log_2_FC ≥ 0.58, equivalent to a 1.5-fold difference). Data points are colored according to fold-change gradient (blue: downregulated; red: upregulated in PostNAC residual tissues; gray circles: non-significant). Prominently upregulated proteins include ACP7, RBP4, COL6A1, COL6A2, COL6A3, STEAP4, APOH, and SSC5D; prominently downregulated proteins include HNRNPM, HNRNPU, XRCC5, DAZAP1, and EIF3G. (B) Hierarchical clustering heatmap showing the relative activity of core biological process signatures across paired clinical specimens (left columns: PreNAC 1-6; right columns: PostNAC 1-6, annotated by top color bars). Color scale reflects row-normalized Z-scores from low (blue) to high (red) pathway enrichment. The blue bounding box highlights biological processes prominently elevated in post-chemotherapy residual lesions (including DNA repair, metabolism of RNA, gene expression/transcription, DNA replication, cell cycle, and programmed cell death); the red bounding box marks stromal and communication signatures exhibiting higher relative enrichment in treatment-naive baseline biopsies (cell-cell communication, vesicle-mediated transport, and extracellular matrix organization). (C-E) Gene Set Enrichment Analysis (GSEA) plots illustrating key oncogenic and metabolic pathways positively enriched in post-NAC chemoresistant residual tissues (PostNAC): (C) Epithelial-to-mesenchymal transition hallmark signature (HALLMARK_EPITHELIAL_MESENCHYMAL_TRANSITION; normalized enrichment score NES = 2.71, P < 0.001); (D) Adipogenesis hallmark signature (HALLMARK_ADIPOGENESIS; NES = 1.35, P = 0.004); (E) Glycerolipid metabolic process Gene Ontology pathway (GOBP_GLYCEROLIPID_METABOLIC_PROCESS; NES = 1.44, P = 0.021).

**Supplementary Figure S3.**
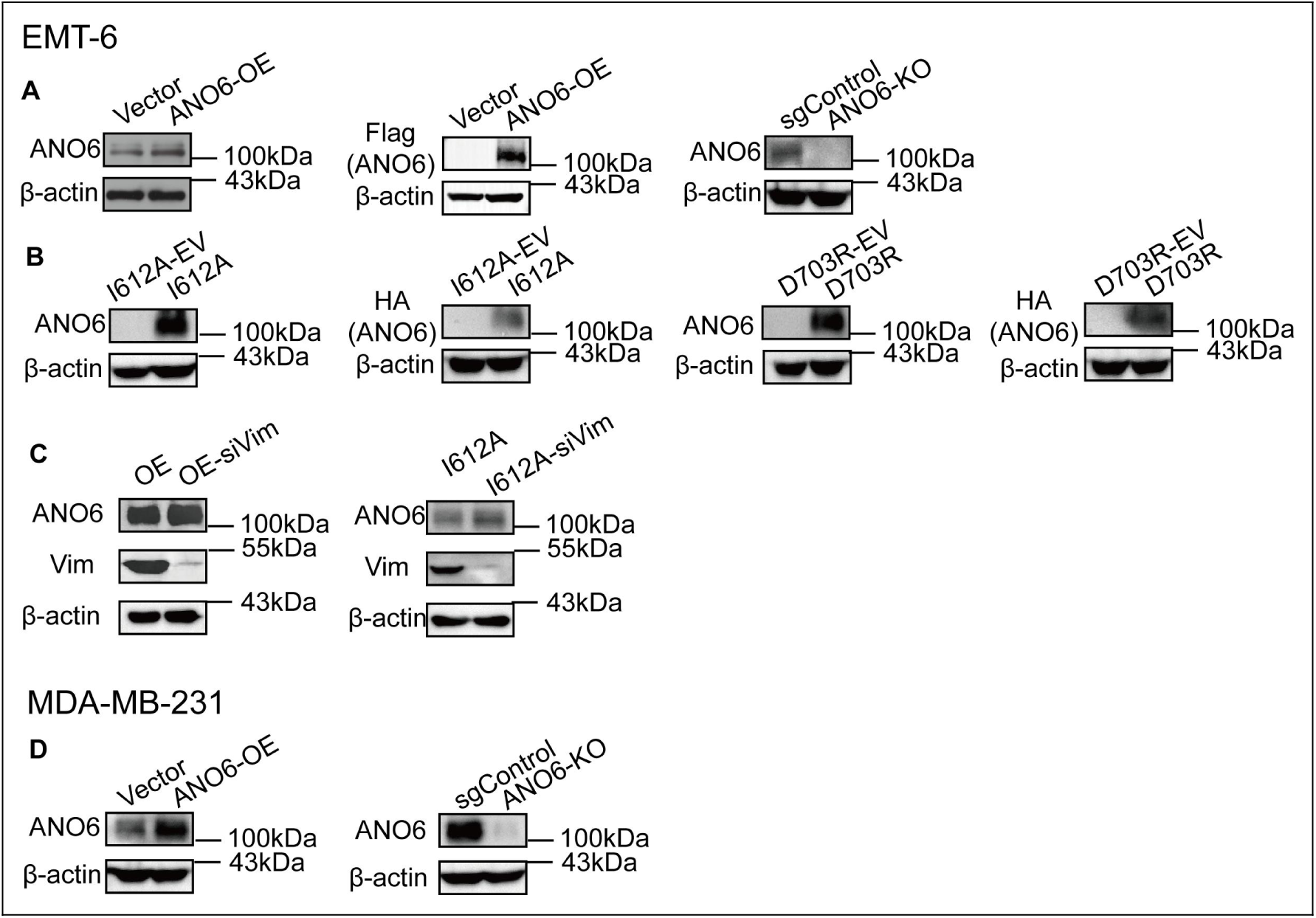
Western blot validation of engineered breast cancer cell models. Representative immunoblot analyses confirming the establishment, genetic manipulation, and reconstitution of experimental cell models in mouse EMT-6 and human MDA-MB-231 breast cancer backgrounds: (A-C) EMT-6 cell models:(A) ANO6 overexpression and knockout: Verification of ectopic ANO6 expression in Vector versus ANO6-OE cells probed with anti-ANO6 and anti-Flag tag antibodies (left and middle), alongside confirmation of CRISPR-Cas9-mediated gene disruption displaying complete loss of endogenous ANO6 protein in ANO6-KO cells relative to non-targeting sgControl cells (right). (B) Conformational mutant reconstitution: Immunoblot validation of stable reconstitution in engineered EMT-6 lines transduced with matched empty vectors or constructs encoding the active scaffolding mutant (I612A-EV vs. I612A) or the scaffolding-compromised mutant (D703R-EV vs. D703R), detected using anti-ANO6 and anti-HA tag antibodies. (C) Vimentin knockdown: Evaluation of siRNA-mediated silencing of Vimentin (siVim) in ANO6-OE (OE vs. OE-siVim) and I612A (I612A vs. I612A-siVim) backgrounds, confirming robust depletion of Vim protein (55 kDa) without altering ANO6 abundance (100 kDa). (D) MDA-MB-231 cell models:Immunoblot validation of ANO6 overexpression (Vector vs. ANO6-OE, left) and CRISPR-Cas9-mediated knockout (sgControl vs. ANO6-KO, right) in human MDA-MB-231 triple-negative breast cancer cells probed with an anti-ANO6 antibody. In all panels, β-actin (43 kDa) served as the internal loading control. Molecular weights are indicated in kilodaltons (kDa) on the right of each blot.

**Supplementary Figure S4.**
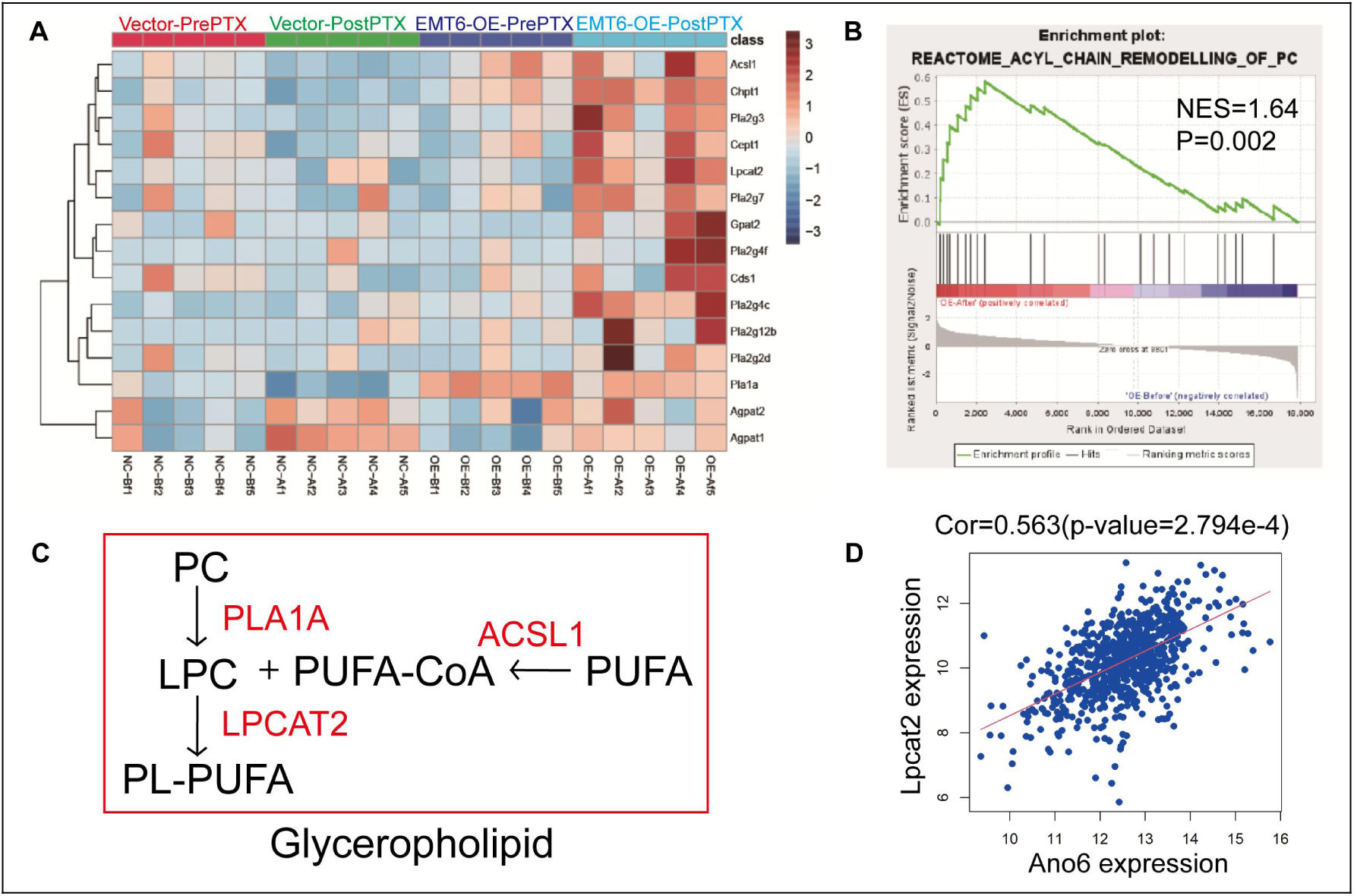
Activation of the Lands cycle and acyl-chain remodeling of phosphatidylcholine in ANO6-overexpressing tumors upon chemotherapy. (A) Hierarchical clustering heatmap displaying the relative expression levels of enzymes governing phospholipid metabolism and acyl-chain remodeling in xenograft tumors across experimental cohorts: Vector-PrePTX (red bar), Vector-PostPTX (green bar,), EMT6-OE-PrePTX (purple bar, ), and EMT6-OE-PostPTX (cyan bar) (n = 5 mice per cohort). Color scale denotes row-normalized Z-scores from low (blue) to high (dark red) expression. Key remodeling enzymes exhibiting prominent upregulation in paclitaxel-treated ANO6-OE lesions include Lpcat2, Acsl1, Pla2g7, and Gpat2. (B) Gene Set Enrichment Analysis (GSEA) plot demonstrating significant positive enrichment of the phosphatidylcholine acyl-chain remodeling pathway (REACTOME_ACYL_CHAIN_REMODELLING_OF_PC; NES = 1.64, P = 0.002) in EMT6-ANO6-OE tumors following paclitaxel challenge. (C) Schematic diagram depicting the Lands cycle remodeling cascade in glycerophospholipid metabolism: cleavage of phosphatidylcholine (PC) by phospholipase A1 (PLA1A) yields lysophosphatidylcholine (LPC), which is subsequently re-acylated with polyunsaturated fatty acyl-CoA (PUFA-CoA, synthesized from free PUFA by ACSL1) via LPCAT2 to generate PUFA-containing phospholipids (PL-PUFA) (rate-limiting catalytic enzymes highlighted in red). (D) Pearson correlation scatter plot demonstrating a significant positive correlation between Ano6 transcript abundance and Lpcat2 expression in breast cancer tissues (r = 0.563, P = 2.794e-4).

**Supplementary Figure S5.**
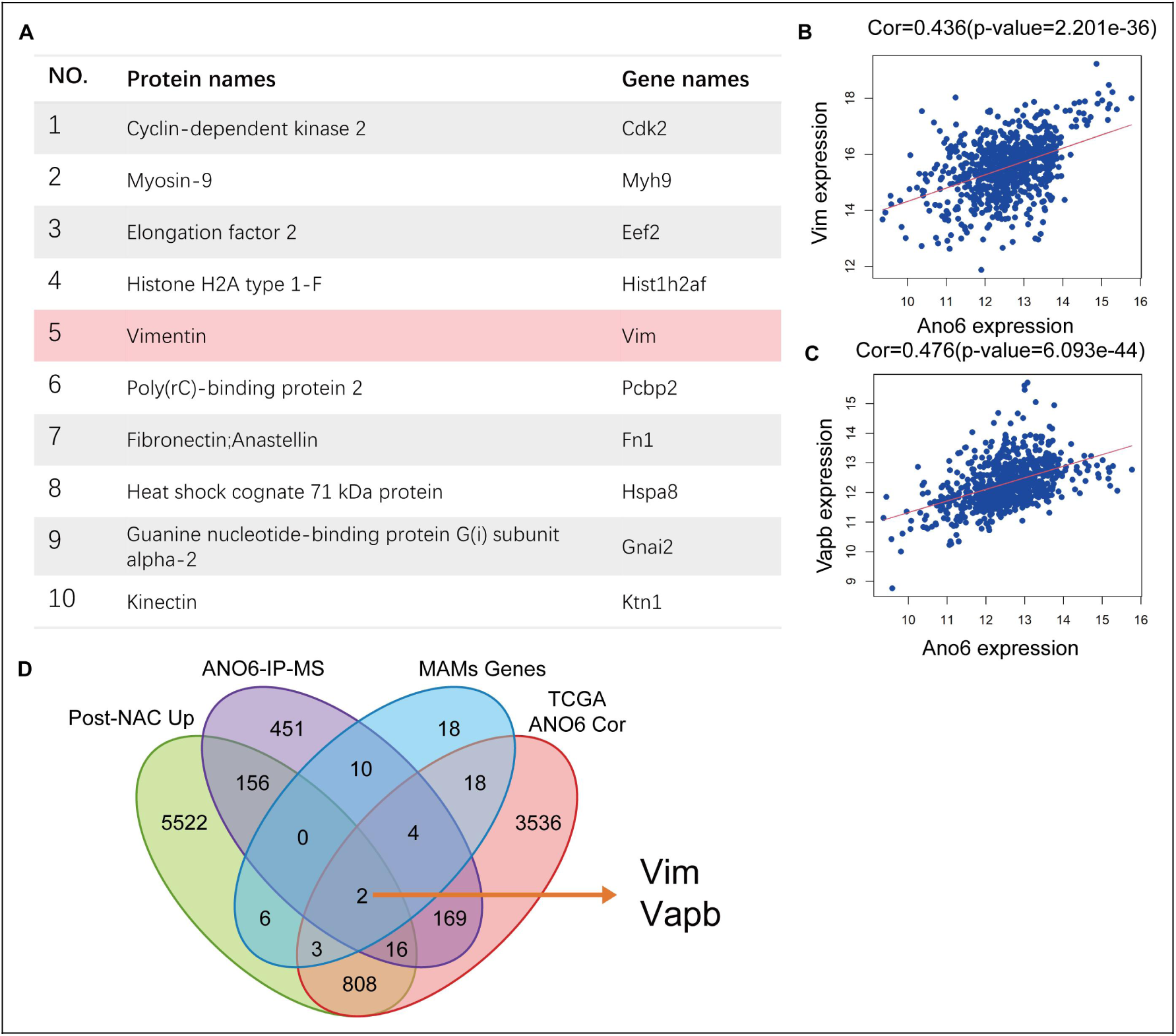
Integrated mass spectrometry and multi-cohort intersection identify Vimentin and VAPB as core ANO6-associated MAMs scaffolding partners. (A) Top 10 candidate interacting proteins identified by co-immunoprecipitation followed by liquid chromatography-tandem mass spectrometry (ANO6-IP-MS) in breast cancer cells. Vimentin (VIM/Vim) is highlighted with light pink shading in row 5. (B, C) Pearson correlation scatter plots demonstrating significant positive associations between Ano6 mRNA expression and Vim (r = 0.436, P = 2.201e-36; B) as well as Vapb (r = 0.476, P = 6.093e-44; C) across breast cancer patient cohorts. Red lines represent linear regression fits. (D) Four-way Venn diagram illustrating the multi-omic filtering strategy intersecting: (1) genes significantly upregulated in post-chemotherapy residual clinical specimens (Post-NAC Up), (2) physical binding candidates identified by ANO6-IP-MS, (3) curated mitochondria-associated ER membrane components (MAMs Genes), and (4) genes exhibiting positive expression correlation with ANO6 in the TCGA breast cancer cohort (TCGA ANO6 Cor). The intersection uniquely converged on two core scaffolding candidates (indicated by arrow): Vim and Vapb.

**Supplementary Figure S6.**
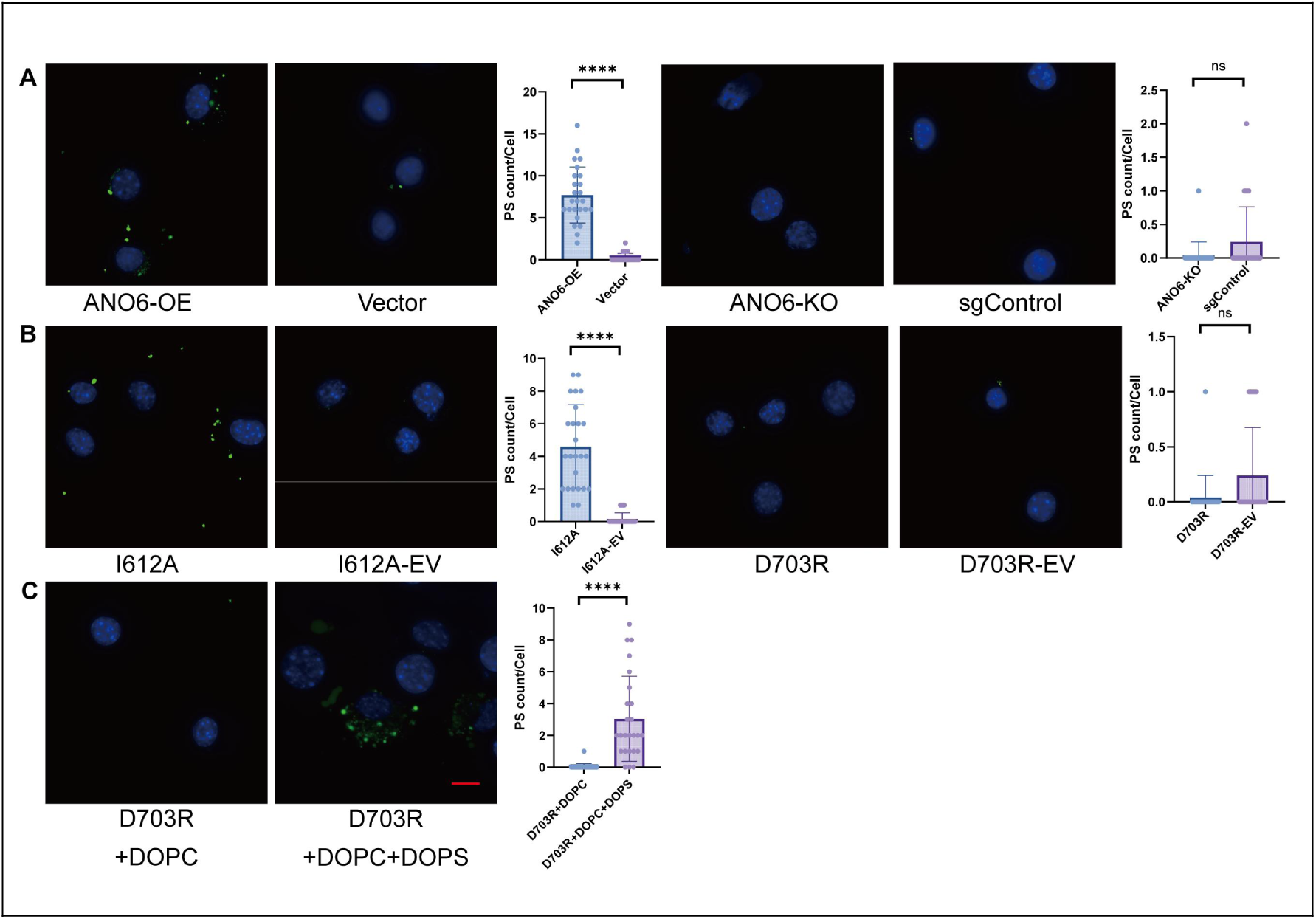
In situ fluorescent imaging of cell-surface phosphatidylserine exposure across engineered ANO6 cell models. exposure across engineered ANO6 cell models.Representative high-resolution live-cell confocal micrographs visualizing cell-surface phosphatidylserine (PS) externalization (green puncta; stained with lactadherin-FITC; scale bar = 25 μm) counterstained with Hoechst 33342 (nuclei, blue), alongside corresponding single-cell dot-plot quantifications of surface PS puncta (PS count/cell): (A) Assessment of surface PS puncta in wild-type ANO6 models, comparing ANO6-overexpressing cells with Vector controls (left; ****P < 0.0001 by two-tailed unpaired Student’s t-test), and ANO6-knockout (ANO6-KO) cells with sgControl cells (right; ns, not significant). (B) Evaluation of conformational mutants showing robust surface PS puncta accumulation in the active scaffolding mutant (I612A) relative to its empty vector control (I612A-EV; P < 0.0001), whereas the scaffolding-compromised mutant/catalytically dead mutant (D703R) displayed baseline-level signals indistinguishable from D703R-EV (right; ns). (C) Exogenous replenishment of serine-containing liposomes (DOPC + DOPS) on D703R cells selectively restores cell-surface PS puncta decoration relative to neutral liposomes (DOPC alone) (P < 0.0001). Data are presented as mean ± SD from three independent biological replicates. ****P < 0.0001; ns, not significant.

**Supplementary Figure S7.**
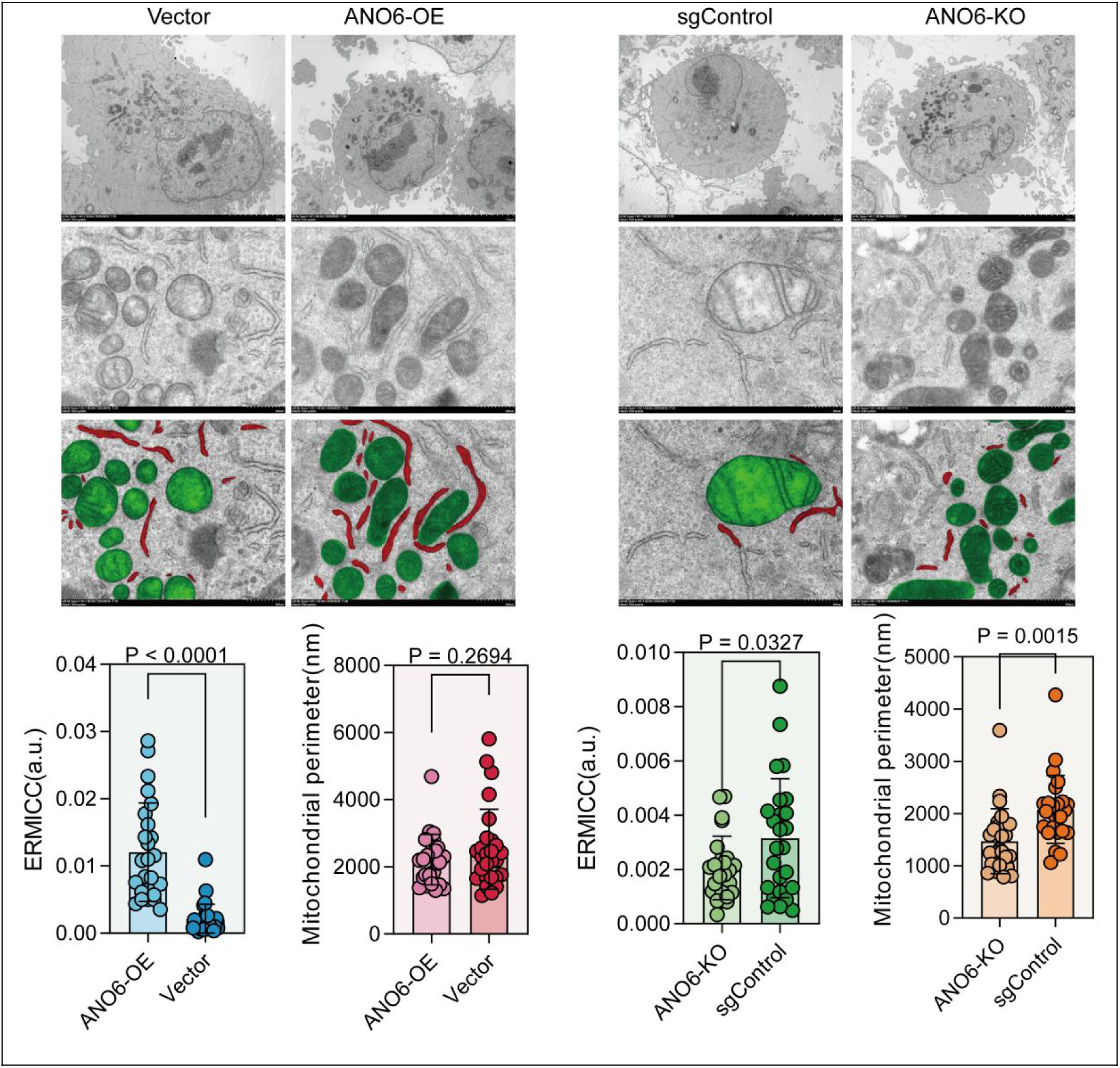
Transmission electron microscopy evaluates the modulation of ER-mitochondria contact ultrastructure by ANO6 abundance in MDA-MB-231. Representative transmission electron microscopy (TEM) micrographs and morphometric quantifications assessing mitochondria-associated endoplasmic reticulum membrane (MAMs) architecture in human MDA-MB-231 triple-negative breast cancer cells across comparative genetic cohorts: control (Vector) versus ANO6-overexpressing (ANO6-OE) cells, and non-targeting control (sgControl) versus ANO6-knockout (ANO6-KO) cells. Top and middle panels: Low-magnification whole-cell overviews (top row) and high-magnification micrographs capturing inter-organellar contact interfaces (middle row) (scale bars indicated in micrographs). In the third row, mitochondria are pseudocolored in green, and endoplasmic reticulum (ER) cisternae closely apposed to the outer mitochondrial membrane (≤60 nm cleft distance) are highlighted in red. Bottom plots (left): Scatter dot-plot quantifications of the ER-mitochondria contact coefficient (ERMICC in arbitrary units [a.u.], left plot; P < 0.0001) and mitochondrial perimeter (nm, right plot; P = 0.2694; ns, not significant) comparing ANO6-OE with Vector cells evaluated by two-tailed unpaired Student’s t-test. Bottom plots (right): Quantitative morphometry of ERMICC (left plot; P = 0.0327) and mitochondrial perimeter (right plot; P = 0.0015) comparing ANO6-KO with sgControl cells. Data are presented as mean ± SD from independent biological replicates. *P < 0.05, **P < 0.01, ****P < 0.0001; ns, not significant.

**Supplementary Figure S8.**
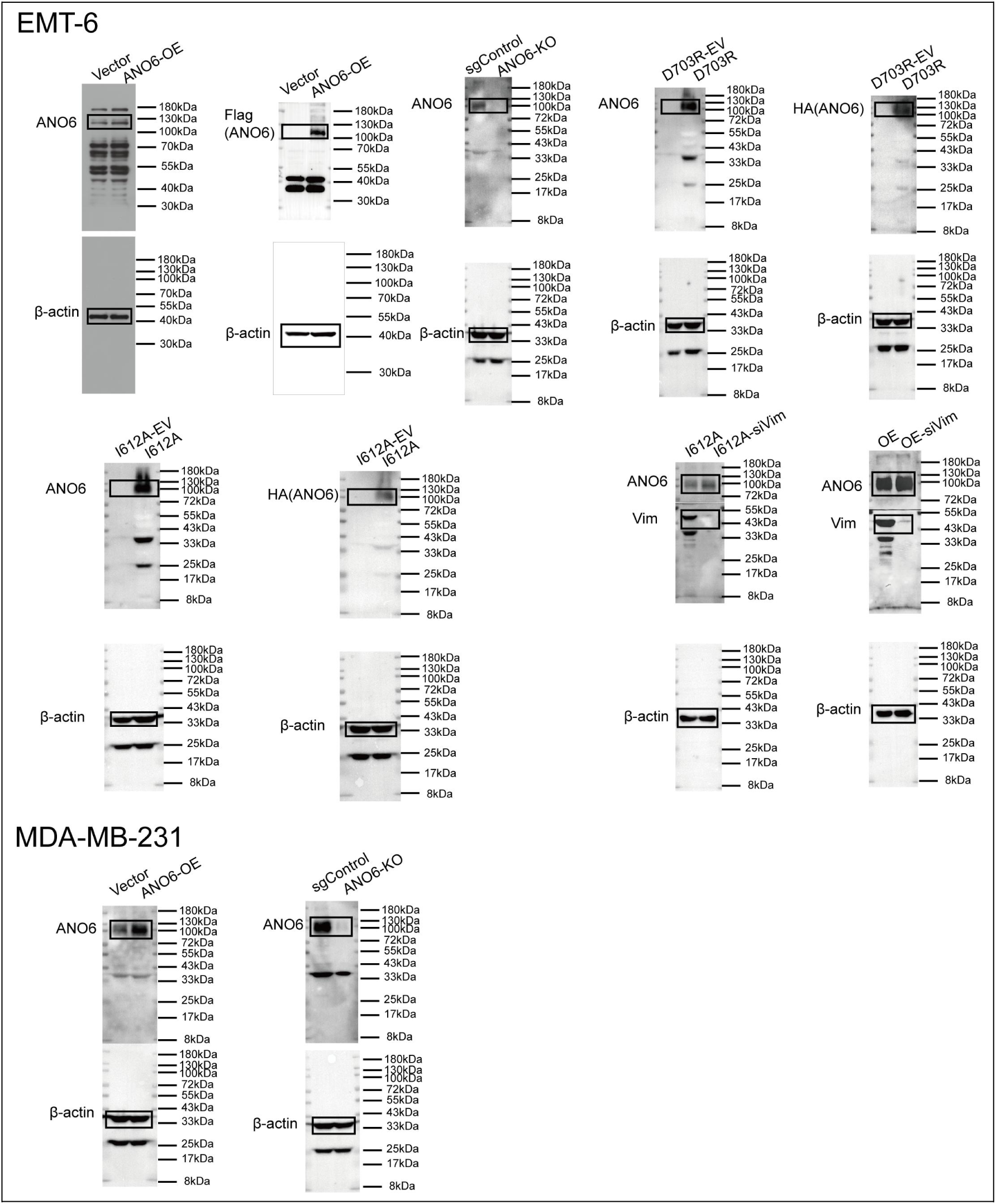
Full-length uncropped immunoblot images corresponding to Supplementary Figure S3. Uncropped Western blot membranes displaying target proteins and their corresponding loading controls (β-actin) used for the genetic validation of engineered mouse EMT-6 and human MDA-MB-231 breast cancer cell models. Black rectangular boxes denote the cropped regions presented in Supplementary Figure S3. Protein molecular weight ladders (in kDa) are indicated adjacent to each membrane.

**Supplementary Figure S9.**
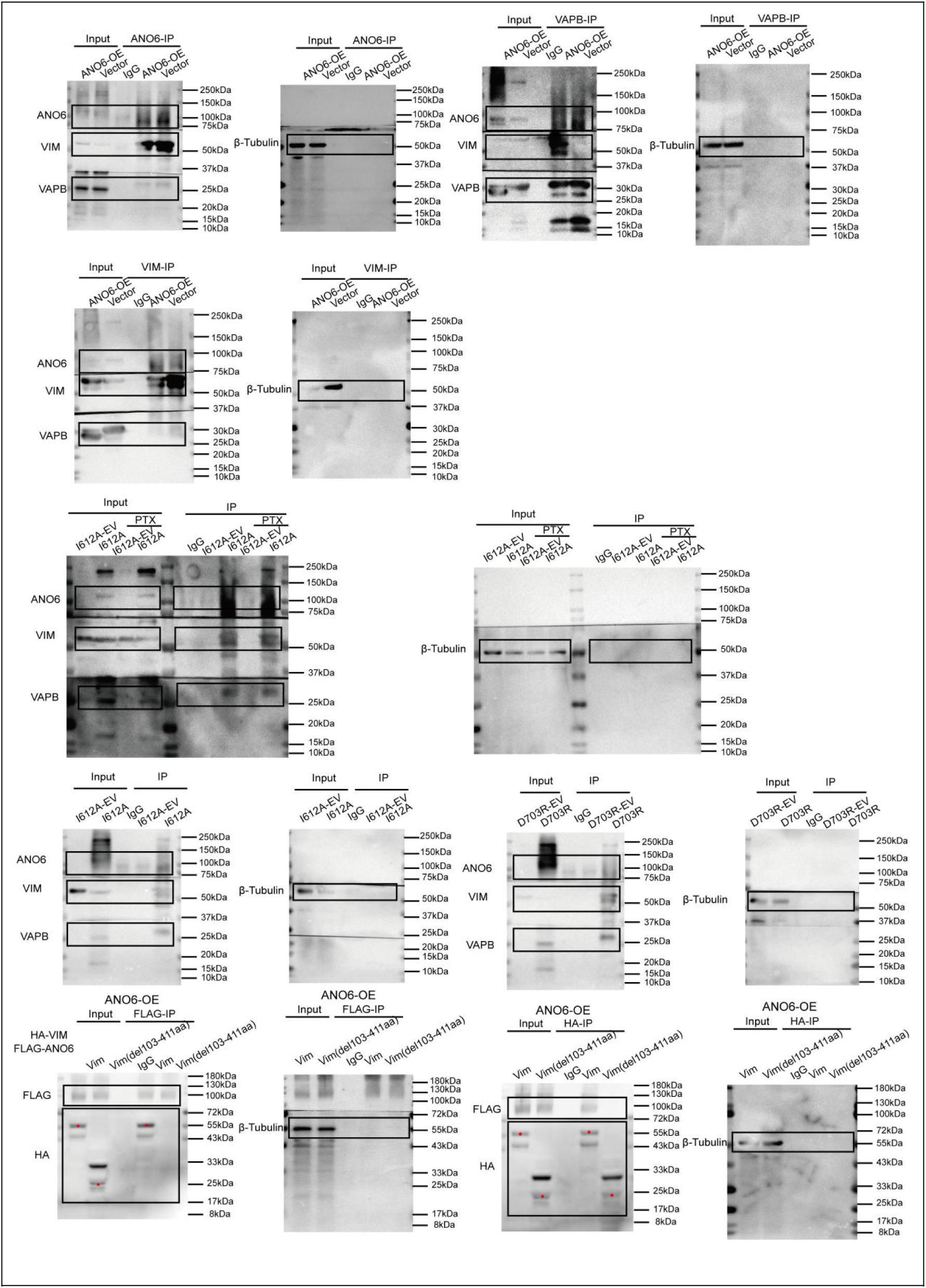
Full-length uncropped immunoblot images corresponding to Co-IP assays in Figure 3 and Figure 6. Uncropped Western blot membranes displaying target proteins and corresponding input loading controls (β-tubulin, 55 kDa) for all co-immunoprecipitation (Co-IP) experiments presented in the manuscript: Rows 1 and 2 (top): Uncropped membranes for reciprocal endogenous Co-IP assays evaluating mutual physical interactions among ANO6, Vimentin (VIM), and VAPB in Vector and ANO6-OE cells following anti-ANO6, anti-VAPB, or anti-VIM pull-down, corresponding to Figure 3C. Row 3 (middle): Uncropped membranes assessing physical interactions of the active scaffolding mutant I612A with VIM and VAPB in the presence or absence of paclitaxel (PTX) challenge, corresponding to Figure 3D. Row 4: Uncropped membranes evaluating physical binding capacities toward VIM and VAPB between the active scaffolding mutant I612A and the scaffold-compromised mutant D703R relative to their matched empty vector controls (I612A-EV and D703R-EV), corresponding to **Figure 3E. Row 5 (bottom): Uncropped membranes for reciprocal Co-IP assays mapping the interaction interface between FLAG-ANO6 and HA-tagged full-length VIM or the rod-domain deletion mutant HA-Vim(Δ103-411aa) following anti-FLAG or anti-HA pull-down, corresponding to Figure 6G. Black rectangular boxes delineate cropped blot areas displayed in main figures. Red asterisks mark target recombinant truncation bands. Protein molecular weight ladders (in kDa) are indicated adjacent to each membrane.

**Supplementary Figure S10.**
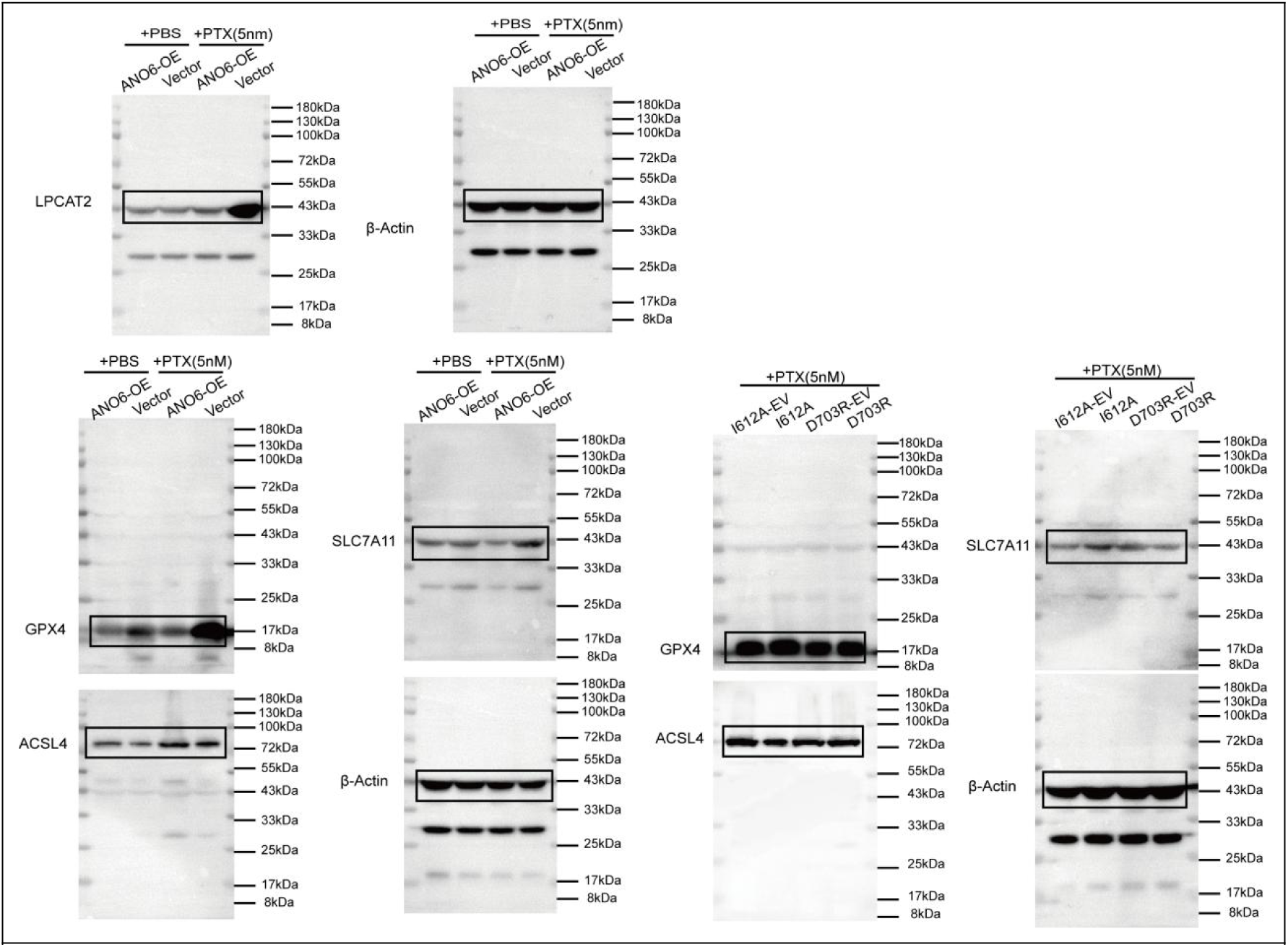
Full-length uncropped immunoblot images corresponding to Main Figure 2 and Figure 4. Uncropped Western blot membranes displaying target proteins and corresponding loading controls (β-actin, 43 kDa) for lipid-remodeling enzymes and canonical ferroptosis surveillance machinery: Top panels: Full-length membranes for LPCAT2 (43 kDa) and matched β-actin in ANO6-OE and Vector cells treated with vehicle (PBS) or paclitaxel (5 nM PTX), corresponding to Figure 2H. Bottom-left panels: Full-length membranes evaluating canonical ferroptosis surveillance proteins GPX4 (17 kDa), ACSL4 (72 kDa), and SLC7A11 (43 kDa) alongside β-actin in ANO6-OE and Vector cells under PBS or PTX challenge, corresponding to Figure 2I. Right panels: Full-length membranes depicting GPX4 (17 kDa), ACSL4 (72 kDa), SLC7A11 (43 kDa), and β-actin across conformational mutant lines (I612A-EV, I612A, D703R-EV, and D703R) under paclitaxel (5 nM PTX) challenge, corresponding to Figure 4F. Black rectangular boxes delineate cropped regions presented in the main figures. Protein molecular weight ladders (in kDa) are indicated alongside each membrane.

**Supplementary Figure S11.**
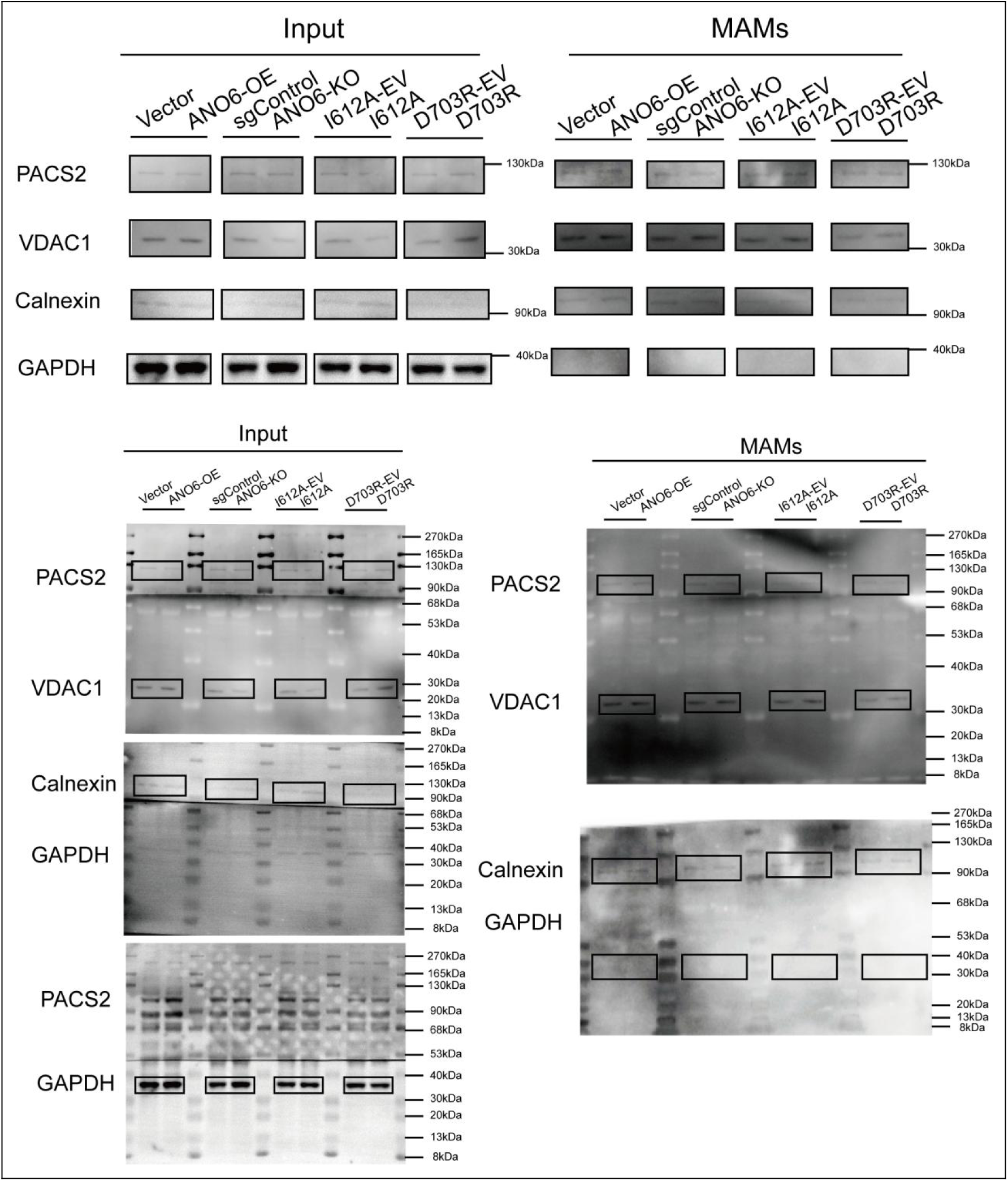
Validation of subcellular fractionation purity for mitochondria-associated endoplasmic reticulum membranes (MAMs). Representative immunoblot analyses and full-length uncropped membrane scans verifying the biochemical purity of isolated mitochondria-associated endoplasmic reticulum membrane (MAMs) fractions compared with whole-cell lysates (Input) across engineered EMT6 cell models (Vector, ANO6-OE, sgControl, ANO6-KO, I612A-EV, I612A, D703R-EV, and D703R) under paclitaxel challenge (5 nM PTX, 24 h) corresponding to Main Figure 5: Top panels: Cropped immunoblot bands displaying compartment-specific marker proteins: PACS2 (canonical MAMs marker, 130 kDa), VDAC1 (outer mitochondrial membrane marker, 30 kDa), Calnexin (endoplasmic reticulum marker, 90 kDa), and GAPDH (cytosolic marker, 40 kDa). The isolated MAMs fractions exhibit marked enrichment of MAMs and organellar contact markers (PACS2, VDAC1, and Calnexin) alongside complete absence of cytosolic contamination (GAPDH), confirming the biochemical purity and structural integrity of the harvested contact microdomains utilized for downstream targeted lipidomics. Bottom panels: Full-length uncropped Western blot membranes corresponding to Input (left column) and isolated MAMs fractions (right column) for each indicated protein target. Black rectangular boxes denote the specific regions cropped for the representative panels displayed above. Molecular weight protein ladders (in kDa) are indicated alongside each membrane.

**Supplementary Figure S12.**
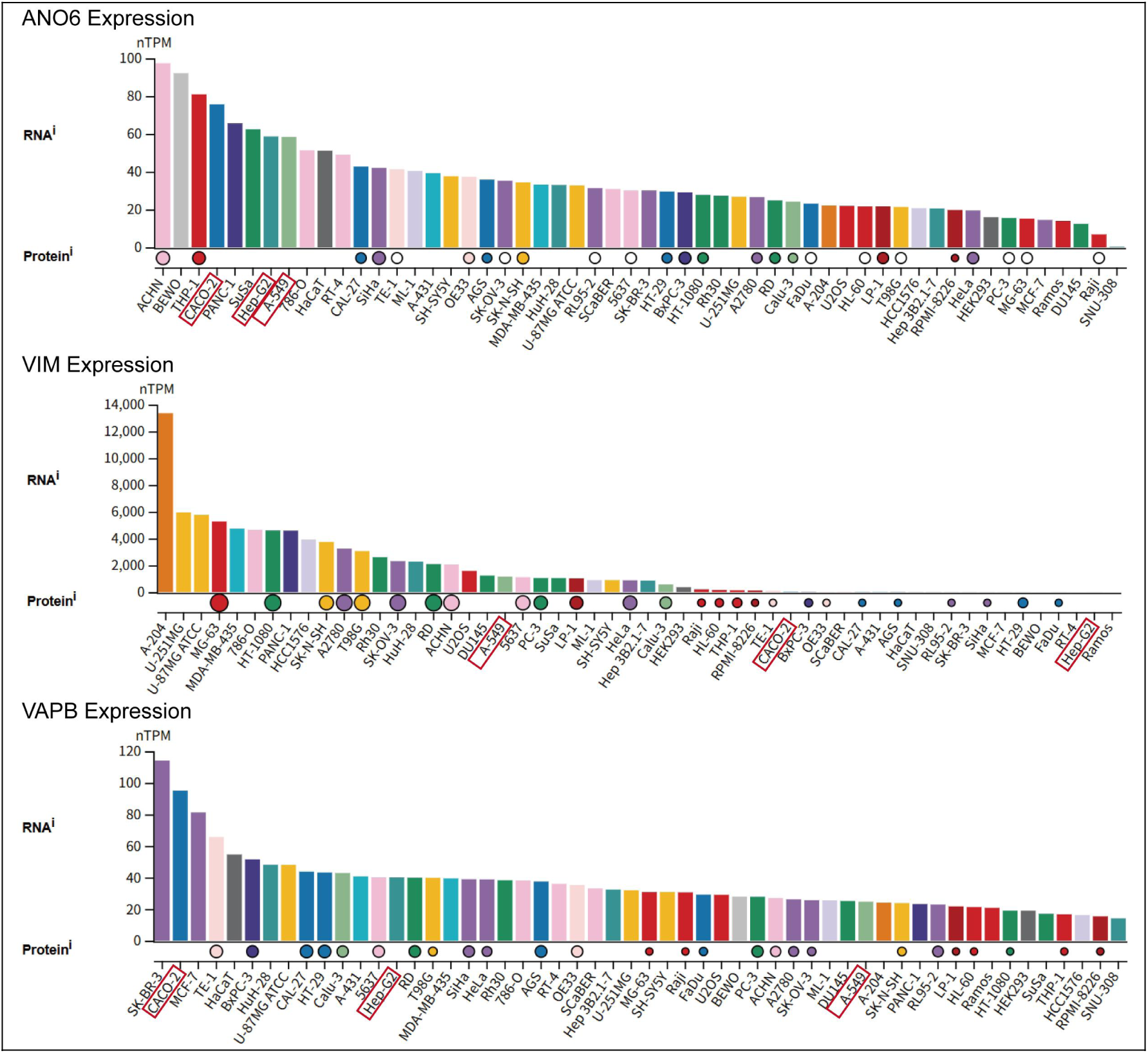
Endogenous expression landscapes of ANO6, VIM, and VAPB across human cancer cell lines in The Human Protein Atlas. Transcript (RNA abundance, normalized transcripts per million, nTPM) and corresponding protein expression profiles of ANO6 (top), VIM (middle), and VAPB (bottom) extracted from The Human Protein Atlas (HPA; https://www.proteinatlas.org) across human cancer cell lines, ranked in descending order of expression. The three cell lines selected for cross-cancer validation in Figure 7G—colorectal carcinoma (CACO-2), non-small cell lung carcinoma (A-549), and hepatocellular carcinoma (Hep-G2)—are highlighted with red bounding boxes along the x-axis, showing robust endogenous ANO6 expression alongside baseline abundance of its tethering partners VIM and VAPB.

---

**Supplementary Table S1.**
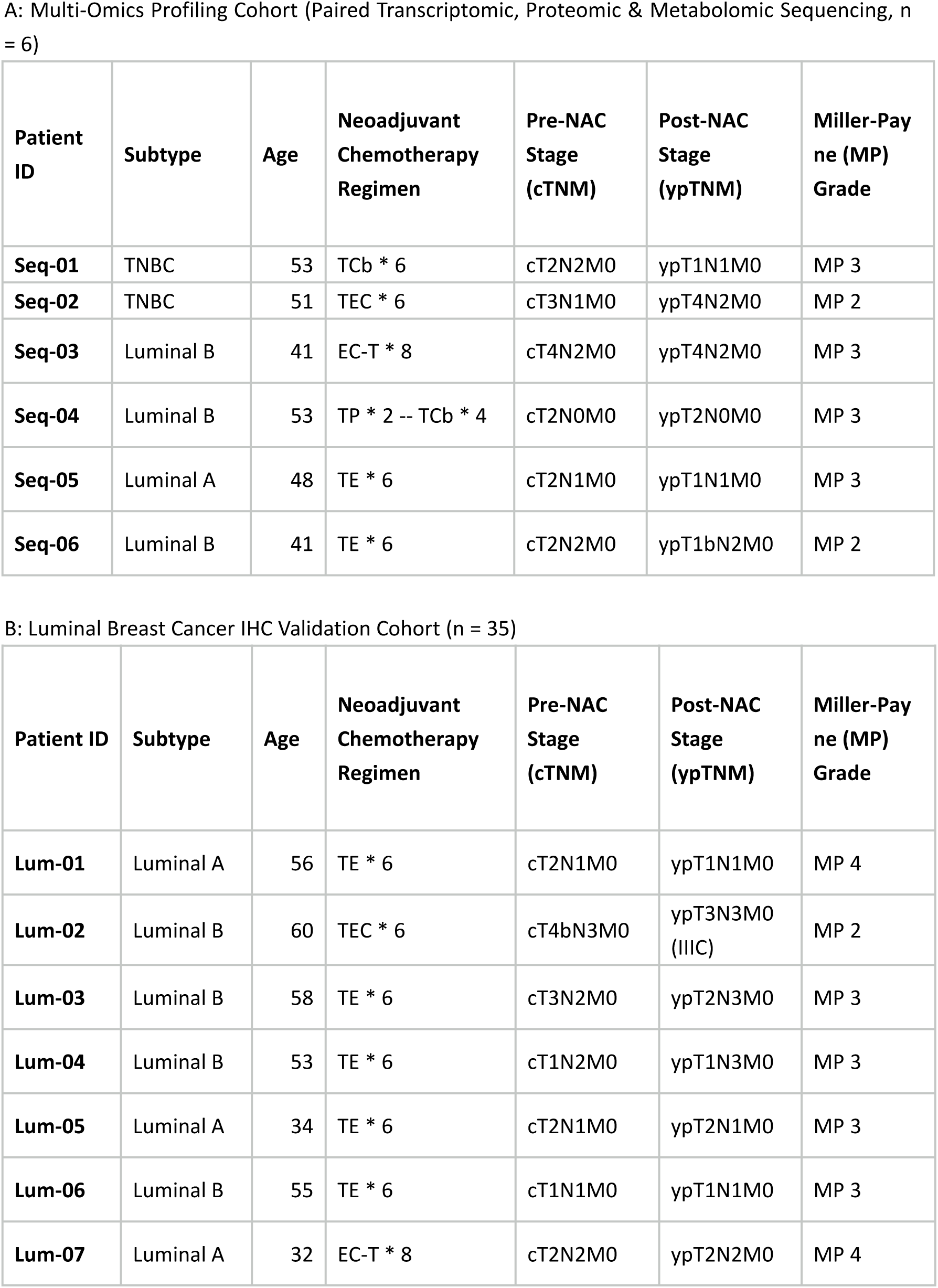
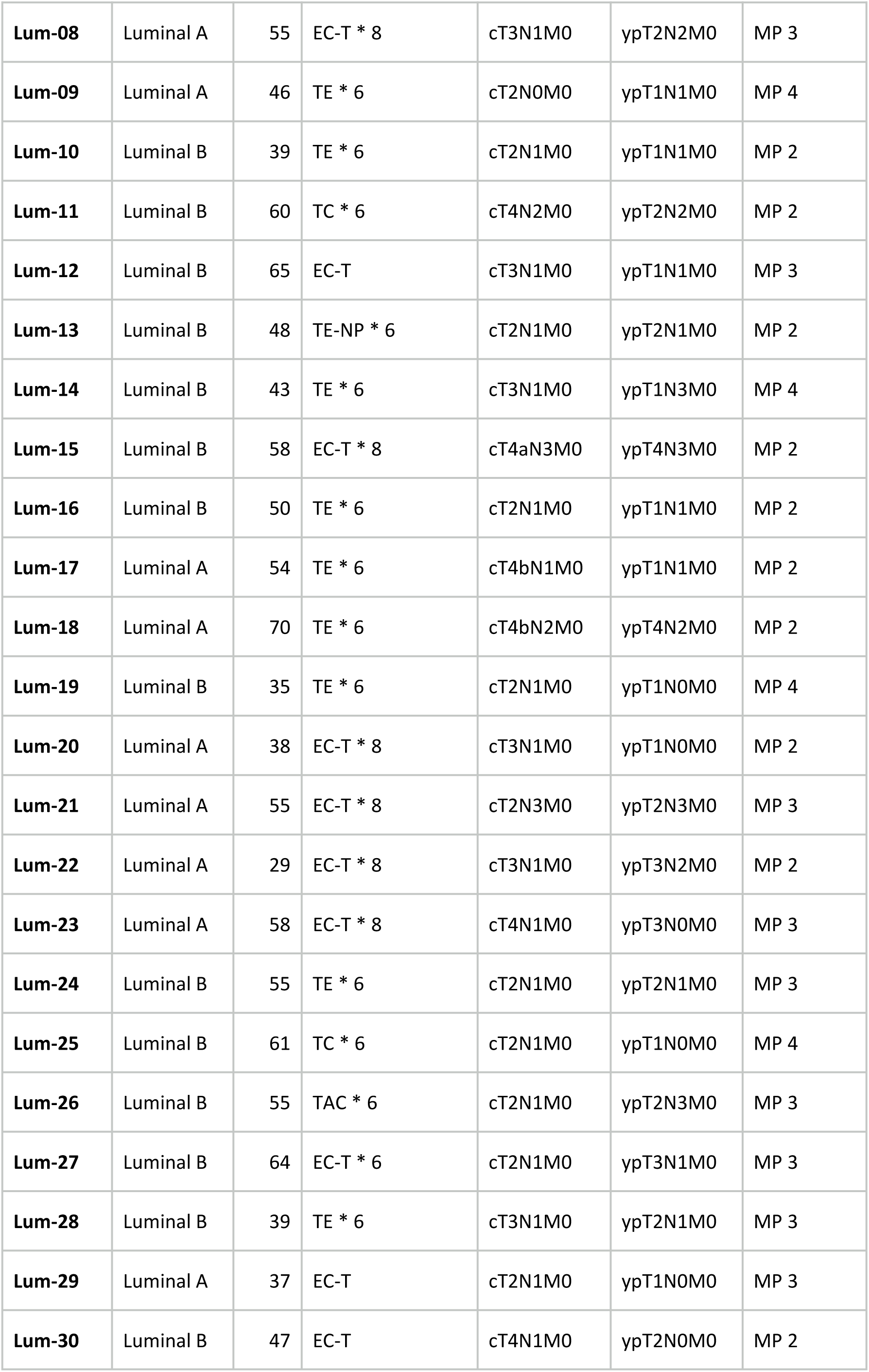
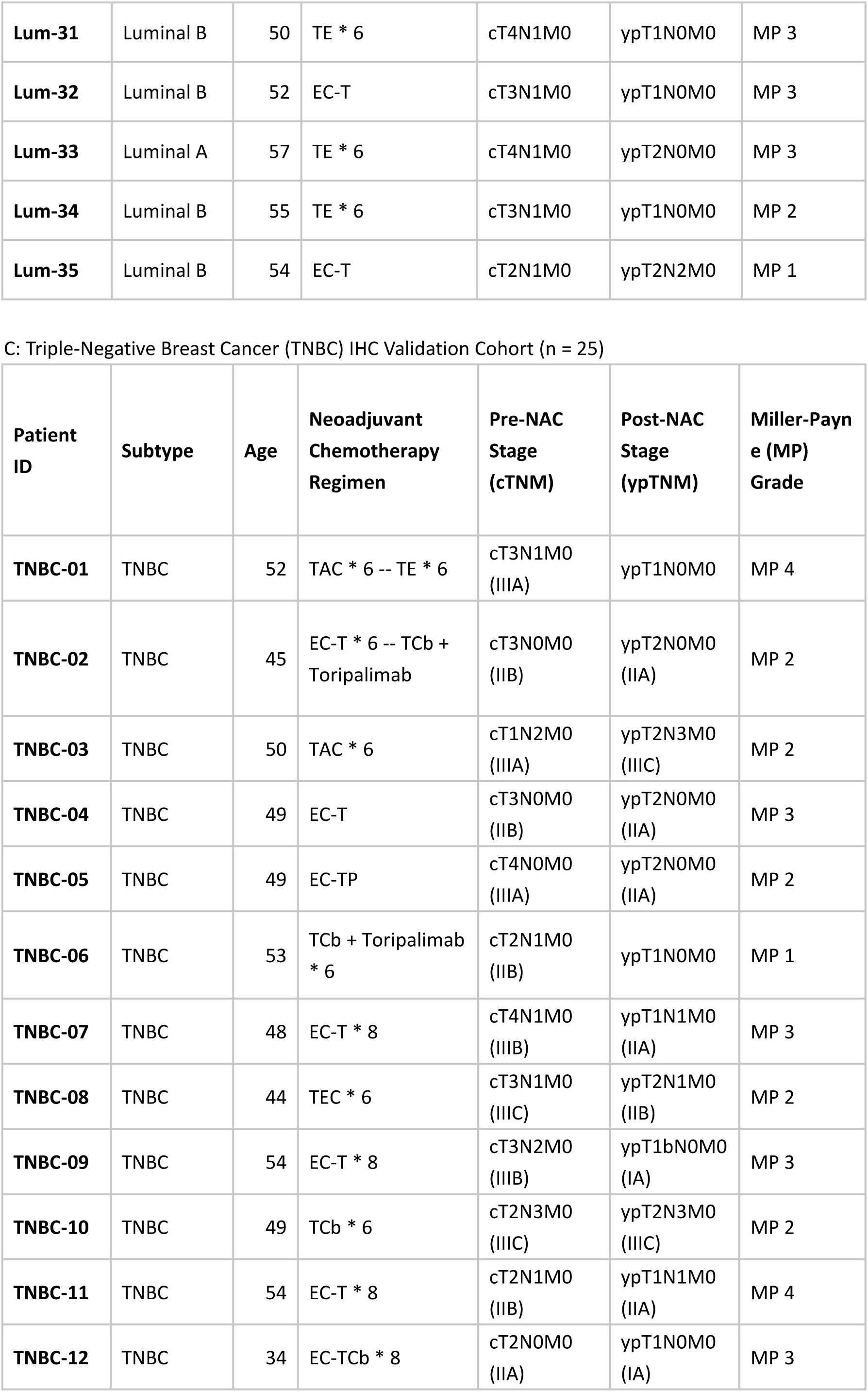
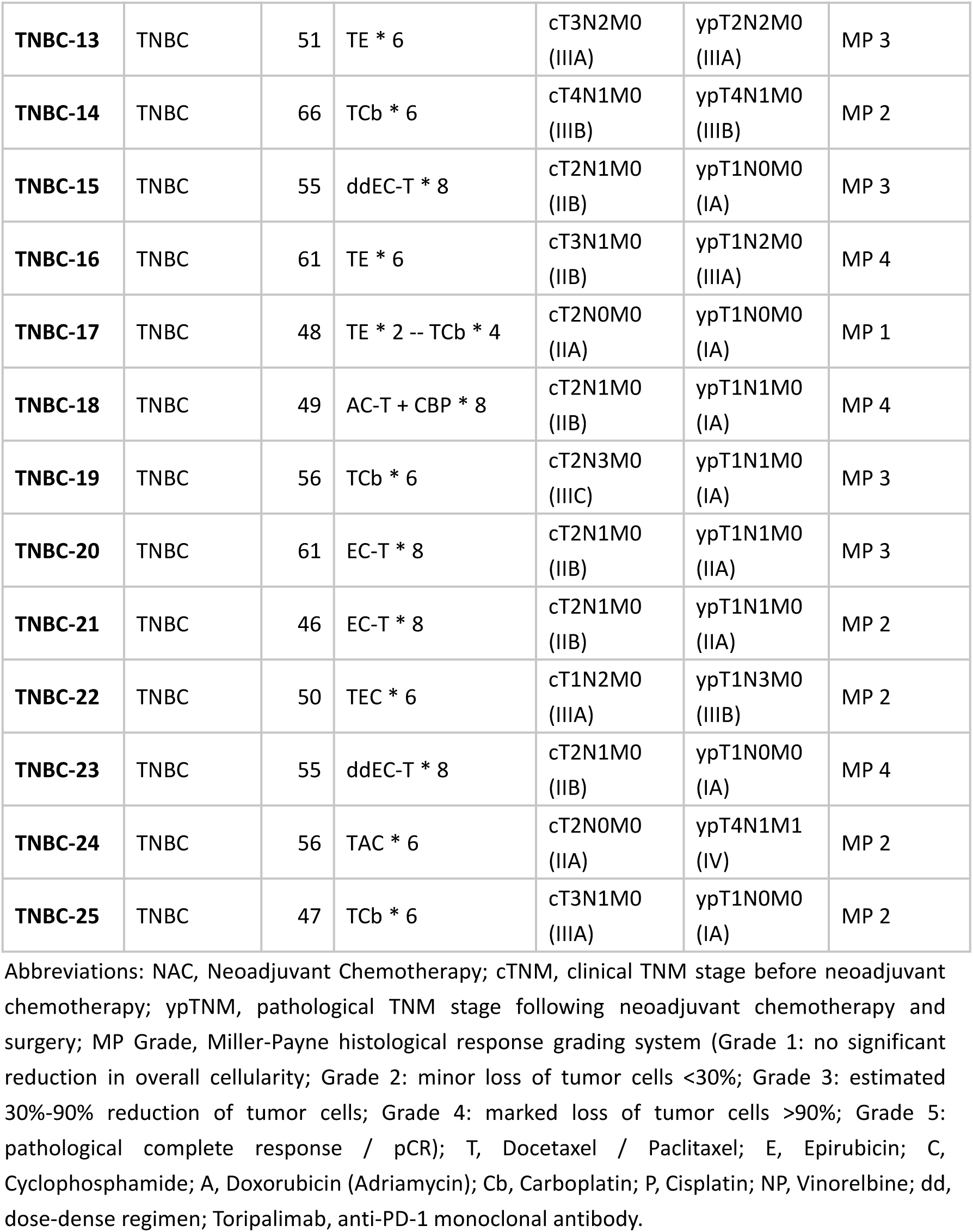
Baseline Clinicopathological Characteristics of Paired Breast Cancer Patient Cohorts (Total n = 66)

**Supplementary Table S2.**
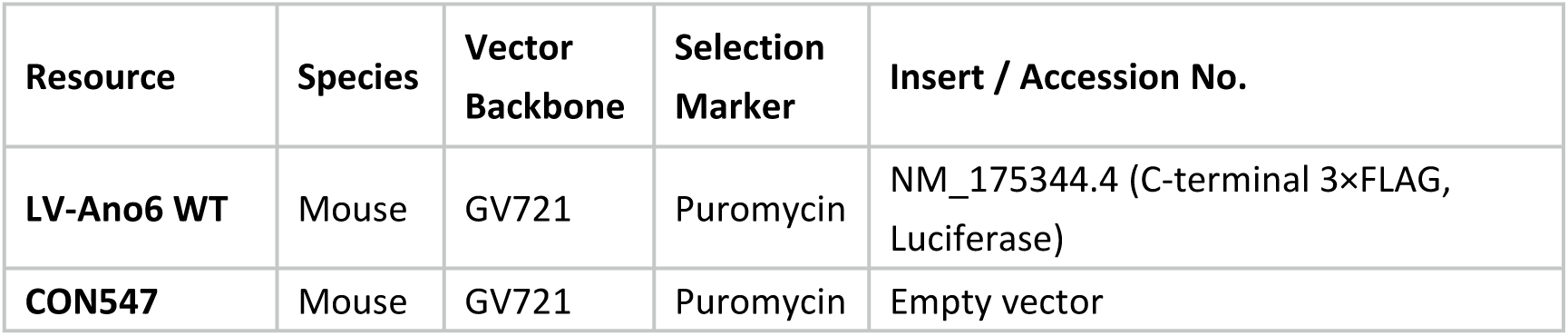
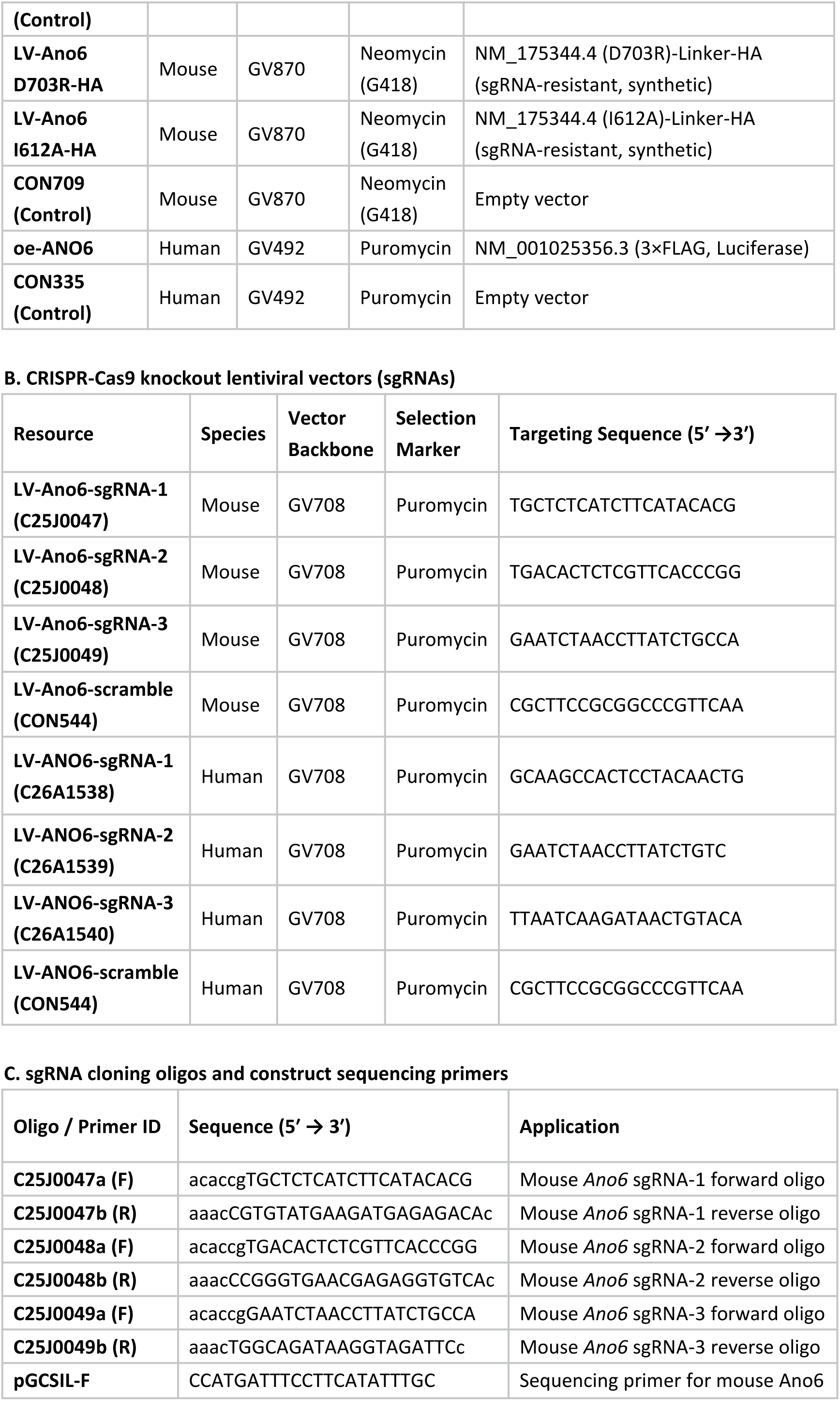
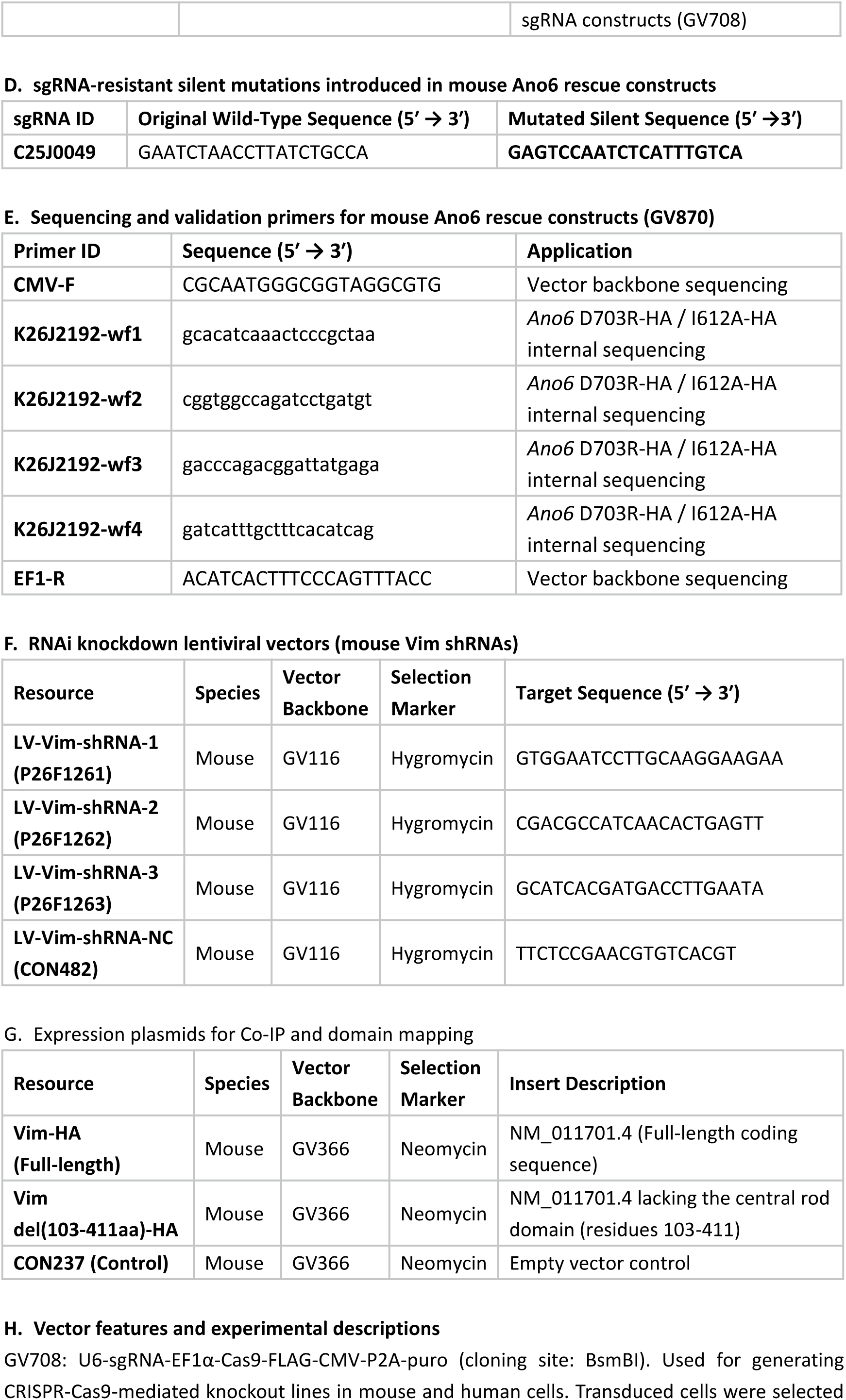
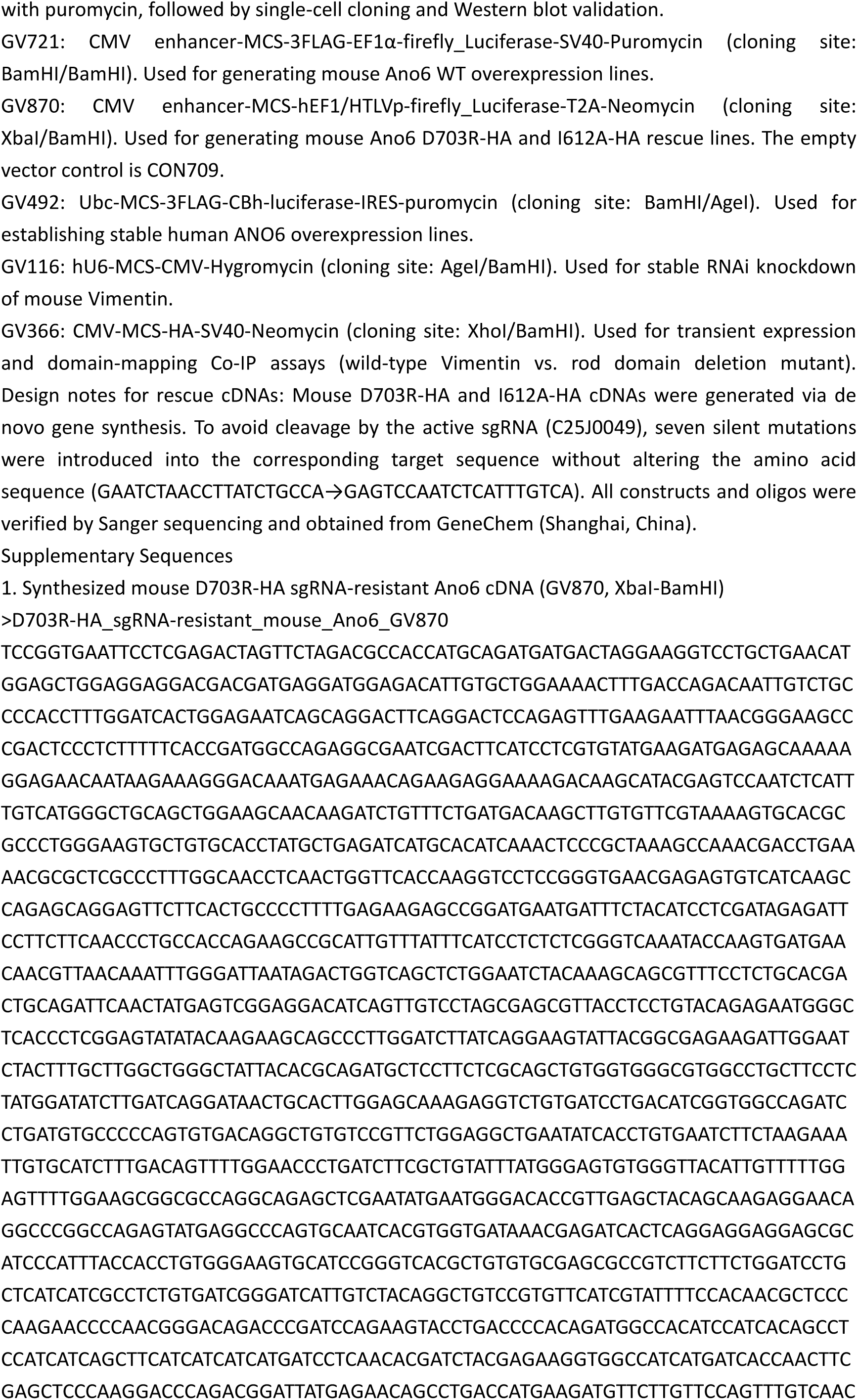
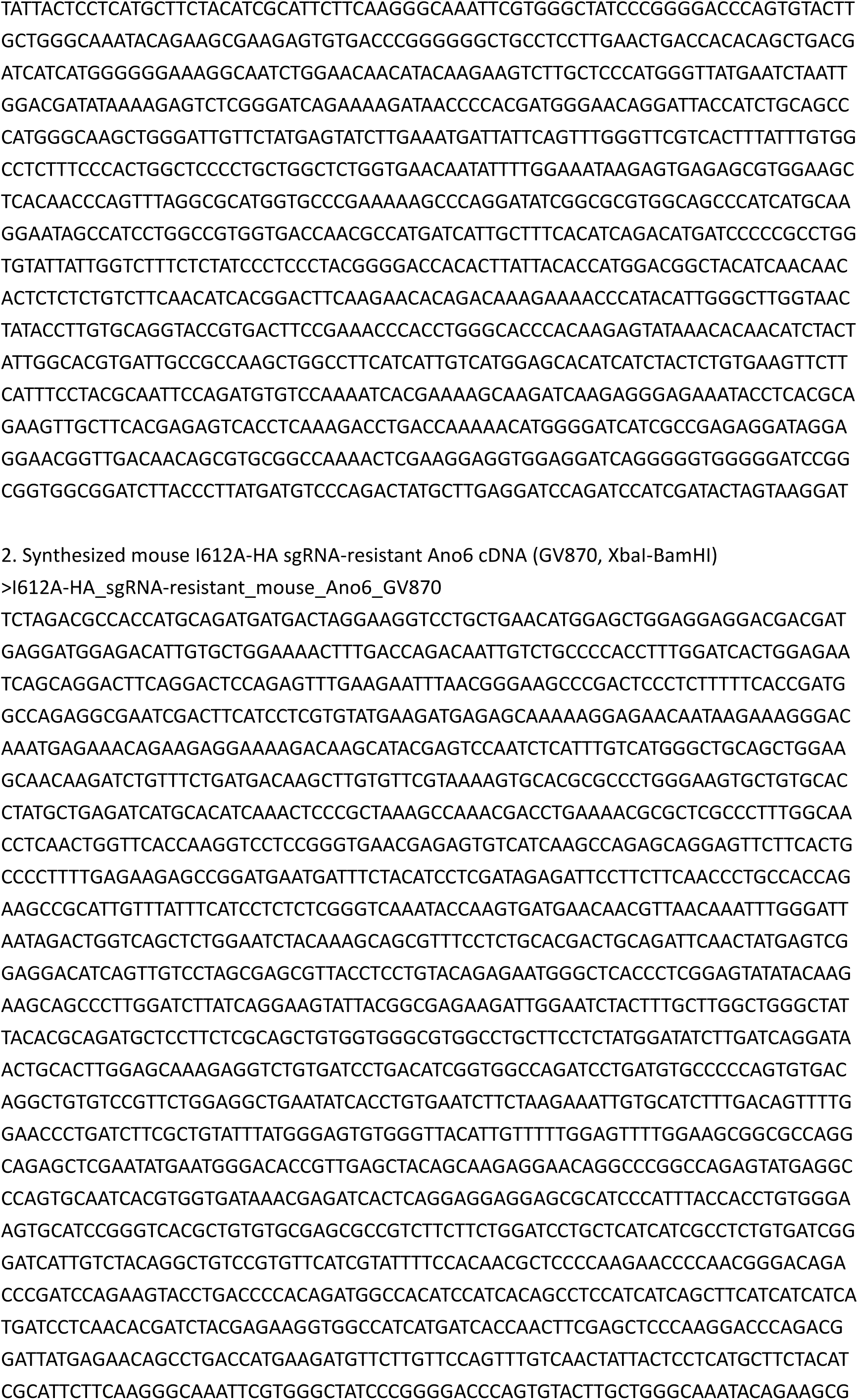
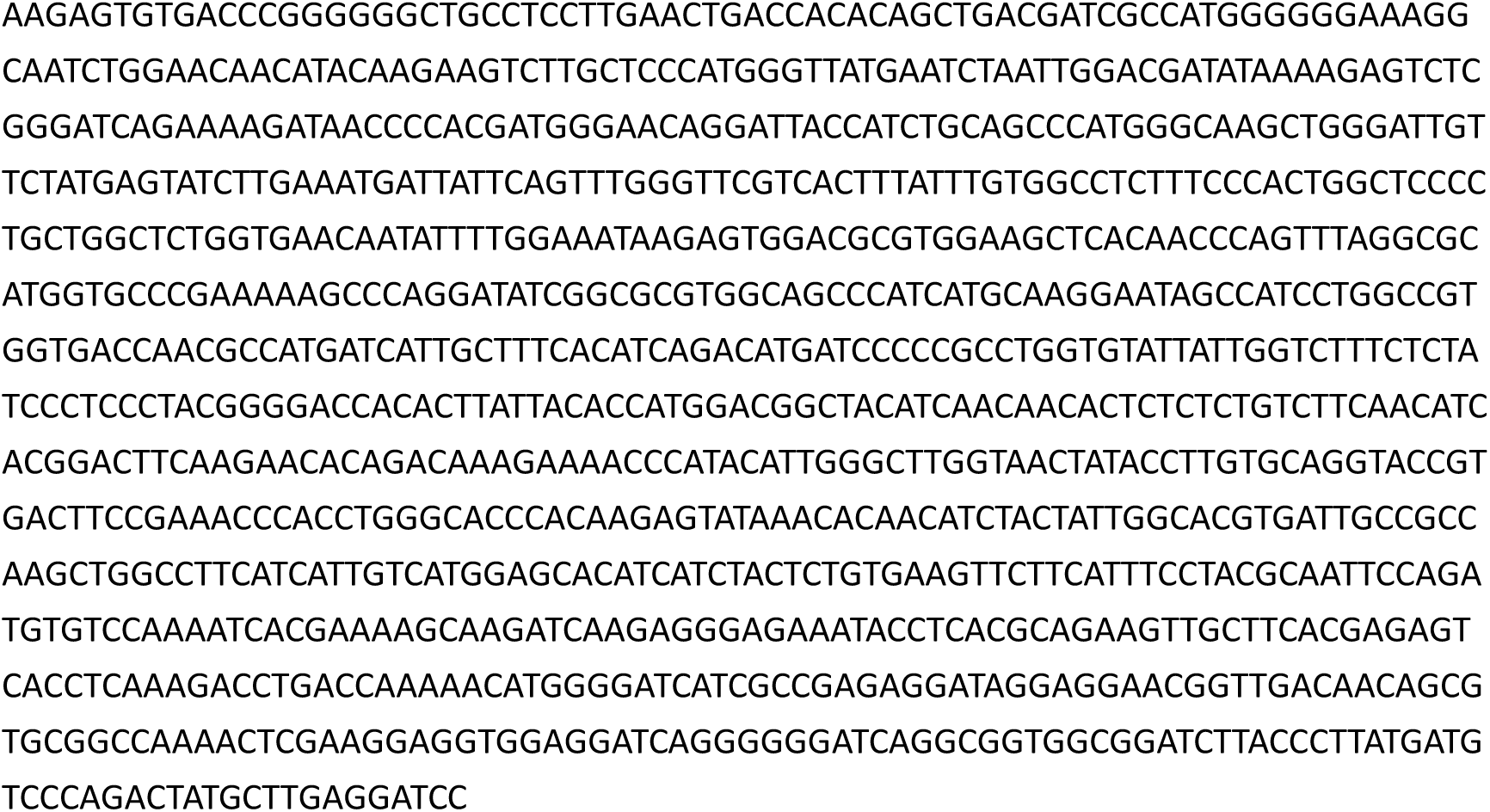
Lentiviruses, plasmids, sgRNAs, and primers for mouse and human Ano6 genetic manipulation and Vim functional studies.

