## Supplementary File for "ANO6 Scaffolds ER-Mitochondria Contacts to Govern Spatial Lipid Compartmentalization and Ferroptosis Evasion in Chemoresistant Breast Cancer"

### Supplementary Figure

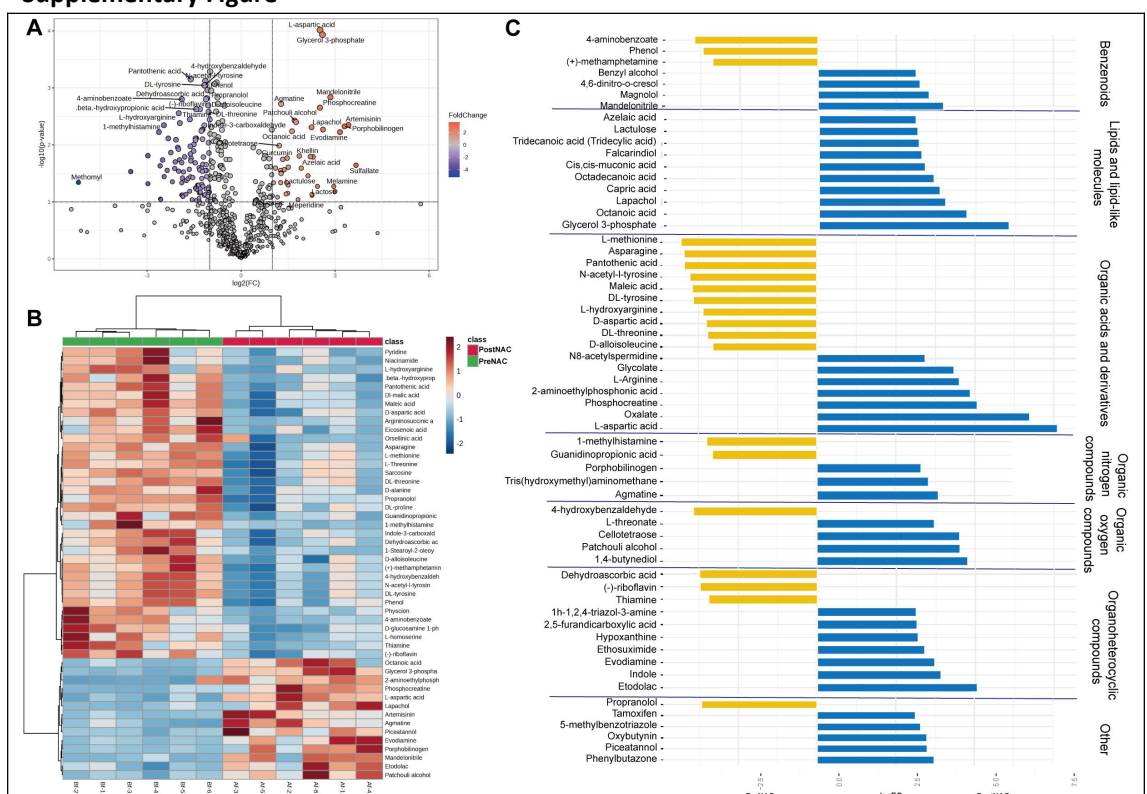

**Supplementary Figure S1. Untargeted metabolomic landscape of paired breast cancer clinical specimens before and after neoadjuvant chemotherapy.**

(A) Volcano plot illustrating differentially abundant metabolites between pre-NAC and post-NAC residual breast cancer lesions (n = 6 paired patient specimens). The horizontal dashed line denotes the statistical significance threshold ($-\log_{10}(P\text{-value}) = 1.3$, corresponding to $P = 0.05$ by two-tailed Student's t-test); vertical dashed lines denote fold-change thresholds ($\log_2FC \geq 0.58$, corresponding to a 1.5-fold difference). Data points are colored according to fold-change gradient (blue: downregulated; red: upregulated in post-NAC residual tissues; gray circles: non-significant). Prominently dysregulated metabolites, including glycerol 3-phosphate, L-aspartic acid, and dehydroascorbic acid, are highlighted. (B) Hierarchical clustering heatmap of significantly altered metabolites across paired patient biopsies. The top column annotations indicate matched clinical cohorts (green: PreNAC; red: PostNAC). Row-normalized relative abundances are expressed as Z-scores, colored along a continuous gradient from blue (low relative abundance) to red (high relative abundance). (C) Class-stratified horizontal bar plot displaying the $\log_2FC$ of representative altered metabolites organized by major biochemical superclasses (Benzenoids, Lipids and lipid-like molecules, Organic acids and derivatives, Organic nitrogen compounds, Organic oxygen compounds, Organoheterocyclic compounds, and Other). Blue bars denote metabolites upregulated in post-chemotherapy residual lesions (PostNAC, $\log_2FC > 0$, highlighted by pronounced accumulation of glycerol 3-phosphate and octanoic acid), and yellow bars denote metabolites enriched in chemotherapy-naïve biopsies (PreNAC, $\log_2FC < 0$, such as 4-aminobenzoate and L-methionine).

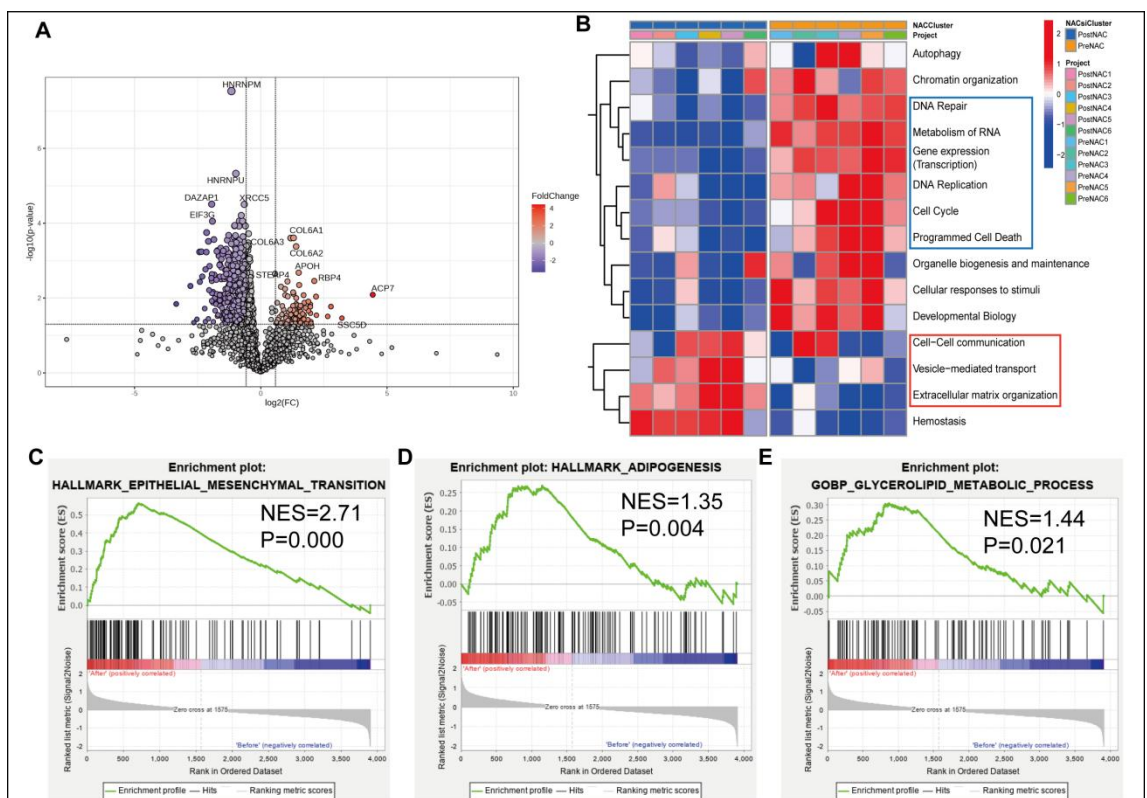

**Supplementary Figure S2. Proteomic landscape and pathway enrichment analysis of paired clinical breast cancer specimens before and after neoadjuvant chemotherapy.**

(A) Volcano plot displaying differentially expressed proteins between pre-NAC baseline biopsies (PreNAC) and post-NAC chemoresistant residual breast cancer tissues (PostNAC) ($n = 6$ paired patient specimens). The horizontal dashed line denotes the statistical significance cutoff ($-\log_{10}(\text{P-value}) = 1.3$, equivalent to $P = 0.05$ by two-tailed Student's t-test); vertical dashed lines represent fold-change thresholds ($\log_2\text{FC} \geq 0.58$, equivalent to a 1.5-fold difference). Data points are colored according to fold-change gradient (blue: downregulated; red: upregulated in PostNAC residual tissues; gray circles: non-significant). Prominently upregulated proteins include ACP7, RBP4, COL6A1, COL6A2, COL6A3, STEAP4, APOH, and SSC5D; prominently downregulated proteins include HNRNPM, HNRNPU, XRCC5, DAZAP1, and EIF3G. (B) Hierarchical clustering heatmap showing the relative activity of core biological process signatures across paired clinical specimens (left columns: PreNAC 1-6; right columns: PostNAC 1-6, annotated by top color bars). Color scale reflects row-normalized Z-scores from low (blue) to high (red) pathway enrichment. The blue bounding box highlights biological processes prominently elevated in post-chemotherapy residual lesions (including DNA repair, metabolism of RNA, gene expression/transcription, DNA replication, cell cycle, and programmed cell death); the red bounding box marks stromal and communication signatures exhibiting higher relative enrichment in treatment-naïve baseline biopsies (cell-cell communication, vesicle-mediated transport, and extracellular matrix organization). (C-E) Gene Set Enrichment Analysis (GSEA) plots illustrating key oncogenic and metabolic pathways positively enriched in post-NAC chemoresistant residual tissues (PostNAC): (C) Epithelial-to-mesenchymal transition hallmark signature (HALLMARK\_EPITHELIAL\_MESENCHYMAL\_TRANSITION; normalized enrichment score NES = 2.71, $P < 0.001$); (D) Adipogenesis hallmark signature (HALLMARK\_ADIPOGENESIS; NES = 1.35, $P = 0.004$); (E) Glycerolipid metabolic process Gene Ontology pathway

(GOBP\_GLYCEROLIPID\_METABOLIC\_PROCESS; NES = 1.44, P = 0.021).

### EMT-6

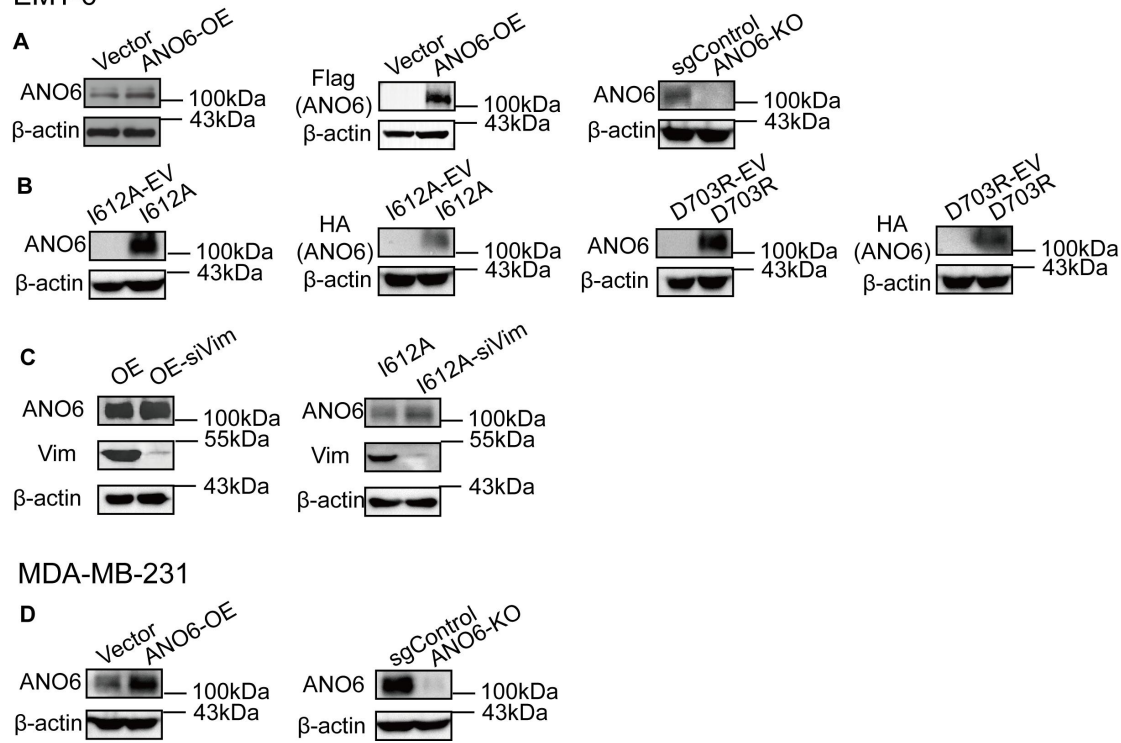

### Supplementary Figure S3. Western blot validation of engineered breast cancer cell models.

Representative immunoblot analyses confirming the establishment, genetic manipulation, and reconstitution of experimental cell models in mouse EMT-6 and human MDA-MB-231 breast cancer backgrounds: (A-C) EMT-6 cell models: (A) ANO6 overexpression and knockout: Verification of ectopic ANO6 expression in Vector versus ANO6-OE cells probed with anti-ANO6 and anti-Flag tag antibodies (left and middle), alongside confirmation of CRISPR-Cas9-mediated gene disruption displaying complete loss of endogenous ANO6 protein in ANO6-KO cells relative to non-targeting sgControl cells (right). (B) Conformational mutant reconstitution: Immunoblot validation of stable reconstitution in engineered EMT-6 lines transduced with matched empty vectors or constructs encoding the active scaffolding mutant (I612A-EV vs. I612A) or the scaffolding-compromised mutant (D703R-EV vs. D703R), detected using anti-ANO6 and anti-HA tag antibodies. (C) Vimentin knockdown: Evaluation of siRNA-mediated silencing of Vimentin (siVim) in ANO6-OE (OE vs. OE-siVim) and I612A (I612A vs. I612A-siVim) backgrounds, confirming robust depletion of Vim protein (55 kDa) without altering ANO6 abundance (100 kDa). (D) MDA-MB-231 cell models: Immunoblot validation of ANO6 overexpression (Vector vs. ANO6-OE, left) and CRISPR-Cas9-mediated knockout (sgControl vs. ANO6-KO, right) in human MDA-MB-231 triple-negative breast cancer cells probed with an anti-ANO6 antibody. In all panels, β-actin (43 kDa) served as the internal loading control. Molecular weights are indicated in kilodaltons (kDa) on the right of each blot.

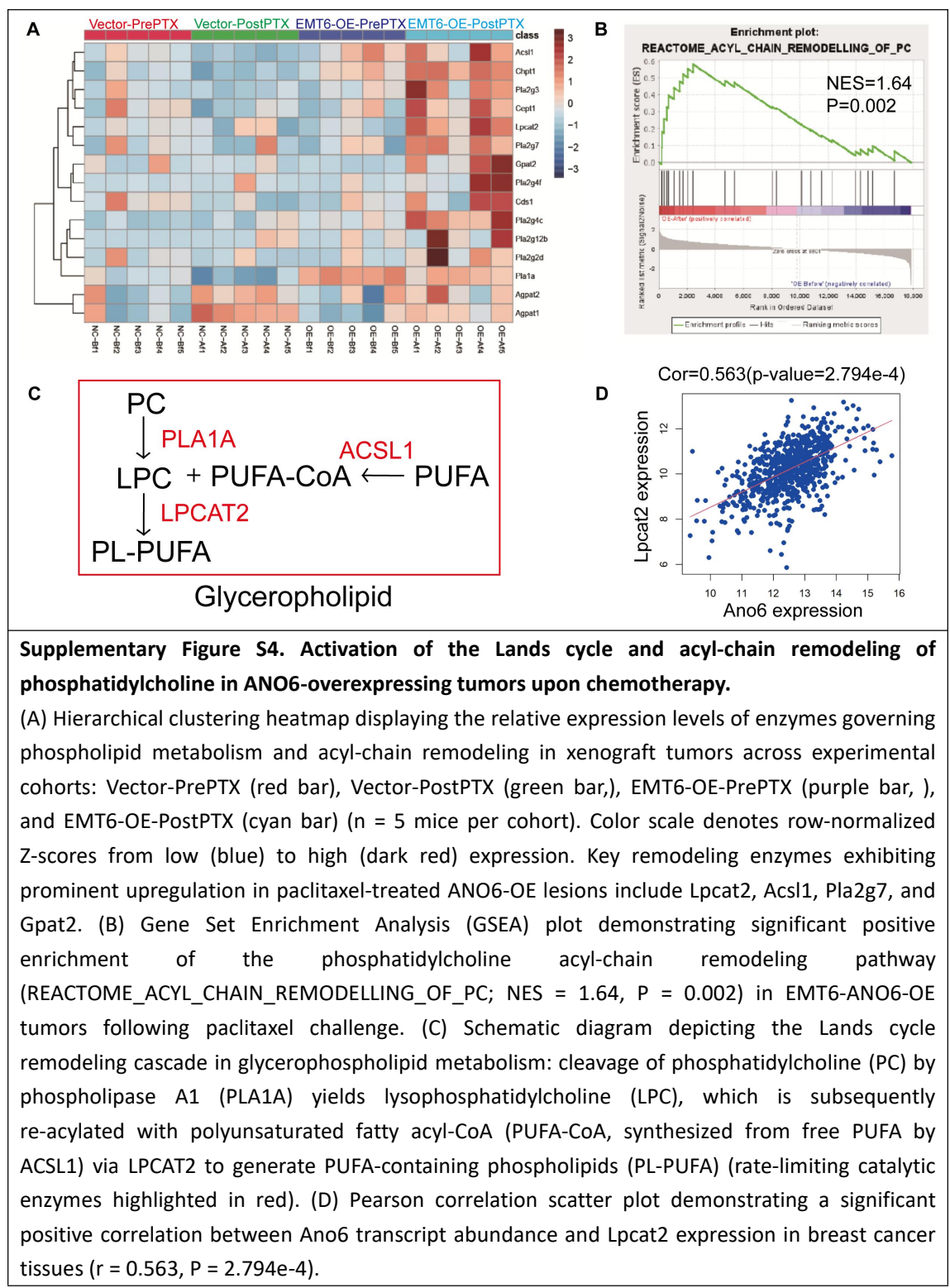

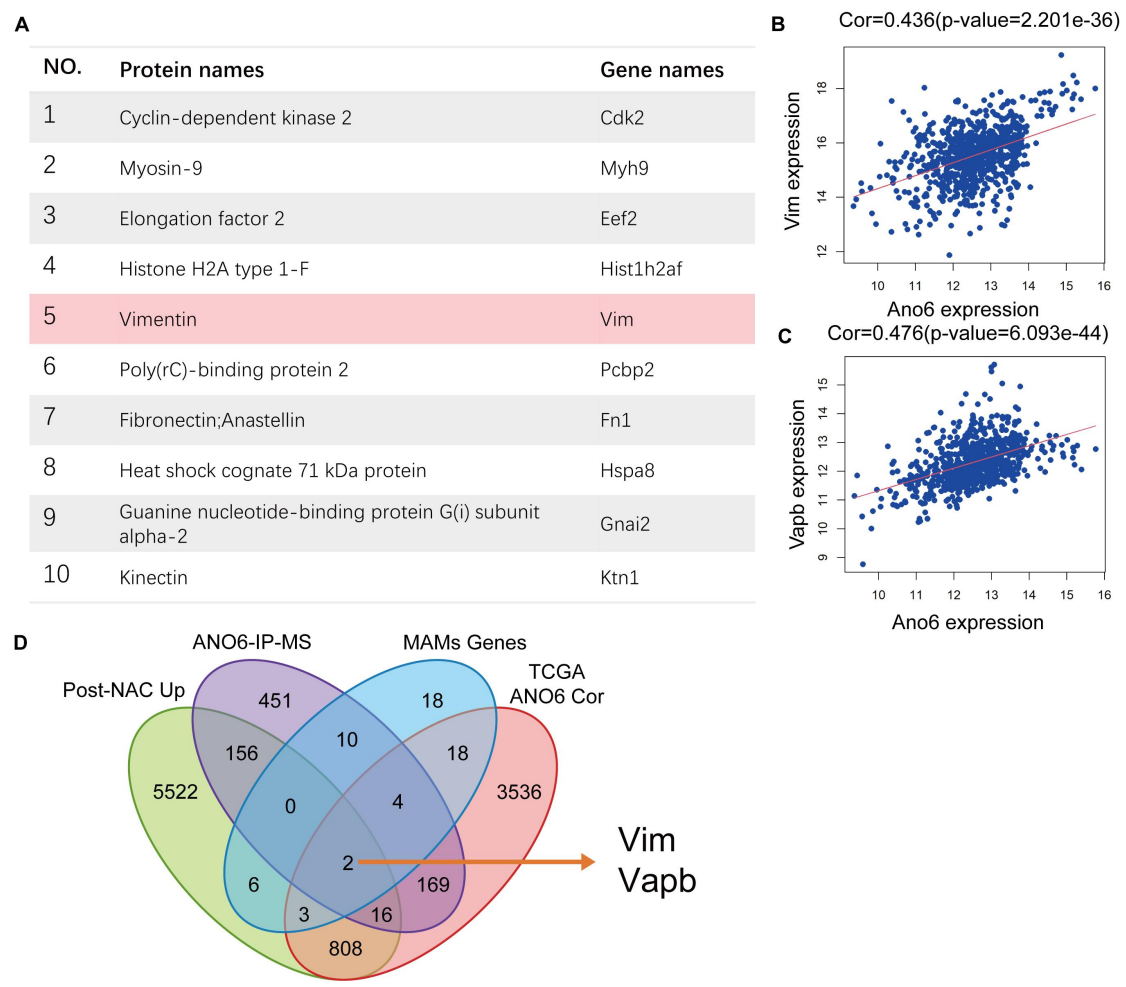

**Supplementary Figure S5. Integrated mass spectrometry and multi-cohort intersection identify Vimentin and VAPB as core ANO6-associated MAMs scaffolding partners.**

(A) Top 10 candidate interacting proteins identified by co-immunoprecipitation followed by liquid chromatography-tandem mass spectrometry (ANO6-IP-MS) in breast cancer cells. Vimentin (VIM/Vim) is highlighted with light pink shading in row 5. (B, C) Pearson correlation scatter plots demonstrating significant positive associations between Ano6 mRNA expression and Vim ($r = 0.436$, $P = 2.201 \times 10^{-36}$; B) as well as Vapb ($r = 0.476$, $P = 6.093 \times 10^{-44}$; C) across breast cancer patient cohorts. Red lines represent linear regression fits. (D) Four-way Venn diagram illustrating the multi-omic filtering strategy intersecting: (1) genes significantly upregulated in post-chemotherapy residual clinical specimens (Post-NAC Up), (2) physical binding candidates identified by ANO6-IP-MS, (3) curated mitochondria-associated ER membrane components (MAMs Genes), and (4) genes exhibiting positive expression correlation with ANO6 in the TCGA breast cancer cohort (TCGA ANO6 Cor). The intersection uniquely converged on two core scaffolding candidates (indicated by arrow): Vim and Vapb.

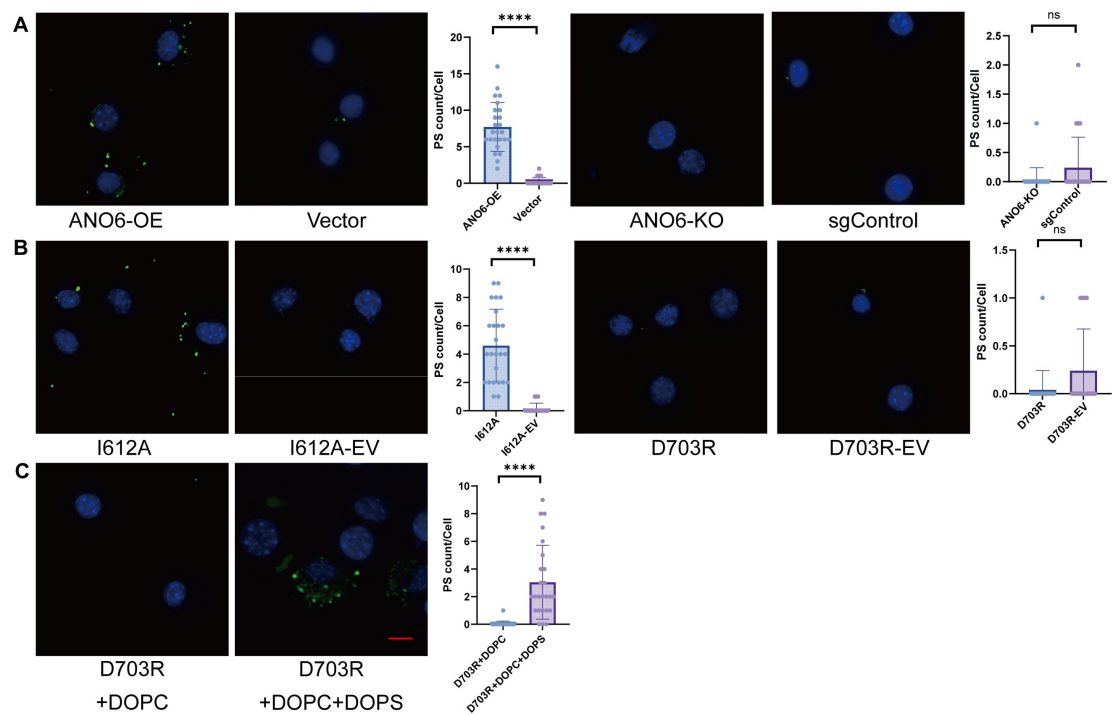

**Supplementary Figure S6. In situ fluorescent imaging of cell-surface phosphatidylserine exposure across engineered ANO6 cell models.**

exposure across engineered ANO6 cell models. Representative high-resolution live-cell confocal micrographs visualizing cell-surface phosphatidylserine (PS) externalization (green puncta; stained with lactadherin-FITC; scale bar = 25 μm) counterstained with Hoechst 33342 (nuclei, blue), alongside corresponding single-cell dot-plot quantifications of surface PS puncta (PS count/cell): (A) Assessment of surface PS puncta in wild-type ANO6 models, comparing ANO6-overexpressing cells with Vector controls (left; \*\*\*\*$P < 0.0001$ by two-tailed unpaired Student's t-test), and ANO6-knockout (ANO6-KO) cells with sgControl cells (right; ns, not significant). (B) Evaluation of conformational mutants showing robust surface PS puncta accumulation in the active scaffolding mutant (I612A) relative to its empty vector control (I612A-EV; $P < 0.0001$), whereas the scaffolding-compromised mutant/catalytically dead mutant (D703R) displayed baseline-level signals indistinguishable from D703R-EV (right; ns). (C) Exogenous replenishment of serine-containing liposomes (DOPC + DOPS) on D703R cells selectively restores cell-surface PS puncta decoration relative to neutral liposomes (DOPC alone) ($P < 0.0001$). Data are presented as mean ± SD from three independent biological replicates. \*\*\*\*$P < 0.0001$; ns, not significant.

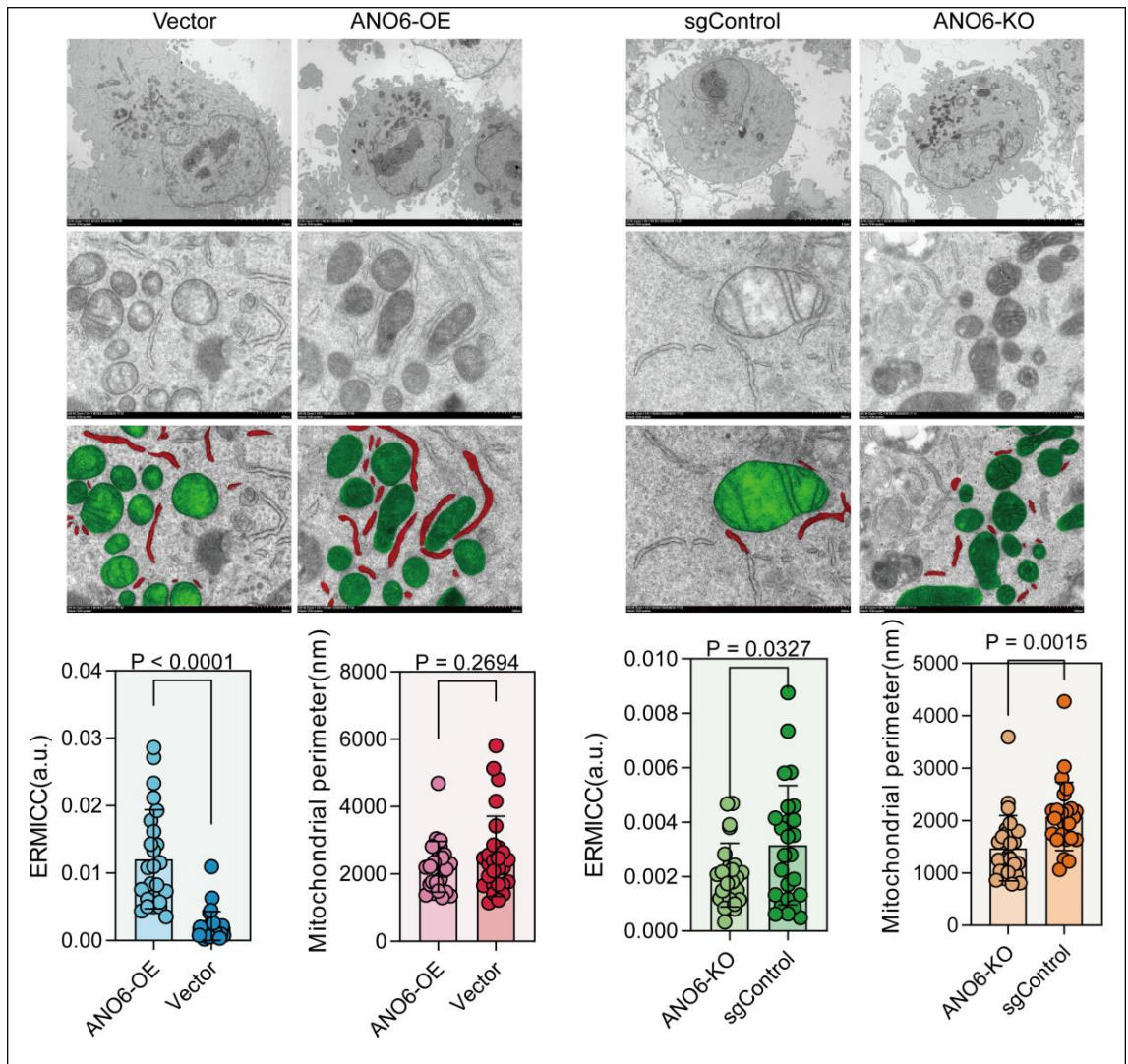

**Supplementary Figure S7. Transmission electron microscopy evaluates the modulation of ER-mitochondria contact ultrastructure by ANO6 abundance in MDA-MB-231.**

Representative transmission electron microscopy (TEM) micrographs and morphometric quantifications assessing mitochondria-associated endoplasmic reticulum membrane (MAMs) architecture in human MDA-MB-231 triple-negative breast cancer cells across comparative genetic cohorts: control (Vector) versus ANO6-overexpressing (ANO6-OE) cells, and non-targeting control (sgControl) versus ANO6-knockout (ANO6-KO) cells. Top and middle panels: Low-magnification whole-cell overviews (top row) and high-magnification micrographs capturing inter-organellar contact interfaces (middle row) (scale bars indicated in micrographs). In the third row, mitochondria are pseudocolored in green, and endoplasmic reticulum (ER) cisternae closely apposed to the outer mitochondrial membrane ($\leq 60$ nm cleft distance) are highlighted in red. Bottom plots (left): Scatter dot-plot quantifications of the ER-mitochondria contact coefficient (ERMICC in arbitrary units [a.u.], left plot; $P < 0.0001$) and mitochondrial perimeter (nm, right plot; $P = 0.2694$; ns, not significant) comparing ANO6-OE with Vector cells evaluated by two-tailed unpaired Student's t-test. Bottom plots (right): Quantitative morphometry of ERMICC (left plot; $P = 0.0327$) and mitochondrial perimeter (right plot; $P = 0.0015$) comparing ANO6-KO with sgControl cells. Data are presented as mean ± SD from independent biological replicates. \*$P < 0.05$, \*\*$P < 0.01$, \*\*\*\*$P < 0.0001$; ns, not significant.

### EMT-6

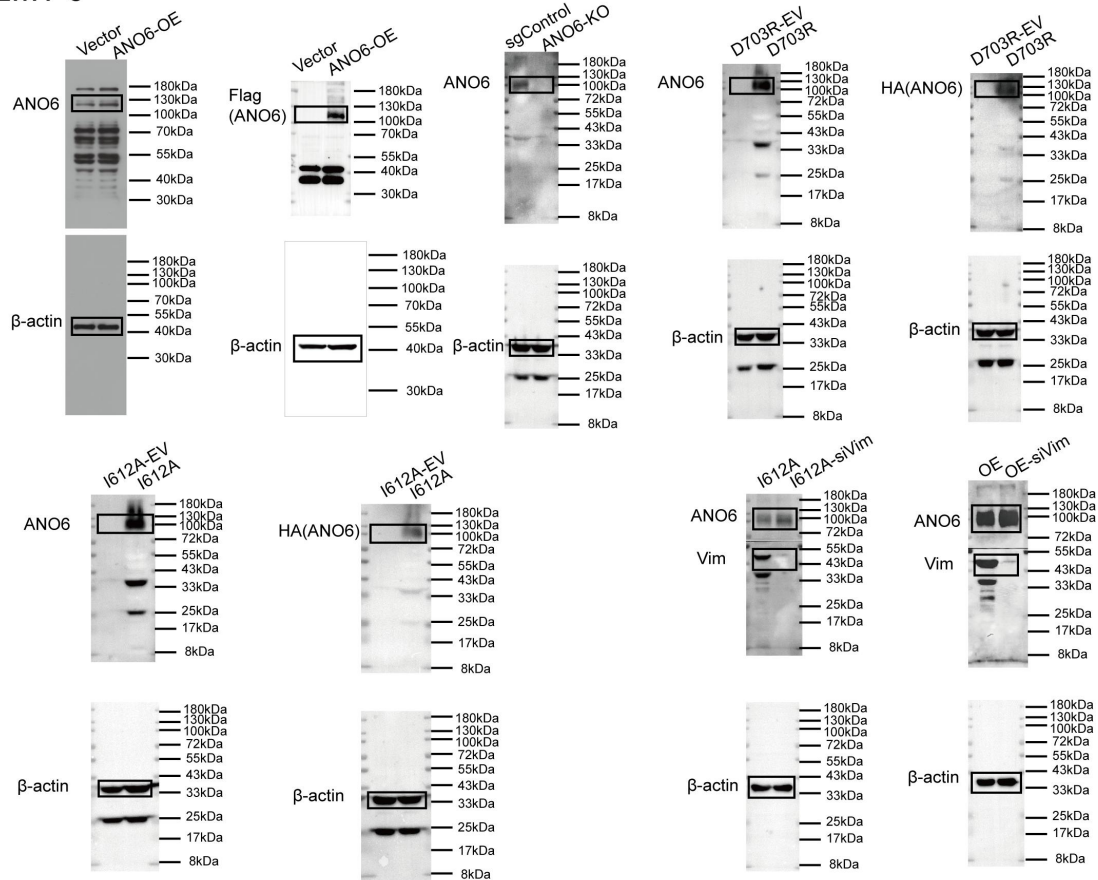

### MDA-MB-231

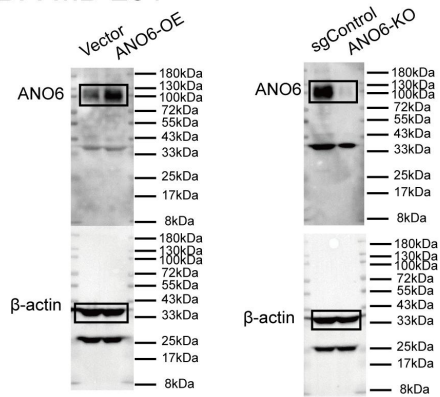

**Supplementary Figure S8. Full-length uncropped immunoblot images corresponding to Supplementary Figure S3.**

Uncropped Western blot membranes displaying target proteins and their corresponding loading controls (β-actin) used for the genetic validation of engineered mouse EMT-6 and human MDA-MB-231 breast cancer cell models. Black rectangular boxes denote the cropped regions presented in Supplementary Figure S3. Protein molecular weight ladders (in kDa) are indicated adjacent to each membrane.

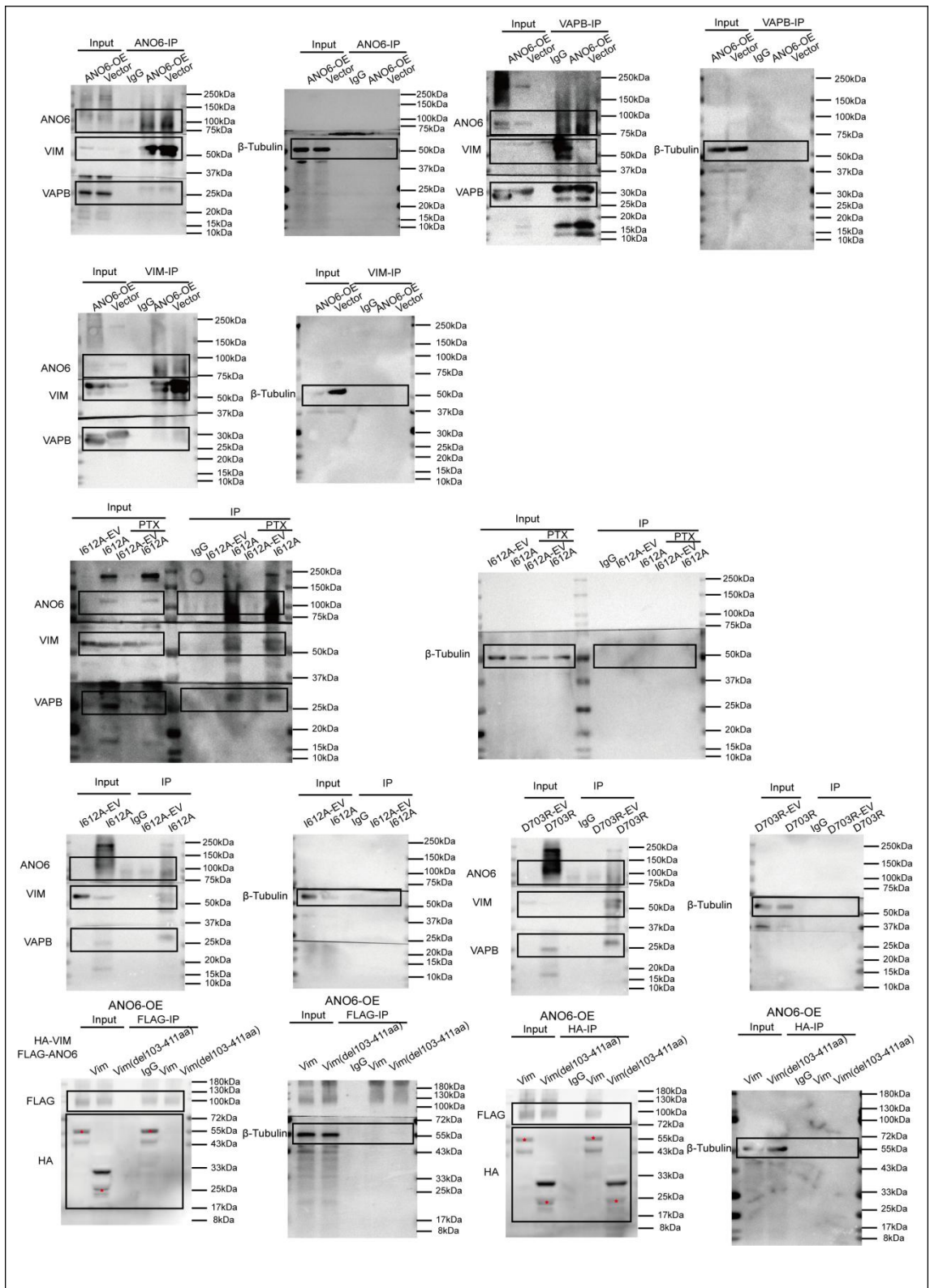

**Supplementary Figure S9. Full-length uncropped immunoblot images corresponding to Co-IP assays in Figure 3 and Figure 6.**

Uncropped Western blot membranes displaying target proteins and corresponding input loading controls (β-tubulin, 55 kDa) for all co-immunoprecipitation (Co-IP) experiments presented in the manuscript: Rows 1 and 2 (top): Uncropped membranes for reciprocal endogenous Co-IP assays evaluating mutual physical interactions among ANO6, Vimentin (VIM), and VAPB in Vector and

ANO6-OE cells following anti-ANO6, anti-VAPB, or anti-VIM pull-down, corresponding to Figure 3C. Row 3 (middle): Uncropped membranes assessing physical interactions of the active scaffolding mutant I612A with VIM and VAPB in the presence or absence of paclitaxel (PTX) challenge, corresponding to Figure 3D. Row 4: Uncropped membranes evaluating physical binding capacities toward VIM and VAPB between the active scaffolding mutant I612A and the scaffold-compromised mutant D703R relative to their matched empty vector controls (I612A-EV and D703R-EV), corresponding to \*\*Figure 3E. Row 5 (bottom): Uncropped membranes for reciprocal Co-IP assays mapping the interaction interface between FLAG-ANO6 and HA-tagged full-length VIM or the rod-domain deletion mutant HA-Vim(Δ103-411aa) following anti-FLAG or anti-HA pull-down, corresponding to Figure 6G. Black rectangular boxes delineate cropped blot areas displayed in main figures. Red asterisks mark target recombinant truncation bands. Protein molecular weight ladders (in kDa) are indicated adjacent to each membrane.

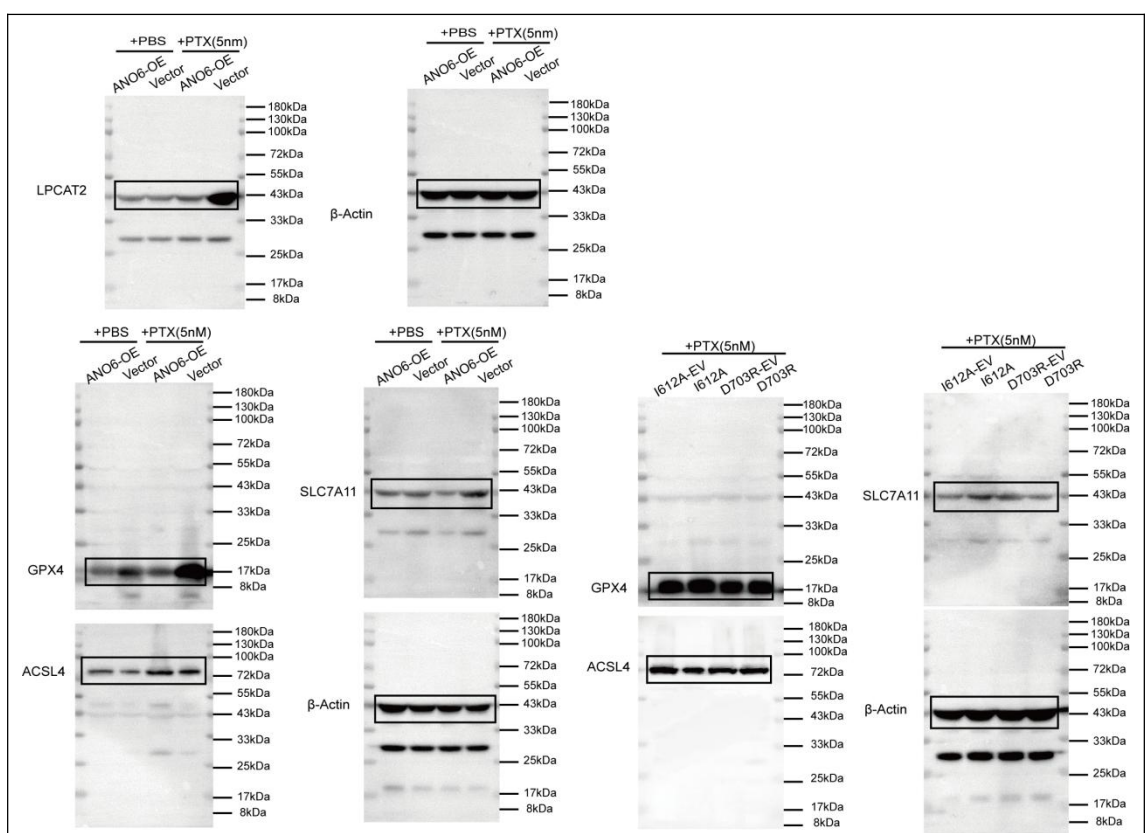

**Supplementary Figure S10. Full-length uncropped immunoblot images corresponding to Main Figure 2 and Figure 4.**

Uncropped Western blot membranes displaying target proteins and corresponding loading controls (β-actin, 43 kDa) for lipid-remodeling enzymes and canonical ferroptosis surveillance machinery: Top panels: Full-length membranes for LPCAT2 (43 kDa) and matched β-actin in ANO6-OE and Vector cells treated with vehicle (PBS) or paclitaxel (5 nM PTX), corresponding to Figure 2H. Bottom-left panels: Full-length membranes evaluating canonical ferroptosis surveillance proteins GPX4 (17 kDa), ACSL4 (72 kDa), and SLC7A11 (43 kDa) alongside β-actin in ANO6-OE and Vector cells under PBS or PTX challenge, corresponding to Figure 2I. Right panels: Full-length membranes depicting GPX4 (17 kDa), ACSL4 (72 kDa), SLC7A11 (43 kDa), and β-actin across conformational mutant lines (I612A-EV, I612A, D703R-EV, and D703R) under paclitaxel (5

nM PTX) challenge, corresponding to Figure 4F. Black rectangular boxes delineate cropped regions presented in the main figures. Protein molecular weight ladders (in kDa) are indicated alongside each membrane.

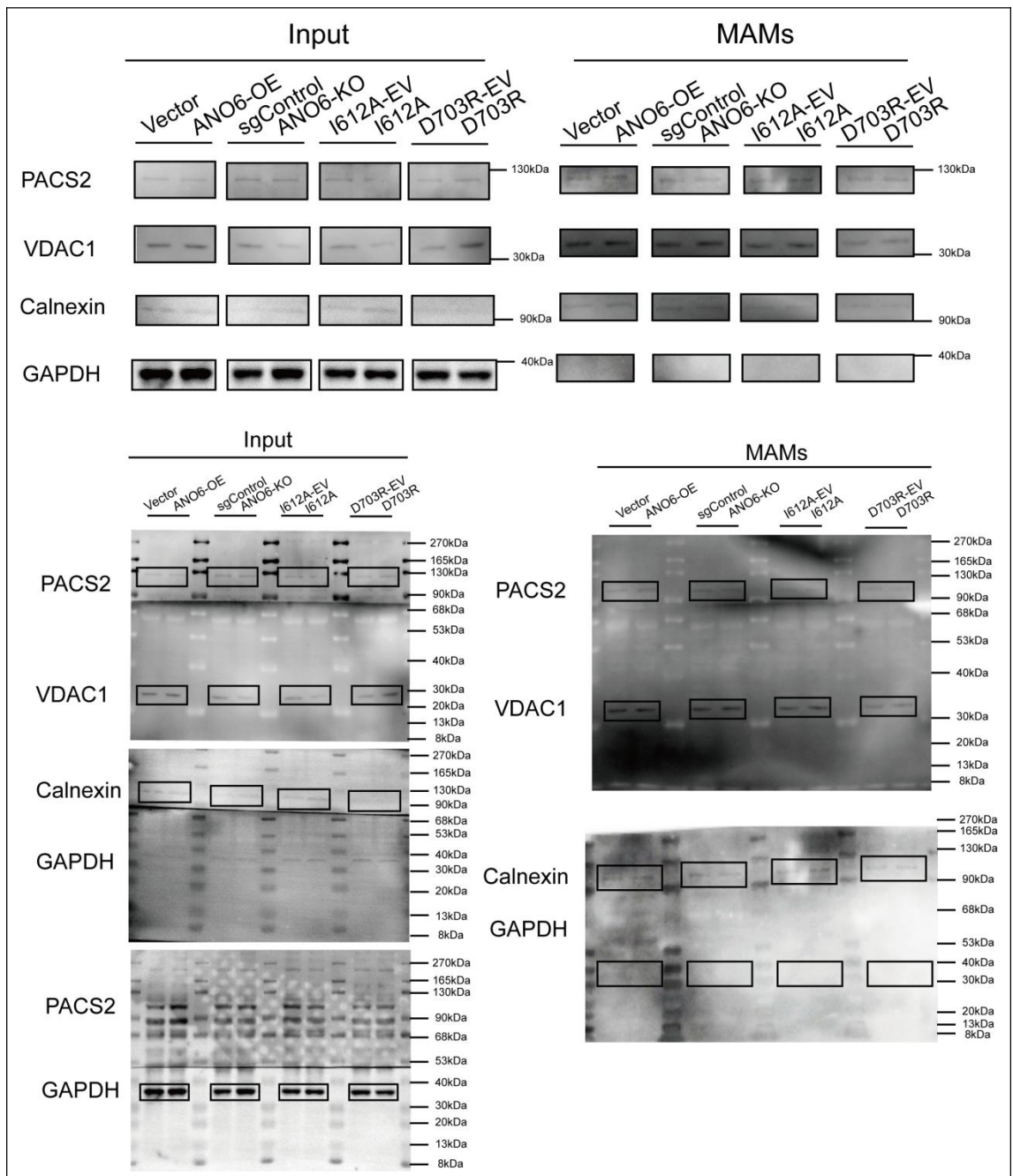

**Supplementary Figure S11. Validation of subcellular fractionation purity for mitochondria-associated endoplasmic reticulum membranes (MAMs).**

Representative immunoblot analyses and full-length uncropped membrane scans verifying the biochemical purity of isolated mitochondria-associated endoplasmic reticulum membrane (MAMs) fractions compared with whole-cell lysates (Input) across engineered EMT6 cell models (Vector, ANO6-OE, sgControl, ANO6-KO, I612A-EV, I612A, D703R-EV, and D703R) under paclitaxel challenge (5 nM PTX, 24 h) corresponding to Main Figure 5: Top panels: Cropped immunoblot bands displaying compartment-specific marker proteins: PACS2 (canonical MAMs marker, 130

kDa), VDAC1 (outer mitochondrial membrane marker, 30 kDa), Calnexin (endoplasmic reticulum marker, 90 kDa), and GAPDH (cytosolic marker, 40 kDa). The isolated MAMs fractions exhibit marked enrichment of MAMs and organellar contact markers (PACS2, VDAC1, and Calnexin) alongside complete absence of cytosolic contamination (GAPDH), confirming the biochemical purity and structural integrity of the harvested contact microdomains utilized for downstream targeted lipidomics. Bottom panels: Full-length uncropped Western blot membranes corresponding to Input (left column) and isolated MAMs fractions (right column) for each indicated protein target. Black rectangular boxes denote the specific regions cropped for the representative panels displayed above. Molecular weight protein ladders (in kDa) are indicated alongside each membrane.

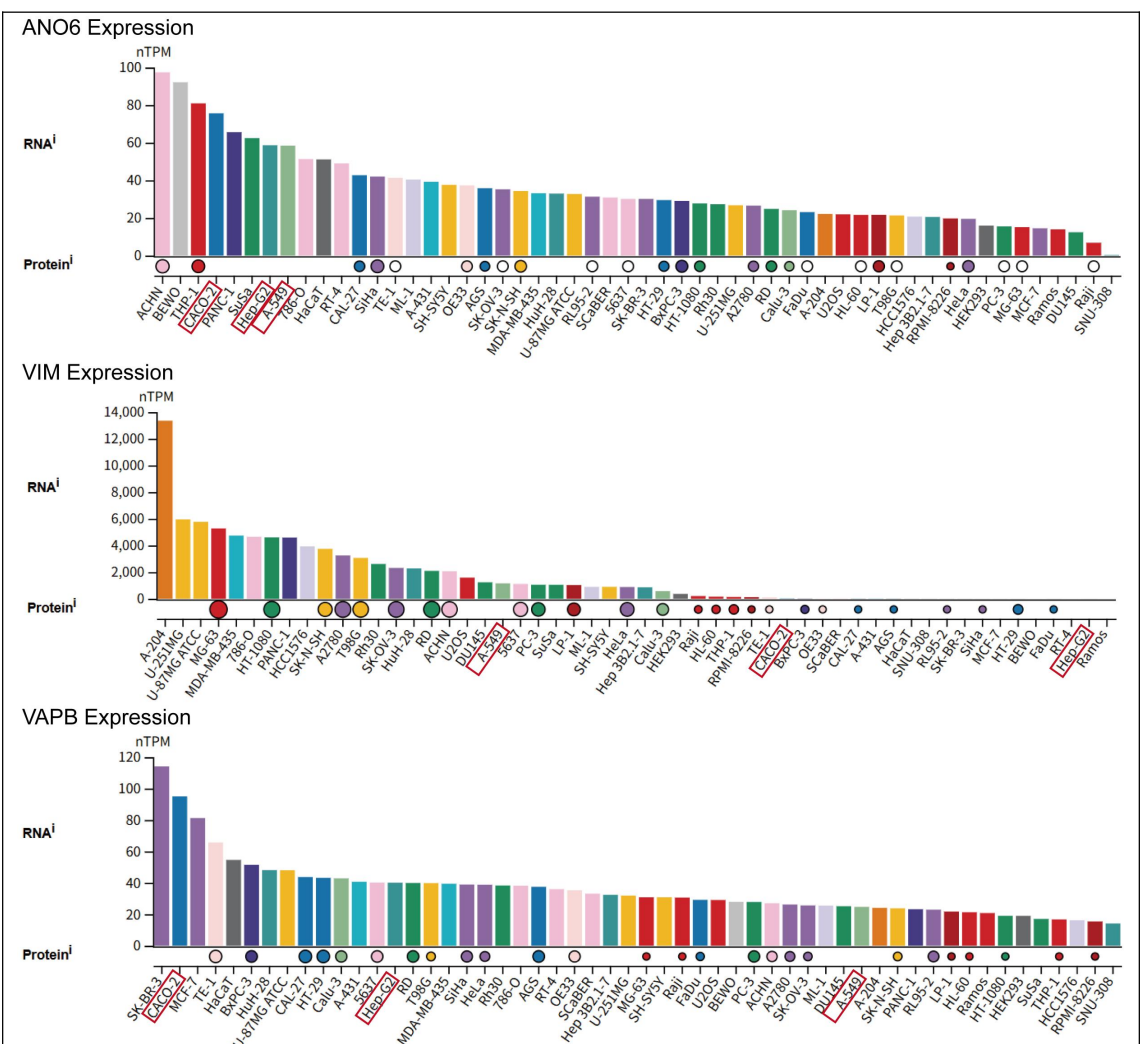

**Supplementary Figure S12. Endogenous expression landscapes of ANO6, VIM, and VAPB across human cancer cell lines in The Human Protein Atlas.**

Transcript (RNA abundance, normalized transcripts per million, nTPM) and corresponding protein expression profiles of ANO6 (top), VIM (middle), and VAPB (bottom) extracted from The Human Protein Atlas (HPA; https://www.proteinatlas.org) across human cancer cell lines, ranked in descending order of expression. The three cell lines selected for cross-cancer validation in Figure 7G—colorectal carcinoma (CACO-2), non-small cell lung carcinoma (A-549), and hepatocellular carcinoma (Hep-G2)—are highlighted with red bounding boxes along the x-axis, showing robust endogenous ANO6 expression alongside baseline abundance of its tethering partners VIM and VAPB.

**Supplementary Table S1.** Baseline Clinicopathological Characteristics of Paired Breast Cancer Patient Cohorts (Total n = 66)

A: Multi-Omics Profiling Cohort (Paired Transcriptomic, Proteomic & Metabolomic Sequencing, n = 6)

| Patient ID | Subtype | Age | Neoadjuvant<br>Chemotherapy<br>Regimen | Pre-NAC<br>Stage<br>(cTNM) | Post-NAC<br>Stage<br>(ypTNM) | Miller-Payne (MP)<br>Grade |
| --- | --- | --- | --- | --- | --- | --- |
| **Seq-01** | TNBC | 53 | TCb * 6 | cT2N2M0 | ypT1N1M0 | MP 3 |
| **Seq-02** | TNBC | 51 | TEC * 6 | cT3N1M0 | ypT4N2M0 | MP 2 |
| **Seq-03** | Luminal B | 41 | EC-T * 8 | cT4N2M0 | ypT4N2M0 | MP 3 |
| **Seq-04** | Luminal B | 53 | TP * 2 -- TCb * 4 | cT2N0M0 | ypT2N0M0 | MP 3 |
| **Seq-05** | Luminal A | 48 | TE * 6 | cT2N1M0 | ypT1N1M0 | MP 3 |
| **Seq-06** | Luminal B | 41 | TE * 6 | cT2N2M0 | ypT1bN2M0 | MP 2 |

B: Luminal Breast Cancer IHC Validation Cohort (n = 35)

| Patient ID | Subtype | Age | Neoadjuvant<br>Chemotherapy<br>Regimen | Pre-NAC<br>Stage<br>(cTNM) | Post-NAC<br>Stage<br>(ypTNM) | Miller-Payne (MP)<br>Grade |
| --- | --- | --- | --- | --- | --- | --- |
| **Lum-01** | Luminal A | 56 | TE * 6 | cT2N1M0 | ypT1N1M0 | MP 4 |
| **Lum-02** | Luminal B | 60 | TEC * 6 | cT4bN3M0 | ypT3N3M0<br>(IIIC) | MP 2 |
| **Lum-03** | Luminal B | 58 | TE * 6 | cT3N2M0 | ypT2N3M0 | MP 3 |
| **Lum-04** | Luminal B | 53 | TE * 6 | cT1N2M0 | ypT1N3M0 | MP 3 |
| **Lum-05** | Luminal A | 34 | TE * 6 | cT2N1M0 | ypT2N1M0 | MP 3 |
| **Lum-06** | Luminal B | 55 | TE * 6 | cT1N1M0 | ypT1N1M0 | MP 3 |
| **Lum-07** | Luminal A | 32 | EC-T * 8 | cT2N2M0 | ypT2N2M0 | MP 4 |
| **Lum-08** | Luminal A | 55 | EC-T * 8 | cT3N1M0 | ypT2N2M0 | MP 3 |
| **Lum-09** | Luminal A | 46 | TE * 6 | cT2N0M0 | ypT1N1M0 | MP 4 |
| **Lum-10** | Luminal B | 39 | TE * 6 | cT2N1M0 | ypT1N1M0 | MP 2 |
| **Lum-11** | Luminal B | 60 | TC * 6 | cT4N2M0 | ypT2N2M0 | MP 2 |
| **Lum-12** | Luminal B | 65 | EC-T | cT3N1M0 | ypT1N1M0 | MP 3 |
| **Lum-13** | Luminal B | 48 | TE-NP * 6 | cT2N1M0 | ypT2N1M0 | MP 2 |
| **Lum-14** | Luminal B | 43 | TE * 6 | cT3N1M0 | ypT1N3M0 | MP 4 |
| **Lum-15** | Luminal B | 58 | EC-T * 8 | cT4aN3M0 | ypT4N3M0 | MP 2 |
| **Lum-16** | Luminal B | 50 | TE * 6 | cT2N1M0 | ypT1N1M0 | MP 2 |
| **Lum-17** | Luminal A | 54 | TE * 6 | cT4bN1M0 | ypT1N1M0 | MP 2 |
| **Lum-18** | Luminal A | 70 | TE * 6 | cT4bN2M0 | ypT4N2M0 | MP 2 |
| **Lum-19** | Luminal B | 35 | TE * 6 | cT2N1M0 | ypT1N0M0 | MP 4 |
| **Lum-20** | Luminal A | 38 | EC-T * 8 | cT3N1M0 | ypT1N0M0 | MP 2 |
| **Lum-21** | Luminal A | 55 | EC-T * 8 | cT2N3M0 | ypT2N3M0 | MP 3 |
| **Lum-22** | Luminal A | 29 | EC-T * 8 | cT3N1M0 | ypT3N2M0 | MP 2 |
| **Lum-23** | Luminal A | 58 | EC-T * 8 | cT4N1M0 | ypT3N0M0 | MP 3 |
| **Lum-24** | Luminal B | 55 | TE * 6 | cT2N1M0 | ypT2N1M0 | MP 3 |
| **Lum-25** | Luminal B | 61 | TC * 6 | cT2N1M0 | ypT1N0M0 | MP 4 |
| **Lum-26** | Luminal B | 55 | TAC * 6 | cT2N1M0 | ypT2N3M0 | MP 3 |
| **Lum-27** | Luminal B | 64 | EC-T * 6 | cT2N1M0 | ypT3N1M0 | MP 3 |
| **Lum-28** | Luminal B | 39 | TE * 6 | cT3N1M0 | ypT2N1M0 | MP 3 |
| **Lum-29** | Luminal A | 37 | EC-T | cT2N1M0 | ypT1N0M0 | MP 3 |
| **Lum-30** | Luminal B | 47 | EC-T | cT4N1M0 | ypT2N0M0 | MP 2 |
| **Lum-31** | Luminal B | 50 | TE * 6 | cT4N1M0 | ypT1N0M0 | MP 3 |
| **Lum-32** | Luminal B | 52 | EC-T | cT3N1M0 | ypT1N0M0 | MP 3 |
| **Lum-33** | Luminal A | 57 | TE * 6 | cT4N1M0 | ypT2N0M0 | MP 3 |
| **Lum-34** | Luminal B | 55 | TE * 6 | cT3N1M0 | ypT1N0M0 | MP 2 |
| **Lum-35** | Luminal B | 54 | EC-T | cT2N1M0 | ypT2N2M0 | MP 1 |

C: Triple-Negative Breast Cancer (TNBC) IHC Validation Cohort (n = 25)

| **Patient ID** | **Subtype** | **Age** | **Neoadjuvant**<br>**Chemotherapy**<br>**Regimen** | **Pre-NAC**<br>**Stage**<br>**(cTNM)** | **Post-NAC**<br>**Stage**<br>**(ypTNM)** | **Miller-Payne (MP)**<br>**Grade** |
| --- | --- | --- | --- | --- | --- | --- |
| **TNBC-01** | TNBC | 52 | TAC * 6 -- TE * 6 | cT3N1M0<br>(IIIA) | ypT1N0M0 | MP 4 |
| **TNBC-02** | TNBC | 45 | EC-T * 6 -- TCb +<br>Toripalimab | cT3N0M0<br>(IIB) | ypT2N0M0<br>(IIA) | MP 2 |
| **TNBC-03** | TNBC | 50 | TAC * 6 | cT1N2M0<br>(IIIA) | ypT2N3M0<br>(IIIC) | MP 2 |
| **TNBC-04** | TNBC | 49 | EC-T | cT3N0M0<br>(IIB) | ypT2N0M0<br>(IIA) | MP 3 |
| **TNBC-05** | TNBC | 49 | EC-TP | cT4N0M0<br>(IIIA) | ypT2N0M0<br>(IIA) | MP 2 |
| **TNBC-06** | TNBC | 53 | TCb + Toripalimab<br>* 6 | cT2N1M0<br>(IIB) | ypT1N0M0 | MP 1 |
| **TNBC-07** | TNBC | 48 | EC-T * 8 | cT4N1M0<br>(IIIB) | ypT1N1M0<br>(IIA) | MP 3 |
| **TNBC-08** | TNBC | 44 | TEC * 6 | cT3N1M0<br>(IIIC) | ypT2N1M0<br>(IIB) | MP 2 |
| **TNBC-09** | TNBC | 54 | EC-T * 8 | cT3N2M0<br>(IIIB) | ypT1bN0M0<br>(IA) | MP 3 |
| **TNBC-10** | TNBC | 49 | TCb * 6 | cT2N3M0<br>(IIIC) | ypT2N3M0<br>(IIIC) | MP 2 |
| **TNBC-11** | TNBC | 54 | EC-T * 8 | cT2N1M0<br>(IIB) | ypT1N1M0<br>(IIA) | MP 4 |
| **TNBC-12** | TNBC | 34 | EC-TCb * 8 | cT2N0M0<br>(IIA) | ypT1N0M0<br>(IA) | MP 3 |
| **TNBC-13** | TNBC | 51 | TE * 6 | cT3N2M0 (IIIA) | ypT2N2M0 (IIIA) | MP 3 |
| **TNBC-14** | TNBC | 66 | TCb * 6 | cT4N1M0 (IIIB) | ypT4N1M0 (IIIB) | MP 2 |
| **TNBC-15** | TNBC | 55 | ddEC-T * 8 | cT2N1M0 (IIB) | ypT1N0M0 (IA) | MP 3 |
| **TNBC-16** | TNBC | 61 | TE * 6 | cT3N1M0 (IIB) | ypT1N2M0 (IIIA) | MP 4 |
| **TNBC-17** | TNBC | 48 | TE * 2 -- TCb * 4 | cT2N0M0 (IIA) | ypT1N0M0 (IA) | MP 1 |
| **TNBC-18** | TNBC | 49 | AC-T + CBP * 8 | cT2N1M0 (IIB) | ypT1N1M0 (IA) | MP 4 |
| **TNBC-19** | TNBC | 56 | TCb * 6 | cT2N3M0 (IIIC) | ypT1N1M0 (IA) | MP 3 |
| **TNBC-20** | TNBC | 61 | EC-T * 8 | cT2N1M0 (IIB) | ypT1N1M0 (IIA) | MP 3 |
| **TNBC-21** | TNBC | 46 | EC-T * 8 | cT2N1M0 (IIB) | ypT1N1M0 (IIA) | MP 2 |
| **TNBC-22** | TNBC | 50 | TEC * 6 | cT1N2M0 (IIIA) | ypT1N3M0 (IIIB) | MP 2 |
| **TNBC-23** | TNBC | 55 | ddEC-T * 8 | cT2N1M0 (IIB) | ypT1N0M0 (IA) | MP 4 |
| **TNBC-24** | TNBC | 56 | TAC * 6 | cT2N0M0 (IIA) | ypT4N1M1 (IV) | MP 2 |
| **TNBC-25** | TNBC | 47 | TCb * 6 | cT3N1M0 (IIIA) | ypT1N0M0 (IA) | MP 2 |

Abbreviations: NAC, Neoadjuvant Chemotherapy; cTNM, clinical TNM stage before neoadjuvant chemotherapy; ypTNM, pathological TNM stage following neoadjuvant chemotherapy and surgery; MP Grade, Miller-Payne histological response grading system (Grade 1: no significant reduction in overall cellularity; Grade 2: minor loss of tumor cells <30%; Grade 3: estimated 30%-90% reduction of tumor cells; Grade 4: marked loss of tumor cells >90%; Grade 5: pathological complete response / pCR); T, Docetaxel / Paclitaxel; E, Epirubicin; C, Cyclophosphamide; A, Doxorubicin (Adriamycin); Cb, Carboplatin; P, Cisplatin; NP, Vinorelbine; dd, dose-dense regimen; Toripalimab, anti-PD-1 monoclonal antibody.

**Supplementary Table S2. Lentiviruses, plasmids, sgRNAs, and primers for mouse and human Ano6 genetic manipulation and Vim functional studies.**

**A. Overexpression lentiviral vectors (Ano6)**

| Resource | Species | Vector Backbone | Selection Marker | Insert / Accession No. |
| --- | --- | --- | --- | --- |
| **LV-Ano6 WT** | Mouse | GV721 | Puromycin | NM_175344.4 (C-terminal 3×FLAG, Luciferase) |
| **CON547** | Mouse | GV721 | Puromycin | Empty vector |
| (Control) |  |  |  |  |
| LV-Ano6 D703R-HA | Mouse | GV870 | Neomycin (G418) | NM_175344.4 (D703R)-Linker-HA (sgRNA-resistant, synthetic) |
| LV-Ano6 I612A-HA | Mouse | GV870 | Neomycin (G418) | NM_175344.4 (I612A)-Linker-HA (sgRNA-resistant, synthetic) |
| CON709 (Control) | Mouse | GV870 | Neomycin (G418) | Empty vector |
| oe-ANO6 | Human | GV492 | Puromycin | NM_001025356.3 (3×FLAG, Luciferase) |
| CON335 (Control) | Human | GV492 | Puromycin | Empty vector |

##### B. CRISPR-Cas9 knockout lentiviral vectors (sgRNAs)

| Resource | Species | Vector Backbone | Selection Marker | Targeting Sequence (5' → 3') |
| --- | --- | --- | --- | --- |
| LV-Ano6-sgRNA-1 (C25J0047) | Mouse | GV708 | Puromycin | TGCTCTCATCTTCATACACG |
| LV-Ano6-sgRNA-2 (C25J0048) | Mouse | GV708 | Puromycin | TGACACTCTCGTTCACCCGG |
| LV-Ano6-sgRNA-3 (C25J0049) | Mouse | GV708 | Puromycin | GAATCTAACCTTATCTGCCA |
| LV-Ano6-scramble (CON544) | Mouse | GV708 | Puromycin | CGCTTCCGCGGCCCGTTCAA |
| LV-ANO6-sgRNA-1 (C26A1538) | Human | GV708 | Puromycin | GCAAGCCACTCCTACAACTG |
| LV-ANO6-sgRNA-2 (C26A1539) | Human | GV708 | Puromycin | GAATCTAACCTTATCTGTC |
| LV-ANO6-sgRNA-3 (C26A1540) | Human | GV708 | Puromycin | TTAATCAAGATAACTGTACA |
| LV-ANO6-scramble (CON544) | Human | GV708 | Puromycin | CGCTTCCGCGGCCCGTTCAA |

##### C. sgRNA cloning oligos and construct sequencing primers

| Oligo / Primer ID | Sequence (5' → 3') | Application |
| --- | --- | --- |
| C25J0047a (F) | acaccgTGCTCTCATCTTCATACACG | Mouse *Ano6* sgRNA-1 forward oligo |
| C25J0047b (R) | aaacCGTGTATGAAGATGAGAGACAc | Mouse *Ano6* sgRNA-1 reverse oligo |
| C25J0048a (F) | acaccgTGACACTCTCGTTCACCCGG | Mouse *Ano6* sgRNA-2 forward oligo |
| C25J0048b (R) | aaacCCGGGTGAACGAGAGGTGTCAc | Mouse *Ano6* sgRNA-2 reverse oligo |
| C25J0049a (F) | acaccgGAATCTAACCTTATCTGCCA | Mouse *Ano6* sgRNA-3 forward oligo |
| C25J0049b (R) | aaacTGGCAGATAAGGTAGATTCC | Mouse *Ano6* sgRNA-3 reverse oligo |
| pGCSIL-F | CCATGATTTCCTTCATATTTGC | Sequencing primer for mouse *Ano6* |
|  |  | sgRNA constructs (GV708) |

##### D. sgRNA-resistant silent mutations introduced in mouse *Ano6* rescue constructs

| sgRNA ID | Original Wild-Type Sequence (5' → 3') | Mutated Silent Sequence (5' → 3') |
| --- | --- | --- |
| C25J0049 | GAATCTAACCTTATCTGCCA | GAGTCCAATCTCATTTGTCA |

##### E. Sequencing and validation primers for mouse *Ano6* rescue constructs (GV870)

| Primer ID | Sequence (5' → 3') | Application |
| --- | --- | --- |
| CMV-F | CGCAATGGGCGGTAGGCGTG | Vector backbone sequencing |
| K26J2192-wf1 | gcacatcaaactcccgctaa | *Ano6* D703R-HA / I612A-HA internal sequencing |
| K26J2192-wf2 | cggtggccagatcctgatgt | *Ano6* D703R-HA / I612A-HA internal sequencing |
| K26J2192-wf3 | gacccagacggattatgaga | *Ano6* D703R-HA / I612A-HA internal sequencing |
| K26J2192-wf4 | gatcatttgctttcacatcag | *Ano6* D703R-HA / I612A-HA internal sequencing |
| EF1-R | ACATCACTTTCCCAGTTTACC | Vector backbone sequencing |

##### F. RNAi knockdown lentiviral vectors (mouse *Vim* shRNAs)

| Resource | Species | Vector Backbone | Selection Marker | Target Sequence (5' → 3') |
| --- | --- | --- | --- | --- |
| LV-*Vim*-shRNA-1 (P26F1261) | Mouse | GV116 | Hygromycin | GTGGAATCCTTGCAAGGAAGAA |
| LV-*Vim*-shRNA-2 (P26F1262) | Mouse | GV116 | Hygromycin | CGACGCCATCAACACTGAGTT |
| LV-*Vim*-shRNA-3 (P26F1263) | Mouse | GV116 | Hygromycin | GCATCACGATGACCTTGAATA |
| LV-*Vim*-shRNA-NC (CON482) | Mouse | GV116 | Hygromycin | TTCTCCGAACGTGTCACGT |

##### G. Expression plasmids for Co-IP and domain mapping

| Resource | Species | Vector Backbone | Selection Marker | Insert Description |
| --- | --- | --- | --- | --- |
| *Vim*-HA (Full-length) | Mouse | GV366 | Neomycin | NM_011701.4 (Full-length coding sequence) |
| *Vim* del(103-411aa)-HA | Mouse | GV366 | Neomycin | NM_011701.4 lacking the central rod domain (residues 103-411) |
| CON237 (Control) | Mouse | GV366 | Neomycin | Empty vector control |

##### H. Vector features and experimental descriptions

GV708: U6-sgRNA-EF1α-Cas9-FLAG-CMV-P2A-puro (cloning site: BsmBI). Used for generating CRISPR-Cas9-mediated knockout lines in mouse and human cells. Transduced cells were selected with puromycin, followed by single-cell cloning and Western blot validation.

GV721: CMV enhancer-MCS-3FLAG-EF1α-firefly\_Luciferase-SV40-Puromycin (cloning site: BamHI/BamHI). Used for generating mouse Ano6 WT overexpression lines.

GV870: CMV enhancer-MCS-hEF1/HTLVp-firefly\_Luciferase-T2A-Neomycin (cloning site: XbaI/BamHI). Used for generating mouse Ano6 D703R-HA and I612A-HA rescue lines. The empty vector control is CON709.

GV492: Ubc-MCS-3FLAG-CBh-luciferase-IRES-puromycin (cloning site: BamHI/AgeI). Used for establishing stable human ANO6 overexpression lines.

GV116: hU6-MCS-CMV-Hygromycin (cloning site: AgeI/BamHI). Used for stable RNAi knockdown of mouse Vimentin.

GV366: CMV-MCS-HA-SV40-Neomycin (cloning site: XhoI/BamHI). Used for transient expression and domain-mapping Co-IP assays (wild-type Vimentin vs. rod domain deletion mutant). Design notes for rescue cDNAs: Mouse D703R-HA and I612A-HA cDNAs were generated via de novo gene synthesis. To avoid cleavage by the active sgRNA (C25J0049), seven silent mutations were introduced into the corresponding target sequence without altering the amino acid sequence (GAATCTAACCTTATCTGCCA→GAGTCCAATCTCATTTGTCA). All constructs and oligos were verified by Sanger sequencing and obtained from GeneChem (Shanghai, China).

##### Supplementary Sequences

###### 1. Synthesized mouse D703R-HA sgRNA-resistant Ano6 cDNA (GV870, XbaI-BamHI)

>D703R-HA\_sgRNA-resistant\_mouse\_Ano6\_GV870

```
TCCGGTGAATTCCTCGAGACTAGTTCTAGACGCCACCATGCAGATGATGACTAGGAAGGTCCTGCTGAACAT
GGAGCTGGAGGAGGACGACGATGAGGATGGAGACATTGTGCTGGAAAACTTTGACCAGACAATTGTCTGC
CCCACCTTTGGATCACTGGAGAATCAGCAGGACTTCAGGACTCCAGAGTTTGAAGAATTTAACGGGAAGCC
CGACTCCCTCTTTTTCACCGATGGCCAGAGGCGAATCGACTTCATCCTCGTGTATGAAGATGAGAGCAAAAA
GGAGAACAATAAGAAAGGGACAAATGAGAAACAGAAGAGGAAAAGACAAGCATACGAGTCCAATCTCATT
TGTCATGGGCTGCAGCTGGAAGCAACAAGATCTGTTTCTGATGACAAGCTTGTGTTCGTAAAAGTGCACGC
GCCCTGGGAAGTGCTGTGCACCTATGCTGAGATCATGCACATCAAACTCCCGCTAAAGCCAAACGACCTGAA
AACGCGCTCGCCCTTTGGCAACCTCAACTGGTTCACCAAGGTCCTCCGGGTGAACGAGAGTGTCATCAAGC
CAGAGCAGGAGTTCTTCACTGCCCCTTTTGAGAAGAGCCGGATGAATGATTTCTACATCCTCGATAGAGATT
CCTTCTTCAACCCTGCCACCAGAAGCCGCATTGTTTATTTCATCCTCTCTCGGGTCAAATACCAAGTGATGAA
CAACGTTAACAAATTTGGGATTAATAGACTGGTCAGCTCTGGAATCTACAAAGCAGCGTTTCCTCTGCACGA
CTGCAGATTCAACTATGAGTCGGAGGACATCAGTTGTCCTAGCGAGCGTTACCTCCTGTACAGAGAATGGGC
TCACCCTCGGAGTATATACAAGAAGCAGCCCTTGGATCTTATCAGGAAGTATTACGGCGAGAAGATTGGAAT
CTACTTTGCTTGGCTGGGCTATTACACGCAGATGCTCCTTCTCGCAGCTGTGGTGGGCGTGGCCTGCTTCCTC
TATGGATATCTTGATCAGGATAACTGCACTTGGAGCAAAGAGGTCTGTGATCCTGACATCGGTGGCCAGATC
CTGATGTGCCCCCAGTGTGACAGGCTGTGTCCGTTCTGGAGGCTGAATATCACCTGTGAATCTTCTAAGAAA
TTGTGCATCTTTGACAGTTTTGGAACCCTGATCTTCGCTGTATTTATGGGAGTGTGGGTTACATTGTTTTTGG
AGTTTTGGAAGCGGCGCCAGGCAGAGCTCGAATATGAATGGGACACCGTTGAGCTACAGCAAGAGGAACA
GGCCCGGCCAGAGTATGAGGCCCAGTGCAATCACGTGGTGATAAACGAGATCACTCAGGAGGAGGAGCGC
ATCCCATTTACCACCTGTGGGAAGTGCATCCGGGTCACGCTGTGTGCGAGCGCCGTCTTCTTCTGGATCCTG
CTCATCATCGCCTCTGTGATCGGGATCATTGTCTACAGGCTGTCCGTGTTCATCGTATTTTCCACAACGCTCCC
CAAGAACCCCAACGGGACAGACCCGATCCAGAAGTACCTGACCCCACAGATGGCCACATCCATCACAGCCT
CCATCATCAGCTTCATCATCATCATGATCCTCAACACGATCTACGAGAAGGTGGCCATCATGATCACCAACTTC
GAGCTCCCAAGGACCCAGACGGATTATGAGAACAGCCTGACCATGAAGATGTTCTTGTTCCAGTTTGTCAAC
```

TATTACTCCTCATGCTTCTACATCGCATTCTTCAAGGGCAAATTCGTGGGCTATCCCGGGGACCCAGTGTACTT
GCTGGGCAAATACAGAAGCGAAGAGTGTGACCCGGGGGGCTGCCTCCTTGAACTGACCACACAGCTGACG
ATCATCATGGGGGGAAAGGCAATCTGGAACAACATACAAGAAGTCTTGCTCCCATGGGTTATGAATCTAATT
GGACGATATAAAAGAGTCTCGGGATCAGAAAAGATAACCCCACGATGGGAACAGGATTACCATCTGCAGCC
CATGGGCAAGCTGGGATTGTTCTATGAGTATCTTGAAATGATTATTCAGTTTGGGTTCGTCACTTTATTTGTGG
CCTCTTTCCCACTGGCTCCCCTGCTGGCTCTGGTGAACAATATTTTGGAAATAAGAGTGAGAGCGTGGAAGC
TCACAACCCAGTTTAGGCGCATGGTGCCCGAAAAAGCCCAGGATATCGGCGCGTGGCAGCCCATCATGCAA
GGAATAGCCATCCTGGCCGTGGTGACCAACGCCATGATCATTGCTTTCACATCAGACATGATCCCCCGCCTGG
TGTATTATTGGTCTTTCTCTATCCCTCCCTACGGGGACCACACTTATTACACCATGGACGGCTACATCAACAAC
ACTCTCTCTGTCTTCAACATCACGGACTTCAAGAACACAGACAAAGAAAACCCATACATTGGGCTTGGTAAC
TATACCTTGTGCAGGTACCGTGACTTCCGAAACCCACCTGGGCACCCACAAGAGTATAAACACAACATCTACT
ATTGGCACGTGATTGCCGCCAAGCTGGCCTTCATCATTGTCATGGAGCACATCATCTACTCTGTGAAGTTCTT
CATTTCCTACGCAATTCCAGATGTGTCCAAAATCACGAAAAGCAAGATCAAGAGGGAGAAATACCTCACGCA
GAAGTTGCTTCACGAGAGTCACCTCAAAGACCTGACCAAAAACATGGGGATCATCGCCGAGAGGATAGGA
GGAACGGTTGACAACAGCGTGCGGCCAAAACTCGAAGGAGGTGGAGGATCAGGGGGTGGGGGATCCGG
CGGTGGCGGATCTTACCCTTATGATGTCCCAGACTATGCTTGAGGATCCAGATCCATCGATACTAGTAAGGAT

### 2. Synthesized mouse I612A-HA sgRNA-resistant Ano6 cDNA (GV870, XbaI-BamHI)

>I612A-HA\_sgRNA-resistant\_mouse\_Ano6\_GV870

TCTAGACGCCACCATGCAGATGATGACTAGGAAGGTCCTGCTGAACATGGAGCTGGAGGAGGACGACGAT
GAGGATGGAGACATTGTGCTGGAAAACTTTGACCAGACAATTGTCTGCCCCACCTTTGGATCACTGGAGAA
TCAGCAGGACTTCAGGACTCCAGAGTTTGAAGAATTTAACGGGAAGCCCGACTCCCTCTTTTTCACCGATG
GCCAGAGGCGAATCGACTTCATCCTCGTGTATGAAGATGAGAGCAAAAAGGAGAACAATAAGAAAGGGAC
AAATGAGAAACAGAAGAGGAAAAGACAAGCATACGAGTCCAATCTCATTTGTCATGGGCTGCAGCTGGAA
GCAACAAGATCTGTTTCTGATGACAAGCTTGTGTTCGTAAAAGTGCACGCGCCCTGGGAAGTGCTGTGCAC
CTATGCTGAGATCATGCACATCAAACTCCCGCTAAAGCCAAACGACCTGAAAACGCGCTCGCCCTTTGGCAA
CCTCAACTGGTTCACCAAGGTCCTCCGGGTGAACGAGAGTGTCATCAAGCCAGAGCAGGAGTTCTTCACTG
CCCCTTTTGAGAAGAGCCGGATGAATGATTTCTACATCCTCGATAGAGATTCCTTCTTCAACCCTGCCACCAG
AAGCCGCATTGTTTATTTCATCCTCTCTCGGGTCAAATACCAAGTGATGAACAACGTTAACAAATTTGGGATT
AATAGACTGGTCAGCTCTGGAATCTACAAAGCAGCGTTTCCTCTGCACGACTGCAGATTCAACTATGAGTCG
GAGGACATCAGTTGTCCTAGCGAGCGTTACCTCCTGTACAGAGAATGGGCTCACCCTCGGAGTATATACAAG
AAGCAGCCCTTGGATCTTATCAGGAAGTATTACGGCGAGAAGATTGGAATCTACTTTGCTTGGCTGGGCTAT
TACACGCAGATGCTCCTTCTCGCAGCTGTGGTGGGCGTGGCCTGCTTCCTCTATGGATATCTTGATCAGGATA
ACTGCACTTGGAGCAAAGAGGTCTGTGATCCTGACATCGGTGGCCAGATCCTGATGTGCCCCCAGTGTGAC
AGGCTGTGTCCGTTCTGGAGGCTGAATATCACCTGTGAATCTTCTAAGAAATTGTGCATCTTTGACAGTTTTG
GAACCCTGATCTTCGCTGTATTTATGGGAGTGTGGGTTACATTGTTTTTGGAGTTTTGGAAGCGGCGCCAGG
CAGAGCTCGAATATGAATGGGACACCGTTGAGCTACAGCAAGAGGAACAGGCCCGGCCAGAGTATGAGGC
CCAGTGCAATCACGTGGTGATAAACGAGATCACTCAGGAGGAGGAGCGCATCCCATTTACCACCTGTGGGA
AGTGCATCCGGGTCACGCTGTGTGCGAGCGCCGTCTTCTTCTGGATCCTGCTCATCATCGCCTCTGTGATCGG
GATCATTGTCTACAGGCTGTCCGTGTTCATCGTATTTTCCACAACGCTCCCCAAGAACCCCAACGGGACAGA
CCCGATCCAGAAGTACCTGACCCCACAGATGGCCACATCCATCACAGCCTCCATCATCAGCTTCATCATCATCA
TGATCCTCAACACGATCTACGAGAAGGTGGCCATCATGATCACCAACTTCGAGCTCCCAAGGACCCAGACG
GATTATGAGAACAGCCTGACCATGAAGATGTTCTTGTTCCAGTTTGTCAACTATTACTCCTCATGCTTCTACAT
CGCATTCTTCAAGGGCAAATTCGTGGGCTATCCCGGGGACCCAGTGTACTTGCTGGGCAAATACAGAAGCG

AAGAGTGTGACCCGGGGGGCTGCCTCCTTGAACTGACCACACAGCTGACGATCGCCATGGGGGGAAAGG
CAATCTGGAACAACATACAAGAAGTCTTGCTCCCATGGGTTATGAATCTAATTGGACGATATAAAAGAGTCTC
GGGATCAGAAAAGATAACCCCACGATGGGAACAGGATTACCATCTGCAGCCCATGGGCAAGCTGGGATTGT
TCTATGAGTATCTTGAAATGATTATTCAGTTTGGGTTCGTCACTTTATTTGTGGCCTCTTTCCCACTGGCTCCCC
TGCTGGCTCTGGTGAACAATATTTTGGAAATAAGAGTGGACGCGTGGAAGCTCACAACCCAGTTTAGGCGC
ATGGTGCCCGAAAAAGCCCAGGATATCGGCGCGTGGCAGCCCATCATGCAAGGAATAGCCATCCTGGCCGT
GGTGACCAACGCCATGATCATTGCTTTCACATCAGACATGATCCCCCGCCTGGTGTATTATTGGTCTTTCTCTA
TCCCTCCCTACGGGGACCACACTTATTACACCATGGACGGCTACATCAACAACACTCTCTCTGTCTTCAACATC
ACGGACTTCAAGAACACAGACAAAGAAAACCCATACATTGGGCTTGGTAACTATACCTTGTGCAGGTACCGT
GACTTCCGAAACCCACCTGGGCACCCACAAGAGTATAAACACAACATCTACTATTGGCACGTGATTGCCGCC
AAGCTGGCCTTCATCATTGTCATGGAGCACATCATCTACTCTGTGAAGTTCTTCATTTCCTACGCAATTCCAGA
TGTGTCCAAAATCACGAAAAGCAAGATCAAGAGGGAGAAATACCTCACGCAGAAGTTGCTTCACGAGAGT
CACCTCAAAGACCTGACCAAAAACATGGGGATCATCGCCGAGAGGATAGGAGGAACGGTTGACAACAGCG
TGCGGCCAAAACTCGAAGGAGGTGGAGGATCAGGGGGGATCAGGCGGTGGCGGATCTTACCCTTATGATG
TCCCAGACTATGCTTGAGGATCC
